# γ-Protocadherins control synapse formation and peripheral branching of touch sensory neurons

**DOI:** 10.1101/2022.05.25.493080

**Authors:** Shan Meltzer, Katelyn Comeau, Anda Chirila, Emmanuella Osei-Asante, Michelle DeLisle, Qiyu Zhang, Brian T. Kalish, Aniqa Tasnim, Erica Huey, Leah C. Fuller, Erin K. Flaherty, Julie L. Lefebvre, Tom Maniatis, Andrew M. Garrett, Joshua A. Weiner, David D. Ginty

## Abstract

Light touch sensation begins with activation of low-threshold mechanoreceptor (LTMR) endings in the skin and propagation of their signals to the spinal cord and brainstem. We found that the clustered protocadherin gamma (*Pcdhg*) gene locus, which encodes 22 cell-surface homophilic binding proteins, is required in somatosensory neurons for normal behavioral reactivity to a range of tactile stimuli. Developmentally, distinct Pcdhg isoforms mediate LTMR synapse formation through neuron-neuron interactions and peripheral axonal branching through neuron-glia interactions. The Pcdhgc3 isoform mediates homophilic interactions between sensory axons and spinal cord neurons to promote synapse formation *in vivo*, and is sufficient to induce postsynaptic specializations *in vitro*. Moreover, loss of Pcdhgs and somatosensory synaptic inputs to the dorsal horn lead to fewer corticospinal synapses onto dorsal horn neurons. These findings reveal essential roles for Pcdhg isoform diversity in somatosensory neuron synapse formation, peripheral axonal branching, and step-wise assembly of central mechanosensory circuitry.

## Introduction

The skin, our largest organ, mediates our sense of touch, which is essential for fundamental tasks ranging from object recognition to social exchange. The sense of touch also contributes to brain development, such that deprivation or over-reactivity to touch during early stages of animal development can result in aberrant brain development and function (Hertenstein et al., 2006; Main and Stadtman, 1981; Orefice, 2020). Light touch sensation begins with activation of low-threshold mechanoreceptor (LTMR) axonal endings in the skin, and the relay of LTMR signals to the spinal cord and brainstem. In the skin, LTMR subtypes form highly diverse and intricate end organ structures, which together with their intrinsic physiological properties underlie their unique responses to tactile stimuli (Handler and Ginty, 2021). LTMRs extend central axons to the LTMR-recipient zone (LTMR-RZ) of the spinal cord dorsal horn and to the brainstem, where they form synapses with second-order neurons that process tactile information (Abraira and Ginty, 2013; Abraira et al., 2017; Olausson et al., 2002). Collectively, the exquisitely patterned LTMR peripheral endings and their central synaptic connections enable detection and central representations of myriad tactile stimuli. Yet, of all the senses, touch remains one of the least understood at the developmental level (Jenkins and Lumpkin, 2017; Meltzer et al., 2021; Olson et al., 2016), and the molecular mechanisms that orchestrate LTMR peripheral branching and central synaptic connectivity are poorly understood.

Based on their action potential conduction velocity, cutaneous LTMRs are classified as Aβ-, Aδ-, or C-LTMRs (Horch et al., 1977). LTMRs are further distinguished by the cutaneous end organs with which they associate, rates of adaptation to indentation of the skin (rapidly adapting (RA), intermediate adapting (IA), or slowly adapting (SA)), and preferred stimuli (Handler and Ginty, 2021). In mouse hairy skin, guard hair follicles account for ∼1% of the trunk hair follicles and are associated with Aβ RA-LTMRs, Aβ SAI-LTMRs, and Aβ field-LTMRs, which are differentially sensitive to hair deflection, skin indentation, and stroking (Abraira and Ginty, 2013; Burgess et al., 1968; Rutlin et al., 2014). Awl/auchene and zigzag hair follicles, which comprise ∼99% of trunk hair follicles, are innervated by Aβ RA-LTMRs (Awl/auchene hair only), Aβ field-LTMRs, Aδ-LTMRs, and C-LTMRs (Bai et al., 2015; Li et al., 2011). In the glabrous skin, LTMRs form a variety of end organ types associated with Aβ RA-LTMRs and Aβ SA-LTMRs (Johnson, 1983; Zimmerman et al., 2014). Individual LTMR subtypes form a range of branch numbers and patterns, which establishes their physiological receptive fields (Handler and Ginty, 2021; Jenkins and Lumpkin, 2017; Meltzer et al., 2021). For example, a single Aβ field-LTMR neuron branches extensively (20-180 branches), with each branch forming a circumferential ending around a different hair follicle (Bai et al., 2015). Similarly, in glabrous skin, individual Aβ RA-LTMRs branch multiple times (average ∼7 branches), and each branch typically innervates a single Meissner corpuscle (Neubarth et al., 2020). How LTMR branching patterns are controlled by extrinsic cues and how interactions between sensory neurons and surrounding non-neuronal cells promote branching across the different LTMR subtypes remains unknown.

In the spinal cord dorsal horn, LTMR terminals are organized in a somatotopic and highly overlapping manner in LTMR-RZ, which spans lamina II_iv_ through V (Abraira et al., 2017). Most LTMR-RZ interneurons receive convergent histologically-defined synaptic inputs from two or more LTMR subtypes and, underscoring the essential role of this spinal cord region, LTMR-RZ interneurons regulate tactile reactivity and behavior. Like all dorsal root ganglia (DRG) neurons, LTMRs are glutamatergic and provide the majority of glutamatergic synaptic inputs into the LTMR-RZ. Locally projecting interneurons account for the majority of neurons in the LTMR-RZ, and these neurons are highly interconnected, whereas projection neurons account for a small subset of LTMR-RZ neurons (Abraira et al., 2017). During mouse embryonic development beginning around E14, somatosensory neuron axons penetrate the spinal cord in sequence, with axons of large diameter neurons entering first followed by small diameter neuron axons, and these axons then establish synaptic connections with spinal cord neurons over the next several days to weeks (Mirnics and Koerber, 1995; Ozaki and Snider, 1997). Mechanical stimuli evoked behavioral responses and physiological responses recorded in the dorsal horn are first observed at late gestational ages (Andrews and Fitzgerald, 1994; Fitzgerald, 1985). The precise timing of the formation of synapses between LTMR subtypes and spinal cord neurons, the molecular signals that govern this process, and how LTMR inputs may affect the functional organization of other inputs to the dorsal horn remain poorly understood.

The clustered protocadherins encode a family of diverse cell-surface homophilic proteins and are tandemly arrayed in the genome in α-, β- and γ- subclusters, called *Pcdha*, *Pcdhb* and *Pcdhg*, respectively (Wu and Maniatis, 1999). In the *Pcdha* and *Pcdhg* subclusters, variable exons encoding extracellular, transmembrane, and juxtamembrane domains are spliced to three constant exons (Obata et al., 1998; Sugino et al., 2000; Tasic et al., 2002; Wang et al., 2002a). Thus, each *Pcdhg* gene encodes a protein with a unique extracellular and transmembrane domain, a variable cytoplasmic domain, followed by a shared C-terminal domain (Flaherty and Maniatis, 2020; Schreiner and Weiner, 2010; Thu et al., 2014; Wu and Maniatis, 1999). The clustered Pcdhs play critical roles in many aspects of nervous system development, including dendritic self-avoidance, dendrite arborization, neuronal survival, astrocyte-neuron interactions, and axonal tiling (Chen et al., 2012; 2017; Garrett et al., 2019; Hasegawa et al., 2016; Lefebvre et al., 2012; 2008; Mountoufaris et al., 2018; Prasad and Weiner, 2011). Although Pcdhgs can be found at synapses, especially in postsynaptic density (PSD) fractions of purified synaptosomes (Loh et al., 2016; Phillips et al., 2003; Wang et al., 2002b; Weiner et al., 2005), and they form two-dimensional structures between adjacent membranes (Brasch et al., 2019; Goodman et al., 2017; 2016a; 2016b; Nicoludis et al., 2016; Rubinstein et al., 2015), functional evidence demonstrating a direct role in synapse formation has remained elusive (Peek et al., 2017; Südhof, 2018).

Here, we determined the timing of synapse formation between LTMRs and spinal cord neurons of the LTMR-RZ and then assessed LTMR gene expression profiles during the peak of postnatal synapse formation to identify candidate synaptogenic regulators. This analysis drew attention to the clustered protocadherin gamma (*Pcdhg*) genes, and genetic perturbation experiments revealed that they have crucial roles in somatosensory behaviors, sensory neuron synapse formation, and LTMR peripheral axon branching. Pcdhgc3 is the principal isoform for somatosensory neuron synapse formation; it is necessary and sufficient to promote synapse assembly through homophilic interactions between sensory and spinal cord neurons *in vivo*, and is sufficient to promote postsynaptic specialization in a reduced *in vitro* preparation. Pcdhgs also promote peripheral LTMR axonal branching in the skin by mediating interactions with peripheral glial cells, a function that requires family members other than Pcdhgc3. Interestingly, the reduction of mechanosensory inputs and synaptic drive in the LTMR-RZ of sensory neuron *Pcdhg* mutants diminished spinal cord responses to touch stimuli and reduced corticospinal inputs to the dorsal horn. These observations reveal unique functions for distinct clustered protocadherin family members in somatosensory neuron synapse formation and peripheral axonal branching, define an essential role for Pcdhgc3 homophilic interactions in synapse formation, and demonstrate that sensory inputs to the spinal cord indirectly control the assembly of other components of spinal cord circuitry.

## Results

### *Pcdhgs* are expressed in the postnatal DRG and spinal cord when sensory neurons are forming synaptic connections

To define the postnatal period of synapse formation between mechanosensory neurons and spinal cord neurons, we used previously characterized genetic tools to label synapses of all primary somatosensory neuron subtypes (*Advillin^Cre^*) (Hasegawa et al., 2007) as well as synapses of two representative LTMR subtypes, Aδ-LTMRs (*TrkB^CreER^*) (Rutlin et al., 2014) and Aβ RA-LTMRs (*Ret^CreER^*) (Luo et al., 2009). We used a Cre-dependent synaptophysin-tdTomato reporter line (Ai34) to visualize presynaptic terminals of these sensory neurons during postnatal development (P0, P3, P10, and P21) with a focus on synapses in the thoracic LTMR-RZ. The size of presynaptic sensory terminals increased between P0 and P21 for all three groups of sensory neurons (Figures 1A and 1B). To visualize postsynaptic specialization of these sensory neuron glutamatergic synapses, we stained for the postsynaptic scaffolding protein Homer1, which is present at postsynaptic densities (PSDs) of glutamatergic synapses within the dorsal horn (Gutierrez-Mecinas et al., 2016). Homer1 proteins localize to PSDs distal to the synaptic cleft compared to other excitatory PSD markers (Dani et al., 2010), allowing reliable labeling and quantification of their density (Abraira et al., 2017; Gutierrez-Mecinas et al., 2016). The number and size of Homer1 puncta associated with sensory neuron terminals increased more than 2-fold between P0 and P21 (Figures 1C and S1A). Thus, somatosensory neuron synapses in the dorsal horn are formed throughout early postnatal development (Figure 1D).

**Figure 1.**
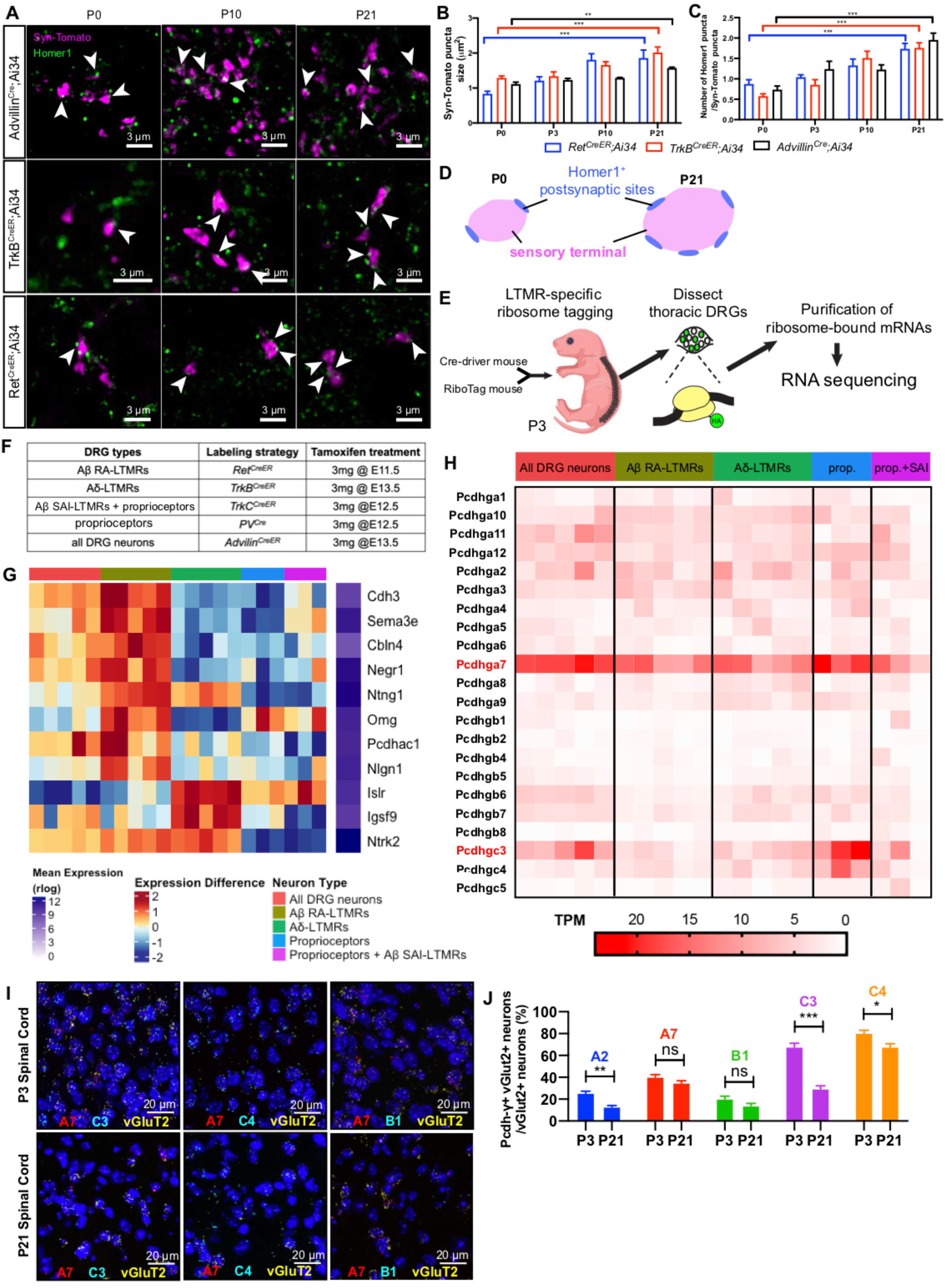
*Pcdhgs* are expressed during postnatal development when somatosensory axons actively form synapses in the dorsal horn. **(A)** IHC images of spinal cord dorsal horn lamina III from P0, P10 and P21 wildtype mice expressing synaptophysin-tdTomato (Ai34) driven by *Advillin^Cre^*, *TrkB^CreER^*, and *Ret^CreER^* to label all sensory neurons, Aδ-LTMRs, and Aβ RA-LTMRs, respectively. Arrowheads point to some of the synapses made between sensory neuron terminals and spinal cord neurons. Scale bars represent 3 μm. **(B)** Quantifications of the sizes of sensory terminals labeled by synaptophysin-tdTomato at P0, P3, P10 and P21. *Ret^CreER^;Ai34*: n = 2 animals for each time point. *TrkB^CreER^;Ai34*: n = 4 animals for P0, 2 animals for P3, 3 animals for P10, and 3 animals for P21. *Advillin^Cre^;Ai34*: n = 3 animals for P0, 2 animals for P3, 2 animals for P10, and 4 animals for P21. Two-way ANOVA. **(C)** Quantifications of the average numbers of Homer1 puncta per sensory terminal labeled by synaptophysin-tdTomato at P0, P3, P10 and P21. Two-way ANOVA. **(D)** Summary of the synapse formation surrounding sensory terminals in the LTMR recipient zone during postnatal development, showing that sensory terminals become larger and more postsynaptic sites form between the sensory terminals and spinal cord neurons. **(E)** Schematic of the RNA sequencing workflow using immunoprecipitation of ribosome-associated mRNA transcripts. **(F)** Genetic labeling strategies for each of DRG neuron groups. **(G)** Heatmaps depicting expression patterns for differentially expressed genes encoding cell adhesion molecules or axon guidance proteins. Each column is one biological replicate, and each row shows the expression level for one gene. **(H)** Heatmap depicting expression patterns of *Pcdhg* genes in the DRGs at P3. Expression levels are represented as TPM values. **(I and J)** RNAscope images (I) and quantification (J) for *Pcdhga2*, *Pcdhga7*, *Pcdhgb1*, *Pcdhgc3*, and *Pcdhgc4* expression levels (n = 3 animals for each time point). SLC17A6(vGluT2) is labeled to visualize the excitatory neurons in the dorsal spinal cord. DAPI staining in blue labels cell nuclei. Student’s unpaired t test. ns, not significant; *p < 0.05; **p < 0.01; ***p < 0.001.

To explore the molecular basis of LTMR synapse formation and other developmental processes, including peripheral target field innervation and end organ formation, across LTMR subtypes, we next performed deep RNA sequencing of major LTMR subtypes, proprioceptors, as well as all DRG neurons to obtain and compare their transcriptome profiles at a relevant developmental stage. P3 was chosen because LTMRs exhibited both central synapse formation (Figures 1A-1C) and peripheral target innervation during this early postnatal age (Meltzer et al., 2021; Rutlin et al., 2014). For this analysis, we used genetic tools that selectively label subsets of LTMRs and proprioceptors in combination with mice that express the epitope-tagged ribosomal protein RPL22^HA^ (RiboTag mice), enabling isolation of mRNA transcripts in these populations using immunoprecipitated polyribosomes (Figures 1E and 1F) (Sanz et al., 2009). RNA libraries were prepared from acutely isolated DRGs and paired-end sequenced using an Illumina NextSeq 500 platform at an average depth of ∼13,000 genes per sample. This sensitive approach allowed for the detection of lowly-expressed genes in Aβ RA-LTMRs, Aβ SAI-LTMRs, Aδ-LTMRs, and proprioceptors. Principal component analysis (PCA) showed that biological replicates for the same neuronal subtype cluster with each other, suggesting that differences in gene expression observed across groups are not caused by batch effects (Figure S1B).

To assess the validity of the transcriptome profiles, we examined expression patterns of several marker genes, including genes whose expression patterns were previously identified in prior DRG sequencing studies, subtype-uniquely enriched genes (SUEGs), and differentially expressed transmembrane proteins. While expression patterns of several genes in developing LTMRs were also detected in mature LTMRs, including *Ntrk2*, *Calb2*, *Adtrp* and *Cdh3* (Figures 1G) (Sharma et al., 2020; Usoskin et al., 2015; Zheng et al., 2019), many SUEGs found in P3 Aβ RA-LTMRs, Aβ SAI-LTMRs, and Aδ-LTMRs were identified here for the first time (Figures S1C and S1D). We hypothesized that differentially expressed transmembrane molecules in developing LTMRs may regulate their synapse formation and peripheral innervation. Thus, we examined expression patterns of cell adhesion molecules and axon guidance proteins previously implicated in neural circuit assembly (Südhof, 2012; 2018; Togashi et al., 2009). This analysis revealed several candidate differentially expressed genes encoding transmembrane proteins in P3 LTMR subtypes (Figures 1G).

Among genes encoding transmembrane proteins implicated in developmental processes, *Pcdhg* subcluster genes stood out because they exhibited interesting differences in the expression levels of isoforms, with *Pcdhga7* and *Pcdhgc3* isoforms being expressed at the highest levels (Figure 1H). Expression of *Pcdhgs* isoforms in P3 DRG neurons was confirmed with single-molecule RNA fluorescent *in situ* hybridization (smRNA-FISH), using probes against 5 representative isoforms (Figure S1E). While *Pcdhgc3* is evenly expressed throughout the DRG, *Pcdhga7* is enriched in a subset of DRG neurons (Figure S1E). We also examined expression of these representative isoforms in the P3 and P21 spinal cord, using vGluT2 as a marker for spinal cord excitatory neurons (Figure 1I). Similar to its DRG expression, *Pcdhgc3* was detected throughout the spinal cord, with 67.7 ± 3.5 % of excitatory neurons expressing this isoform (Figure 1J). *Pcdhgc4* was also detected at a high level in the spinal cord, consistent with its role in spinal cord neuron survival (Garrett et al., 2019). In addition, there is a significant decrease in the level of expression of three out of five isoforms between P3 and P21 (Figure 1J). The expression patterns of *Pcdhg* isoforms in developing somatosensory and spinal cord neurons led us to hypothesize that they may contribute to somatosensory neuron central and peripheral axon development.

### *Pcdhgs* function in primary sensory neurons for normal tactile behaviors and sensorimotor integration

Although *Pcdhgs* are broadly expressed in the developing DRG and dispensable for DRG neuron survival (Prasad and Weiner, 2011), their roles in LTMR development and function remain unknown. To examine this, we used a *Pcdhg* conditional allele (*Pcdhg^fcon3^*), to bypass the neonatal lethality associated with constitutive *Pcdhg* family deletion, and *Advillin^Cre^*to selectively delete all 22 *Pcdhg* isoforms in the DRG during embryonic development (Hasegawa et al., 2007; Lefebvre et al., 2008; Prasad et al., 2008). In the *Pcdhg^fcon3^*allele, GFP is fused to the carboxy-terminal constant exon 3 shared by all isoforms and is flanked by loxP sites, such that loss of GFP expression indicates of Cre-mediated excision of the constant exon 3. As expected, *Advillin^Cre^* efficiently mediated deletion of *Pcdhgs* in virtually all DRG neurons but not in the spinal cord (Figure 2A).

**Figure 2.**
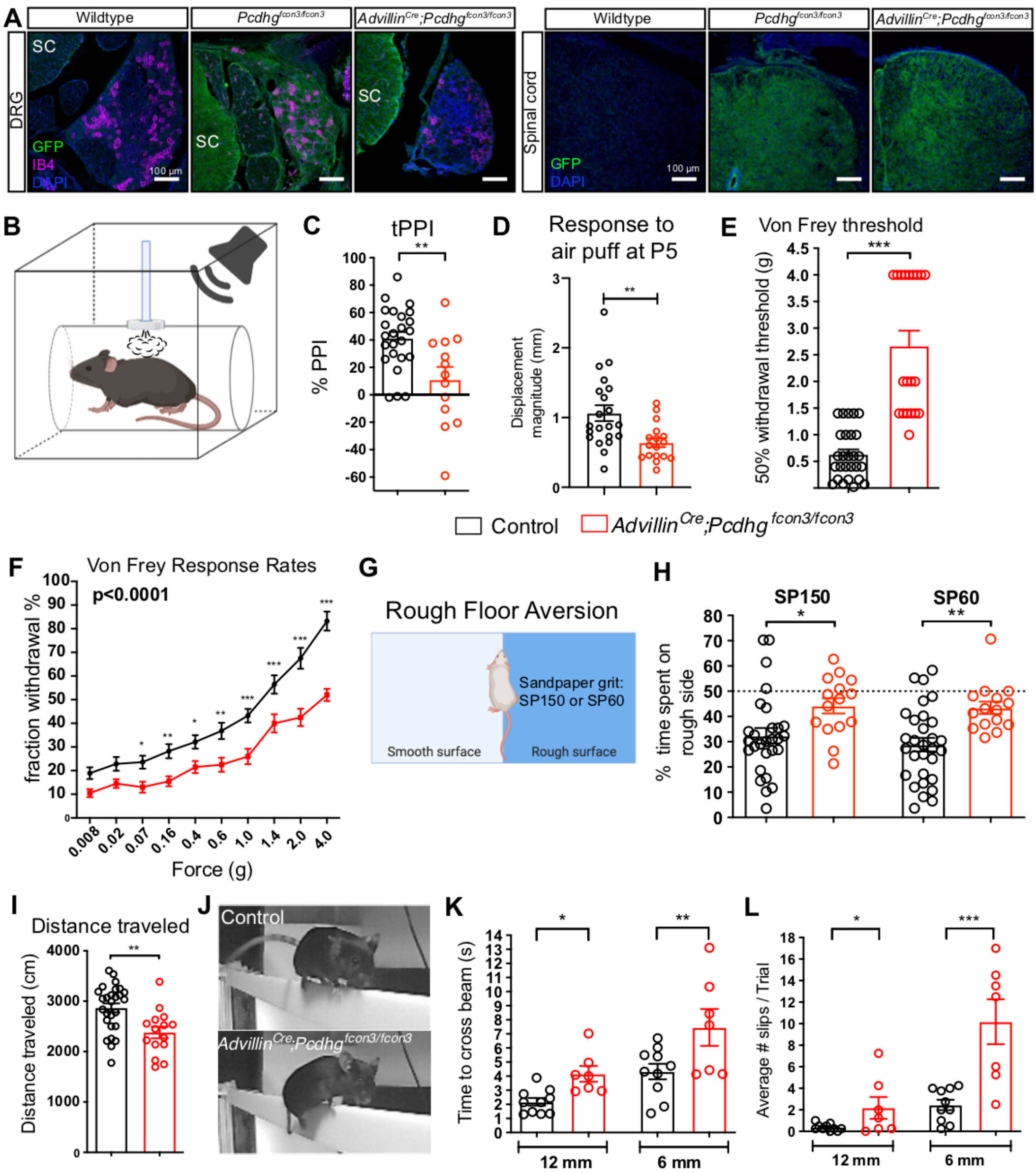
*Pcdhgs* function in primary sensory neurons for normal tactile behaviors and sensorimotor integration. **(A)** IHC images of DRG and spinal cord showing the GFP fused Pcdhg proteins in wildtype (negative control), *Pcdhg^fcon3/fcon3^* (positive control) and *Advillin^Cre^;Pcdhg^fcon3/fcon3^* mice. IB4 labels lamina IIi nonpeptidergic nociceptors in the DRG. **(B)** Diagram for the tactile PPI behavior assay. **(C)** Percentage of inhibition of startle response to 125 dB noise in littermate control and *Advillin^Cre^;Pcdhg^fcon3/fcon3^* mice when the startle noise is preceded by a light air puff of 0.9 PSI (500 ms ISI). Student’s unpaired t test. **(D)** Quantification of the average movement of the back in response to 1.0 PSI air puff applied to the back hairy skin of P5 littermate control and *Advillin^Cre^;Pcdhg^fcon3/fcon3^*pups. Student’s unpaired t test. **(E and F)** Von Frey thresholds (E) and response rates (F) for littermate control and *Advillin^Cre^;Pcdhg^fcon3/fcon3^* mice. Two-way ANOVA. Fisher’s LSD post hoc test. **(G)** Diagram for the rough floor aversion assay. **(H)** Percentages of time spent on rough side for littermate control and *Advillin^Cre^;Pcdhg^fcon3/fcon3^* mice. Student’s unpaired t test. **(I)** Distance traveled in an open field test. Student’s unpaired t test. **(J)** Example images of the balance beam test. **(K and L)** Total time it took for each animal to cross the beam (K) and average number of slips per trial (L) for littermate control and *Advillin^Cre^;Pcdhg^fcon3/fcon3^* mice. Student’s unpaired t test. Each dot represents an animal. ns, not significant; *p < 0.05; **p < 0.01; ***p < 0.001.

To determine whether *Pcdhgs* in somatosensory neurons are essential for LTMR function and somatosensation, tactile sensitivity of *Advillin^Cre^;Pcdhg^fcon3/fcon3^* mutant mice was tested. At weaning age, *Advillin^Cre^;Pcdhg^fcon3/fcon3^*mutants exhibited reduced body weight, but quickly gained weight to a normal range by 6 weeks of age when behavioral assays were conducted (Figure S2A). A tactile prepulse inhibition of the startle reflex assay (tactile PPI) was performed on mutant and littermate control mice (Orefice et al., 2019; 2016) to assess hairy skin sensitivity. For the tactile PPI assay, a light air puff prepulse (0.9 PSI) is applied to back hairy skin and this mechanical “pre-pulse” stimulus is followed by a startle pulse of broadband white noise (125 dB) that elicits an acoustic startle response (Figure 2B). Mechanosensory stimulation by the air puff prepulse attenuates the acoustic startle response in control mice (Figure 2C). Adult *Advillin^Cre^;Pcdhg^fcon3/fcon3^* mutant mice exhibited a reduction in tactile PPI (Figure 2C) but not the response to air puff alone (Figure S2B), suggesting a reduction in hairy skin mechanosensation. On the other hand, the mutants performed normally in an acoustic version of PPI (Figure S2C). To extend this finding and determine whether hairy skin sensitivity defects emerge during postnatal development, repeated air puffs (1.0 PSI) were applied to the back hairy skin of P5 pups, and the magnitude of evoked movement was measured. Consistent with the adult tactile PPI findings, P5 *Advillin^Cre^;Pcdhg^fcon3/fcon3^* mutants exhibited reduced reactivity to air puffs compared to littermate controls (Figure 2D).

We also asked whether glabrous skin sensitivity to mechanical stimuli is altered in *Advillin^Cre^;Pcdhg^fcon3/fcon3^*mice. To do so, mice were subjected to von Frey filament stimulation of hindpaw glabrous skin (Frey, 1896). While littermate controls withdrew their paws in a manner proportional to the forces tested, *Advillin^Cre^;Pcdhg^fcon3/fcon3^*mice displayed lower hindpaw withdrawal responses to both low and high forces and a higher withdrawal threshold (Figures 2E and 2F). The rough floor aversion test was performed as an additional measurement of glabrous skin sensitivity (Figure 2G) (Choi et al., 2020; Zimmerman et al., 2019). Here, *Advillin^Cre^;Pcdhg^fcon3/fcon3^*mice exhibited decreased aversion to rough textured floors compared to littermate controls (Figure 2H).

Because somatosensory feedback contributes to the control of locomotion and corrective movements (Paixão et al., 2019; Panek et al., 2014; Rossignol et al., 2006), we also examined general locomotion in *Advillin^Cre^;Pcdhg^fcon3/fcon3^*mice and their littermate controls using the open field assay. *Advillin^Cre^;Pcdhg^fcon3/fcon3^*mice exhibited a decrease in both the total distance traveled (Figure 2I) and the percentage of time spent in the center of the chamber during the 10 minute testing period (Figure S2D). This decrease in the time spent in the center of the chamber may not represent an anxiety-like behavior, as the mutant animals performed normally during the elevated plus maze (EPM) test, a measurement of anxiety-like behavior (Figure S2E) (McGill et al., 2006). The balance beam test was used to assess sensorimotor integration. When traversing a narrow elevated beam, *Advillin^Cre^;Pcdhg^fcon3/fcon3^* mice spent more time crossing the beam (Figures 2J and 2K) and displayed more hindlimb slips than littermate control mice (Figures 2L).

Taken together, these findings demonstrate that mice lacking *Pcdhgs* in somatosensory neurons are hyposensitive to mechanical stimuli acting on different skin regions during both postnatal development and adulthood and they exhibit disrupted sensorimotor function.

### *Pcdhgs* function in primary somatosensory neurons for central synapse formation

The aberrant mechanosensory behaviors of *Advillin^Cre^;Pcdhg^fcon3/fcon3^*mice raised the question of the nature of the dysfunction in mechanoreceptors caused by deletion of *Pcdhgs* in these neurons. Therefore, we next asked whether *Pcdhgs* regulate synapse formation or other aspects of central and peripheral axon development. In the dorsal horn LTMR-RZ, ∼60% of the vGluT1^+^ glutamatergic terminals are somatosensory neuron presynaptic terminals, while ∼40% of vGluT1^+^ terminals are corticospinal presynaptic terminals (Abraira et al., 2017). vGluT1 and Homer1 double staining was used to visualize these two types of excitatory glutamatergic synapses in this region. At P21, the density of both vGluT1 and Homer1 puncta was reduced in *Advillin^Cre^;Pcdhg^fcon3/fcon3^* mutants, with vGluT1 density reduced by 27.4 ± 11.6 % and Homer1^+^ excitatory synapse density reduced by 62.3 ± 9.1 % (Figures 3A-3D), compared to control littermates. We also asked whether this decrease in excitatory synapses is observed at a functional level. For this, whole cell recordings of spinal cord neurons in lamina III from *Advillin^Cre^;Pcdhg^fcon3/fcon3^* mutant mice and littermate controls were used to measure spontaneous miniature excitatory postsynaptic currents (mEPSCs) (Figure 3E). Consistent with the histological analysis, mutant spinal cord neurons displayed a ∼50% reduction in mEPSC frequency, but no change in mEPSC amplitudes (Figures 3E, F and S3A). To determine when the reduction of synapse density emerges, excitatory synaptic densities during neonatal and postnatal times were examined. At P0, vGluT1^+^ puncta could not be reliably detected, presumably due to the low level of vGluT1 protein in presynaptic terminals at this age. However, at P4 the density of vGluT1^+^ puncta appeared normal in the *Advillin^Cre^;Pcdhg^fcon3/fcon3^*mutants (Figures 3A and 3C). On the other hand, the density of Homer1^+^ puncta in the mutants appeared normal at P0, while at P4 it was decreased by 59.0 ± 15.1% compared to control littermates (Figure 3D), consistent with the reduced behavioral response to air puff in P5 mutants (Figure 2D). These findings suggest that loss of *Pcdhgs* in somatosensory neurons leads to a reduction of both vGluT1^+^ presynaptic terminals and Homer1^+^ postsynaptic puncta within the LTMR-RZ and a concomitant loss of functional synapses. This synapse reduction is unlikely to be caused by abnormal targeting of the sensory terminals, as the overall pattern of sensory axonal projections in the dorsal horn and the distribution of vGluT1^+^ presynaptic terminals in *Advillin^Cre^;Pcdhg^fcon3/fcon3^* mutants appeared normal (Figures 3B and S3B).

**Figure 3.**
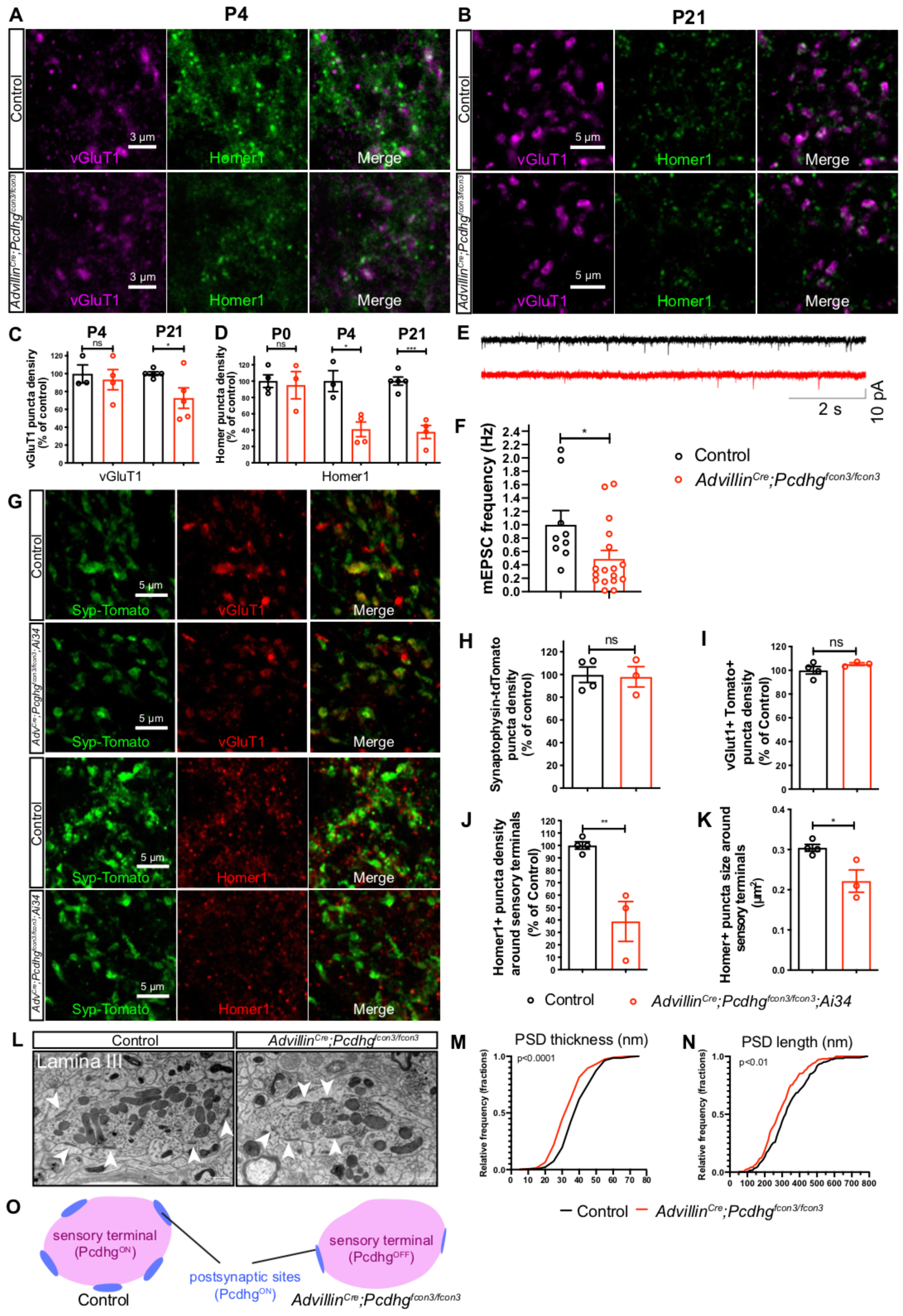
*Pcdhgs* function in primary sensory neurons for postsynaptic specialization of synapses between sensory axon terminals and spinal cord neurons. **(A and B)** IHC images of spinal cord dorsal horn lamina III from P4 (A) and P21 (B) littermate control and *Advillin^Cre^;Pcdhg^fcon3/fcon3^*mice. Sections were immunostained for vGluT1 (presynaptic terminals for Aβ-LTMRs, Aδ-LTMRs, and corticospinal axons) and Homer1 (postsynaptic densities of glutamatergic synapses) to mark the synaptic connections for LTMR and corticospinal presynaptic terminals. (**C and D**) Normalized densities of vGluT1^+^ (C) and Homer1^+^ (D) puncta from littermate control and *Advillin^Cre^;Pcdhg^fcon3/fcon3^* mice. Student’s unpaired t test. **(E)** Representative spontaneous mEPSC traces recorded in random spinal cord neurons in lamina III from P13-P16 littermate control (n = 3 animals) and *Advillin^Cre^;Pcdhg^fcon3/fcon3^*mice (n = 4 animals). **(F)** Presynaptic *Pcdhg* deletion decreases the frequency of spontaneous mEPSCs in the spinal cord neurons. Each dot is the mEPSC frequency of a spinal cord neuron. Mann-Whitney test. **(G)** IHC images of spinal cord lamina III from P21 littermate control and *Advillin^Cre^;Pcdhg^fcon3/fcon3^;Ai34* mice, in which somatosensory neuron presynaptic terminals are labeled with synaptophysin-tdTomato. **(H-K)** Normalized densities of synaptophysin-tdTomato (Ai34) puncta density (H) and vGluT1^+^ Tomato^+^ double positive puncta (I). The size and normalized density of Homer1^+^ puncta around Tomato^+^ sensory terminals is quantified in (K) and (J), respectively. Each dot represents one animal. Student’s unpaired t test. **(L)** EM images of synaptic glomeruli from lamina III in spinal cord dorsal horn from a control animal and a *Advillin^Cre^;Pcdhg^fcon3/fcon3^* animal. Arrowheads point to the postsynaptic sites formed within the glomeruli. Scale bars represent 500 nm. **(M and N)** Quantifications of PSD thickness (M, control 42.1 ± 0.6 nm, 304 synapses from 3 animals; *Advillin^Cre^;Pcdhg^fcon3/fcon3^*: 32.4 ± 0.6 nm, 286 synapses from 3 animals). Quantifications of PSD length (N, control 340.0 ± 7.1 nm; *Advillin^Cre^;Pcdhg^fcon3/fcon3^*: 299.3 ± 6.2 nm). Kolmogorov-Smirnov test. **(O)** Summary of the synaptic formation phenotype in the *Advillin^Cre^;Pcdhg^fcon3/fcon3^*mutants. Fewer and smaller synapses form between sensory terminals and spinal cord neurons in the *Advillin^Cre^;Pcdhg^fcon3/fcon3^* mutants. ns, not significant; *p < 0.05; **p < 0.01; ***p < 0.001.

Since vGluT1 labels both primary somatosensory neuron and corticospinal neuron terminals in the LTMR-RZ, we tested whether the disruption in vGluT1 labeling observed in the *Advillin^Cre^;Pcdhg^fcon3/fcon3^*mutants is a reflection of altered somatosensory neuron synapses, corticospinal neuron synapses, or both. To selectively visualize primary somatosensory neuron terminals, we generated *Advillin^Cre^;Pcdhg^fcon3/fcon3^;Ai34* animals, in which all somatosensory neurons express the presynaptic marker synaptophysin-tdTomato (Figure 3G). Interestingly, the density of primary sensory neuron synaptic terminals in the mutants was comparable to that of controls at P21 (Figure 3H). Similarly, the density of vGluT1 and tdTomato double positive puncta was uncompromised in the mutants (Figure 3I), suggesting that the reduction in the number of vGluT1^+^ puncta in the mutants is not caused by a decrease in sensory neuron presynaptic terminals, but rather a decrease in corticospinal terminals (also see Figures 7H-K). On the other hand, the density of Homer1^+^ puncta surrounding tdTomato^+^ sensory terminals was reduced by 61.2 ± 13.9% in *Advillin^Cre^;Pcdhg^fcon3/fcon3^;Ai34* animals, similar to the overall Homer1^+^ puncta reduction observed throughout lamina III (Figure 3J). Moreover, the size of Homer1^+^ puncta associated with sensory neuron synapses was decreased in *Advillin^Cre^;Pcdhg^fcon3/fcon3^;Ai34* animals (Figure 3K), suggesting that PSDs are smaller in the mutants. To address this further, electron microscopy was used to examine ultrastructural features of excitatory synapses surrounding dorsal horn synaptic glomeruli, which are comprised of axon terminals of primary sensory afferents and synaptic interactions with surrounding spinal cord neurons (Figure 3L) (Ribeiro-da-Silva et al., 1985; Zhang et al., 2019). Indeed, the thickness and length of PSDs associated with synaptic glomeruli in lamina III were reduced in *Advillin^Cre^;Pcdhg^fcon3/fcon3^* mice by 23.1 ± 1.9 % and 12.0 ± 2.8 %, respectively (Figures 3M and 3N), while the overall morphology of glomeruli of mutant mice appeared normal.

Together, these finding show that *Pcdhgs* function in somatosensory neurons to establish proper numbers, function, and ultrastructural properties of postsynaptic specializations formed between somatosensory neurons and dorsal horn neurons (Figure 3O).

### *Pcdhgs* function in primary sensory neurons for LTMR axonal branching in the skin

Pcdhg proteins are found throughout neurons, including in axons and dendrites (Lefebvre et al., 2008; Li et al., 2010; Phillips et al., 2003), and in CNS neurons they regulate many aspects of dendrite development, including dendrite self-avoidance and arborization (Garrett et al., 2012; Gibson et al., 2014; Keeler et al., 2015; Lefebvre et al., 2012; 2015; Molumby et al., 2016). To ask whether loss of *Pcdhgs* in DRG neurons affects LTMR peripheral axon development, the patterns of LTMR endings in the skin of *Advillin^Cre^;Pcdhg^fcon3/fcon3^* mutants and control littermates were visualized using whole mount immunohistochemistry. In back hairy skin, Aβ field-LTMRs and Aβ RA-LTMRs form NFH^+^ circumferential and lanceolate endings around hair follicles, respectively (Bai et al., 2015; Li et al., 2011). In both controls and *Advillin^Cre^;Pcdhg^fcon3/fcon3^* mutants, all guard hairs associated with Merkel cell complexes were innervated by NFH^+^ Aβ field-LTMRs and Aβ RA-LTMRs endings (Figures 4A-4C). However, non-guard hairs, which include awl/auchene and zigzag hairs, exhibited reduced innervation by NFH^+^ Aβ field-LTMRs and Aβ RA-LTMRs in *Advillin^Cre^;Pcdhg^fcon3/fcon3^* mutants (Figures 4B and 4C). Of note, the morphology of sensory neuron endings and associated terminal Schwann cells surrounding hair follicles were unchanged (Figures S4A-S4D), and DRG neuron survival is unaffected by the absence of *Pcdhgs* (Prasad and Weiner, 2011). Because individual Aβ field-LTMRs exhibit tiling and innervate hair follicles in a non-overlapping manner (Kuehn et al., 2019), this finding suggests that Aβ field-LTMRs have reduced axonal branching in the skin. Likewise, the reduction of NFH^+^ lanceolate ending innervation suggests a similar defect in Aβ RA-LTMR axonal branching.

**Figure 4.**
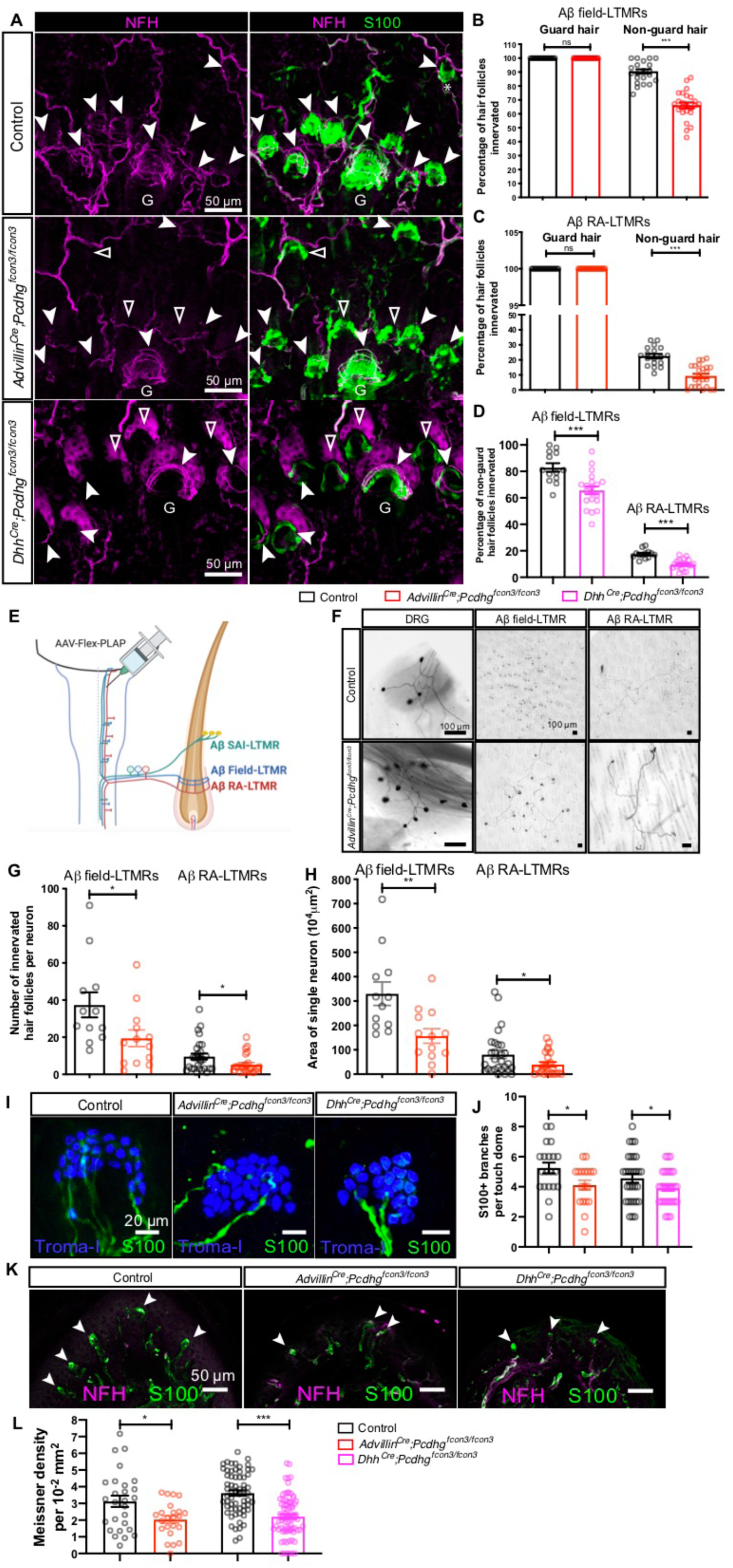
*Pcdhgs* function in LTMRs for axonal branching in the skin. **(A)** Whole-mount immunostainings of adult back hairy skin sections from control and *Advillin^Cre^;Pcdhg^fcon3/fcon3^* mice. Aβ field-LTMRs form circumferential endings around hair follicles and are NFH^+^ (noted by white arrowheads). Aβ RA-LTMRs form lanceolate endings around guard hair follicles (noted as “G”) and awl/auchene hairs. TSCs are labeled using S100 immunostaining (green), which helps visualize the location of LTMR innervation around hair follicles. Hair follicles without any NFH^+^ Aβ field-LTMRs are denoted by triangles. **(B-D)** Quantification of the percentage of guard and non-guard hair follicles innervated by NFH^+^ Aβ field-LTMRs circumferential endings and Aβ RA-LTMRs lanceolate endings for *Advillin^Cre^;Pcdhg^fcon3/fcon3^* mice (n = 958 hair follicles from 3 control animals and n = 1357 hair follicles from 3 *Advillin^Cre^;Pcdhg^fcon3/fcon3^*mice) and *Dhh^Cre^;Pcdhg^fcon3/fcon3^* mice (n = 597 hair follicles from 3 control animals and n = 863 hair follicles from 4 *Dhh^Cre^;Pcdhg^fcon3/fcon3^* mice). Student’s unpaired t test. Each dot represents an imaging field. **(E)** Control and *Advillin^Cre^;Pcdhg^fcon3/fcon3^* mice were injected with AAV-Retro-Flex-PLAP virus to retrogradely label DRG neurons that project to the DCN. **(F)** Example whole-mount AP images of the DRG and back hairy skin. **(G and H)** Quantification of the number of innervated non-guard hair follicles (G) and the area of innervation (H) by Aβ field-LTMRs (n = 12 neurons from 6 control animals and n = 13 neurons from 5 *Advillin^Cre^;Pcdhg^fcon3/fcon3^*mice) and Aβ RA-LTMRs (n = 28 neurons from 6 control animals and n = 27 neurons from 6 *Advillin^Cre^;Pcdhg^fcon3/fcon3^* mice). Student’s unpaired t test. Dots represent individual neurons. **(I and J)** Example whole-mount immunostaining images of the Merkel cell complex. TSCs wrapping around major Aβ SAI-LTMRs branches are labeled with S100 (green), and Merkel cells are labeled with Troma-I (blue). Quantifications of the major S100^+^ branches from *Advillin^Cre^;Pcdhg^fcon3/fcon3^* mice (n = 3 control animals and n = 4 mutant animals) and *Dhh^Cre^;Pcdhg^fcon3/fcon3^* mice (n = 3 control animals and n = 4 mutant animals) are shown in (J). Each dot represents a touch dome. Student’s unpaired t test. **(K and L)** Example immunostaining images of the forepaw glabrous skin. Meissner corpuscles are labeled by S100 (green, for visualizing lamellar cells) and NFH (magenta, for visualizing Aβ RA-LTMRs). Quantifications of the number of Meissner corpuscles normalized by the area of epidermis from *Advillin^Cre^;Pcdhg^fcon3/fcon3^* (n = 4 control animals and n = 4 mutant animals) and *Dhh^Cre^;Pcdhg^fcon3/fcon3^* mice (n = 3 control animals and n = 4 mutant animals) are shown in (L). Dots represent individual skin sections. Student’s unpaired t test. ns, not significant; *p < 0.05; **p < 0.01; ***p < 0.001.

To complement the immunohistological analysis and directly visualize axonal branching patterns of individual Aβ field-LTMRs and Aβ RA-LTMRs in hairy skin, sparse labeling of LTMRs was done by injecting a Cre-dependent alkaline phosphatase reporter virus into the dorsal column nuclei (DCN) of the brainstem, where Aβ field-LTMRs, Aβ RA-LTMRs and Aβ SAI-LTMRs project their axons (Figure 4E) (Bai et al., 2015). Then, whole mount alkaline phosphatase staining was performed to visualize the cell bodies in the DRG and axonal arbors of individual LTMRs in the skin (Figure 4F). This analysis revealed a reduction in both the number of innervated hair follicles per neuron and the area innervated by individual Aβ field-LTMRs and Aβ RA-LTMRs in *Advillin^Cre^;Pcdhg^fcon3/fcon3^* mutants (Figures 4G and 4H). Otherwise, the overall axonal arborization patterns appeared normal and no redundant innervation of hair follicles by the same neuron was observed, indicating that there is not a self-avoidance-like deficit in these neurons in the mutant mice (Figure 4F). Moreover, Aβ SAI-LTMRs associated with clusters of Merkel cells (touch domes) that surround guard hairs in hairy skin exhibited a small but significant reduction in the number of the S100^+^ major branches in the *Advillin^Cre^;Pcdhg^fcon3/fcon3^* mutants (Figures 4I and 4J), while the total number of Merkel cells in touch domes remained the same (Figure S4E). In glabrous skin, the density of Meissner corpuscles was also reduced in the mutants, compared to control littermates (Figures 4K and 4L), suggesting a decrease in axonal branching. Together, these experiments reveal that *Pcdhgs* function cell autonomously to promote Aβ LTMR axon branching in both hairy and glabrous skin.

LTMR peripheral axons closely associate with peripheral glial cells (Reed et al., 2021; Suazo et al., 2022; Zimmerman et al., 2014), although the significance of this association in LTMR development is unclear. To determine if *Pcdhgs* mediate homophilic adhesion between nascent peripheral axons and glial cells to promote axonal branching, *Dhh^Cre^;Pcdhg^fcon3/fcon3^*mutants were generated to selectively delete *Pcdhgs* in developing Schwann cells of the peripheral nervous system, beginning at E12 (Jaegle et al., 2003) (Figure S4G). Remarkably, *Dhh^Cre^;Pcdhg^fcon3/fcon3^* mutants phenocopied *Advillin^Cre^;Pcdhg^fcon3/fcon3^* mutants with respect to the hairy and glabrous skin LTMR innervation deficits (Figures 4A, 4D, and 4I-4L). On the other hand, excitatory synapses in the spinal cord dorsal horn were unaffected in *Dhh^Cre^;Pcdhg^fcon3/fcon3^* mutants at P21 (Figures S4H-S4K). These finding support a model in which Pcdhgs mediate homophilic interactions between nascent LTMR axonal branches and developing glial cells to promote axonal branching. Furthermore, deficits in LTMR branching patterns and central synapses in the *Pcdhg* mutants are dissociable; defects in LTMR peripheral branching patterns occur independently of alterations of LTMR central synapses.

We also tested whether Pcdhgs function to maintain central synapses and peripheral axonal branching after they are established, using the tamoxifen-sensitive *Advillin^CreERT2^* mouse line (Lau et al., 2011) to delete *Pcdhgs* in somatosensory neurons of juvenile mice (*Advillin^CreERT2^*; *Pcdhg^fcon3/fcon3^* mice, 1mg per day, P15-P19; Figures S5A and S5B). Interestingly, late postnatal deletion of *Pcdhgs* recapitulated the reduction of excitatory synapses observed in *Advillin^Cre^;Pcdhg^fcon3/fcon3^* mutants (Figures S5C and S5E-G), while sensory and corticospinal presynaptic vGluT1^+^ terminals did not change significantly (Figure S5D). In hairy skin, *Advillin^CreERT2^;Pcdhg^fcon3/fcon3^*animals exhibited fewer non-guard hairs innervated by NFH^+^ Aβ field-LTMRs and Aβ RA-LTMRs (Figures S5H and S5I), as well as fewer Aβ SAI-LTMR branches (Figures S5J and S5K). These observations suggest that Pcdhg signaling maintains both LTMR central synapses and peripheral axonal arborization patterns after they are formed.

### Differential requirements of *Pcdhg* isoforms for synapse formation and peripheral axonal branching, with Pcdhgc3 solely required for synapse formation

The diversity of *Pcdhg* isoforms raised the question of whether one or more of the same 22 *Pcdhg* isoforms mediate both synapse formation and axonal branching, or whether distinct isoforms contribute to these developmental processes. To address this, somatosensory neuron synapses and axonal branching patterns were examined using a range of *Pcdhg* mutant mice that lack one or more isoforms (Figure 5A). All mutants tested were viable to adulthood because they have intact *Pcdhgc4*, which encodes the isoform required for postnatal viability (Garrett et al., 2019).

**Figure 5.**
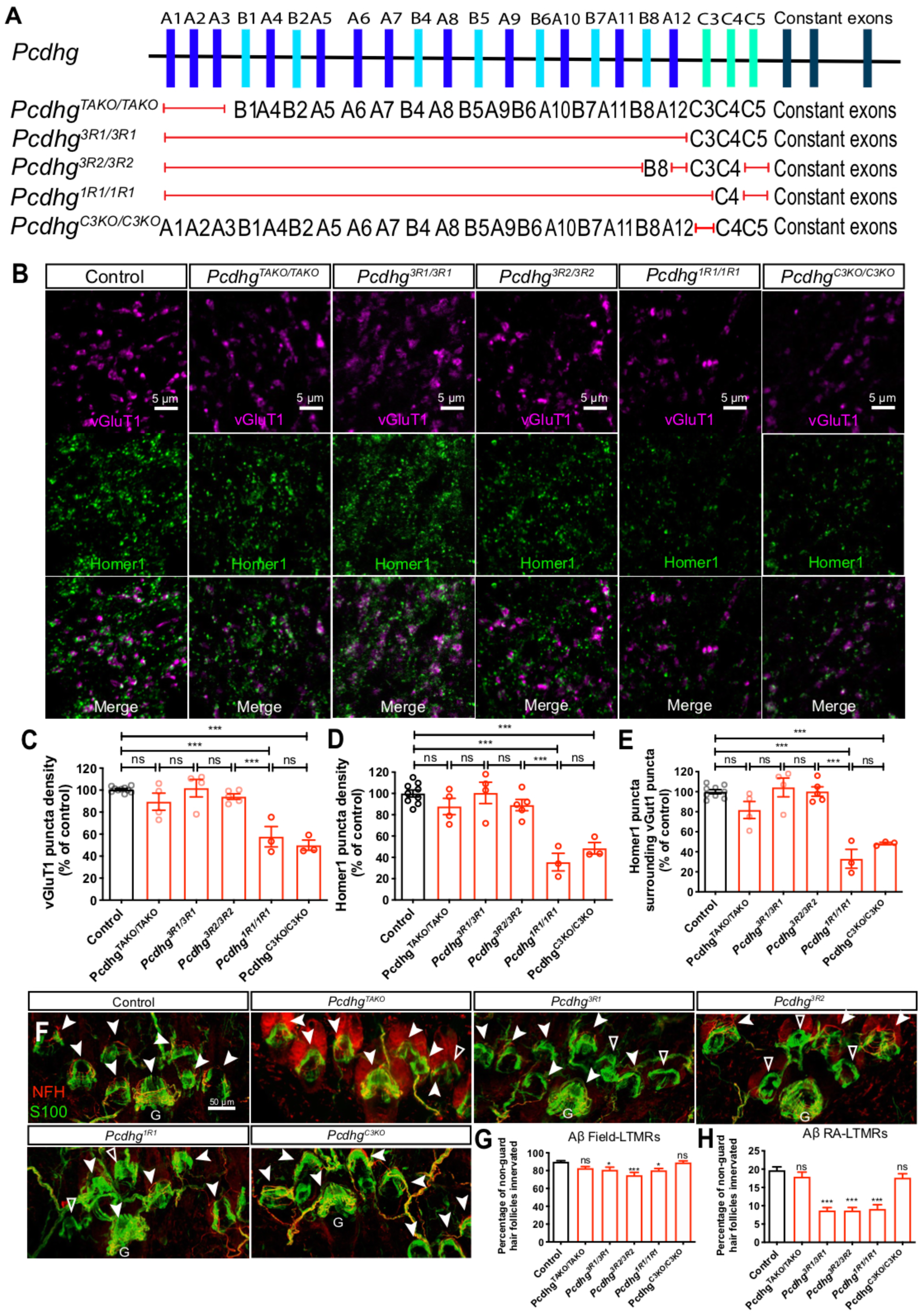
Differential requirements of Pcdhg isoforms for synapse formation and peripheral axonal branching. **(A)** Summary of the *Pcdhg* mutants used. Red lines indicate the corresponding genes are disrupted, while the genes listed are not disrupted. **(B)** IHC images of spinal cord dorsal horn lamina III from P21 control and *Pcdhg* mutants. Sections were immunostained for vGluT1 and Homer1 to mark the synaptic connections around in LTMR and corticospinal presynaptic terminals. (**C-E)** Normalized densities of vGluT1^+^ (C) and Homer1^+^ (D) puncta from P21 control and various *Pcdhg* mutant mice. (E) shows the normalized densities of Homer1^+^ puncta that surrounds vGluT1^+^ puncta. one-way ANOVA with Tukey’s post hoc test. Each dot represents average value of an animal. **(F)** Whole-mount immunostaining of adult back hairy skin sections from control and various *Pcdhg* mutant mice. Aβ field-LTMRs (noted by white arrowheads) and Aβ RA-LTMRs (innervation around the guard hair is denoted by “G”) are NFH^+^. Hair follicles without any NFH^+^ Aβ field-LTMRs are denoted by empty arrowheads. TSCs are labeled using S100 immunostaining (green). Scale bar represents 50 μm. **(G and H)** Quantification of the percentage of non-guard hair follicles innervated by NFH^+^ Aβ field-LTMRs circumferential endings (G) and Aβ RA-LTMRs lanceolate endings (H). n = 649 hair follicles from 5 wildtype animals; n = 336 hair follicles from 2 *Pcdhg^TAKO^*animals; n = 710 hair follicles from 4 *Pcdhg^3R1^* animals; n = 673 hair follicles from 5 *Pcdhg^3R2^* animals; n = 614 hair follicles from 3 *Pcdhg^C3KO^* animals. One-way ANOVA with Tukey’s post hoc test. ns, not significant; *p < 0.05; **p < 0.01; ***p < 0.001.

The triple A-type isoform knockout (TAKO) *Pcdhg^TAKO/TAKO^*mutant showed normal numbers of dorsal spinal cord synapses and Aβ field-LTMR and Aβ RA-LTMR endings in hairy skin (Figures 5B-5H) (Chen et al., 2012). Interestingly, while the *Pcdhg^3R1/3R1^* (*Pcdhgc3*, *Pcdhg4* and *Pcdhgc5* intact) and *Pcdhg^3R2/3R2^* (*Pcdhgb8*, *Pcdhgc3* and *Pcdhgc4* intact) mutants exhibited normal vGluT1 and Homer1 excitatory synapse density in the dorsal horn, they displayed Aβ field-LTMR and Aβ RA-LTMR innervation pattern defects in the hairy skin (Figures 5B-5H). The phenotypes of these two mutants thus corroborate the conclusion of experiments described above that the central and peripheral phenotypes are dissociable and further indicate that the critical isoform(s) for synapse formation is intact in these two mutants (i.e., *Pcdhgc3* and *Pcdhgc4*). Therefore, to determine whether *Pcdhgc3* or *Pcdhgc4* promotes synapse formation, we next examined *Pcdhg^1R1/1R1^* mutants in which only *Pcdhgc4* is intact and expressed. *Pcdhg^1R1/1R1^* mutants showed deficits in both synapse formation and peripheral axonal branching (Figures 5B-5E), suggesting that *Pcdhgc3* is the sole isoform essential for synapse formation.

To further test this idea, additional genetic loss-of-function and gain-of-function approaches were taken. First, we found that the *Pcdhg^C3KO/C3KO^*mutant, which lacks only *Pcdhgc3,* phenocopied the synapse deficits observed in *Advillin^Cre^;Pcdhg^fcon3/fcon3^* mice and showed a reduction in excitatory synapse density; however this mutant did not exhibit an alteration of Aβ field-LTMR and Aβ RA-LTMR innervation patterns in hairy skin (Figures 5B-5H). Second, we asked whether expressing Pcdhgc3 in somatosensory neurons of *Advillin^Cre^;Pcdhg^fcon3/fcon3^* mutants is sufficient to rescue the synapse formation phenotype. For this, the *cC3* mouse line, in which the Pcdhgc3 isoform fused to a fluorescent protein (mCherry) is expressed in a Cre-dependent manner (*ROSA26-CAG::lox-Stop-lox-Pcdhgc3-mCherry*) (Lefebvre et al., 2012), was used to conditionally express Pcdhgc3 in somatosensory neurons (Figure 6A). Thus, in *Advillin^Cre^;Pcdhg^fcon3/fcon3^;cC3* animals, Cre simultaneously deletes all 22 endogenous *Pcdhg* genes and activates Pcdhgc3-mCherry expression in all somatosensory neurons. For comparison, we also asked whether Pcdhga1 is sufficient for synapse formation by testing whether expressing the Pcdhga1 isoform (*ROSA26-CAG::lox-Stop-lox-Pcdhga1-mCherry, cA1*) in the DRGs of *Advillin^Cre^;Pcdhg^fcon3/fcon3^;cA1* animals can rescue the synapse formation deficits observed in the *Pcdhg* cluster knockout. Strikingly, expression of Pcdhgc3, but not Pcdhga1, in somatosensory neurons of *Advillin^Cre^;Pcdhg^fcon3/fcon3^* mice rescued the synapse formation deficits observed in these mutants (Figures 6B-6D). Taken together, these experiments demonstrate that Pcdhgc3 is the key isoform that functions in LTMRs to promote central synapse formation in the spinal cord, whereas isoforms other than Pcdhgc3 mediate proper LTMR peripheral axonal branching.

**Figure 6.**
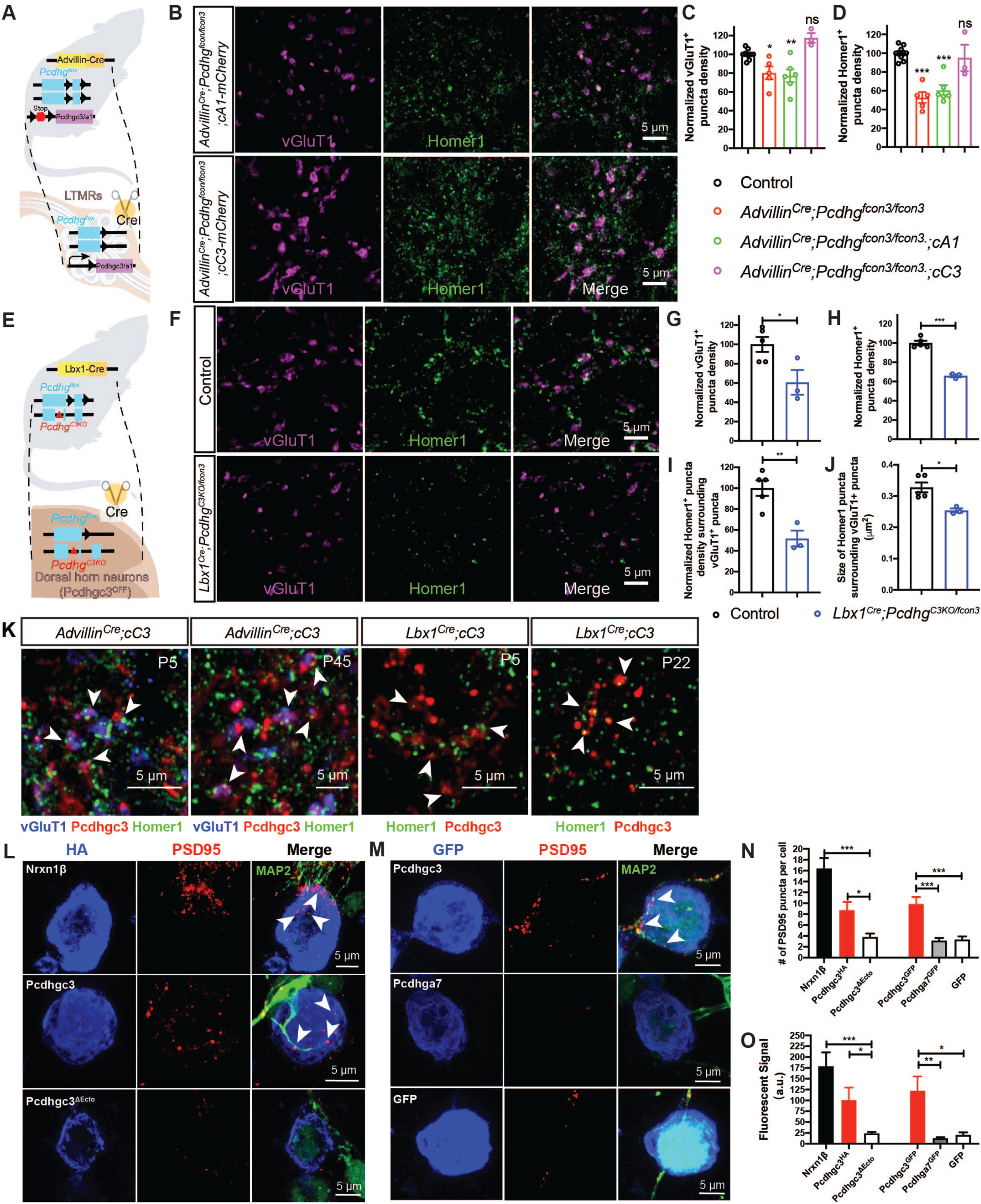
Pcdhgc3 is the only isoform that mediates sensory neuron synapse formation *in vivo* and can promote postsynaptic specialization *in vitro*. **(A)** Experiment design for expressing single *Pcdhg* isoforms (Pcdhga1 or Pcdhgc3) in *Advillin^Cre^;Pcdhg^fcon3/fcon3^* mice. Cre simultaneously deletes all 22 endogenous *Pcdhg* genes and activates the expression of *Pcdhga1-mCherry* or *Pcdhgc3-mCherry* in the DRG. **(B)** IHC images of spinal cord dorsal horn lamina III from *Advillin^Cre^;Pcdhg^fcon3/fcon3^;cA1* and *Advillin^Cre^;Pcdhg^fcon3/fcon3^;cC3* mice. **(C and D)** Normalized densities of vGluT1^+^ (C) and Homer1^+^ (D) puncta, showing that expressing of *Pcdhgc3-mCherry*, but not *Pcdhga1-mCherry*, in the DRGs can fully restore the synapse formation deficits in the *Advillin^Cre^;Pcdhg^fcon3/fcon3^*mutants. Each dot on the plot indicates a normalized value for an animal. One-way ANOVA test. **(E)** Experiment design for deleting both copies of the *Pcdhgc3* isoforms only in the dorsal horn neurons in the *Lbx1^Cre^;Pcdhg^C3KO/fcon3^* mice. Littermate animals with or without *Lbx1^Cre^* were used as controls. **(F)** IHC images of spinal cord dorsal lamina III from control and *Lbx1^Cre^;Pcdhg^C3KO/fcon3^* mice. **(G-J)** Quantifications of the synapses in lamina III, showing that deleting *Pcdhgc3* in the dorsal horn neurons led to reduced densities of vGluT1^+^ (G), Homer1^+^ (H) puncta, as well as the density of Homer1^+^ puncta surrounding vGluT1^+^ terminals (I). Similarly, the average size of Homer1^+^ puncta is reduced in *Lbx1^Cre^;Pcdhg^C3KO/fcon3^* mice (J). Each dot on the plot indicates a normalized value for an animal. Student’s unpaired t test. **(K)** IHC showing the localizations of Pcdhgc3-mCherry in the spinal cord lamina III of P5 and P45 *Advillin^Cre^;cC3* mice and P22 *Lbx1^Cre^;cC3* mice (n = 3 animals for each genotype). **(L)** Representative images of artificial synapse formation assays in which Nrxn1β, Pcdhgc3, or Pcdhgc3^ΔEcto^ is expressed in HEK293T cells that are cocultured with neonatal spinal cord neurons. Immunolabeling for the postsynaptic excitatory synapse marker PSD95 and dendrite marker MAP2 are shown in red and magenta, respectively. Nrxn1β serves as a positive control that can induce excitatory postsynaptic specializations in spinal cord neurons. **(M)** Representative images of *in vitro* synapse formation assays in which Pcdhgc3, Pcdhga7, or GFP is expressed with in HEK 293 cells that are cocultured with neonatal spinal cord neurons. **(N and O)** Average number (N) and intensity (O) of PSD-95 puncta per HEK 293 cell, showing that Pcdhgc3, but not Pcdhgc3^ΔEcto^, Pcdhga7^GFP^, or GFP, can induce presynaptic excitatory specializations. Data are means ± SEM from three independent replicates. One-way ANOVA with Tukey’s post hoc test. ns, not significant; *p < 0.05; **p < 0.01; ***p < 0.001.

### Pcdhgc3 functions both presynaptically and postsynaptically to mediate excitatory synapse formation between sensory neurons and spinal cord neurons

Pcdhgs bind strictly homophilically when expressed in cell lines, and can form isoform-specific homophilic *trans* dimer parallel arrays to generate large two-dimensional structures bridging adjacent membranes (Brasch et al., 2019; Fernandez-Monreal et al., 2009; Frank et al., 2005; Goodman et al., 2022; Obata et al., 1995; Sano et al., 1993). Our finding that Pcdhgc3 functions in somatosensory neurons to promote excitatory synapse formation between sensory neuron axons and spinal cord neurons led us to hypothesize that Pcdhgc3 also functions in spinal cord neurons to promote postsynaptic specialization. To test this idea, *Lbx1^Cre^;Pcdhg^fcon3/fcon3^*animals were generated to excise all 22 *Pcdhgs* in dorsal horn neurons (Sieber et al., 2007). However, the size of the dorsal horn in *Lbx1^Cre^;Pcdhg^fcon3/fcon3^*animals was greatly reduced with no change in the density of synaptic markers (Figures S6A-S6D), consistent with the role of Pcdhgs, and Pcdhgc4 in particular (Garrett et al., 2019), in survival of spinal cord neurons (Prasad et al., 2008; Wang et al., 2002b). Therefore, as an alternate approach, we generated *Lbx1^Cre^;Pcdhg^C3KO/fcon3^*compound mutant animals, which harbor one *Pcdhg^fcon3^* allele and one *Pcdhgc3* null allele. Thus, in *Lbx1^Cre^;Pcdhg^C3KO/fcon3^*mice, which express Cre recombinase in dorsal horn neurons but not DRG neurons, both copies of *Pcdhgc3* are absent only in dorsal horn neurons whereas all other cells of the body are heterozygous for *Pcdhgc3* (Figure 6E). *Lbx1^Cre^;Pcdhg^C3KO/fcon3^* animals did not exhibit any gross alteration in the size of the dorsal horn (Figures S6E and S6F) but did exhibit a significant decrease in the number of dorsal horn excitatory synapses (Figure 6F). Indeed, the density of vGluT1^+^ terminals, Homer1^+^ puncta, as well as the density and size of Homer1^+^ puncta surrounding vGluT1^+^ terminals were reduced in *Lbx1^Cre^;Pcdhg^C3KO/fcon3^*mice to a similar extent as that observed in *Advillin^Cre^;Pcdhg^fcon3/fcon3^*mice (Figures 6G-J). Thus, Pcdhgc3 functions in both DRG neurons and spinal cord neurons, presumably through homophilic interactions, to promote excitatory synapse formation between sensory neurons and spinal cord neurons.

### Pcdhgc3 localizes to synapses in the dorsal horn and in a reduced preparation is sufficient to induce PSD-95 clustering in dendrites of spinal cord neurons

High resolution imaging, immunoelectron microscopy, and biochemical studies demonstrate that Pcdhgs are present in synaptic clefts and concentrated PSD fractions of synaptosomes (Fernandez-Monreal et al., 2009; Loh et al., 2016; Phillips et al., 2003; Wang et al., 2002b), yet whether Pcdhgs can directly promote postsynaptic specialization is unclear. To determine the role of Pcdhgc3 in central somatosensory neuron synapse formation, we visualized the localization of Pcdhg proteins in the central terminals of somatosensory axons by expressing Pcdhgc3 and Pcdhga1 isoforms fused to mCherry in sensory neurons using *Advillin^Cre^* (Lefebvre et al., 2012). Interestingly, while both isoforms were distributed in a punctate manner in central somatosensory neuron axons, they were distributed evenly in peripheral axons (Figures S6G and S6H). Moreover, Pcdhgc3, but not Pcdhga1, was detected in vGluT1^+^ presynaptic puncta at both P5 and in adults (Figure 6K; Figure S6G). Similarly, when expressed in dorsal horn neurons using the *Lbx1^Cre^* driver (Sieber et al., 2007), Pcdhgc3 was present in a punctate pattern, and approximately 50% of Pcdhgc3^+^ puncta were found to contain one or more Homer1^+^ puncta (Figure 6K, n = 3 animals, P20-45). These findings suggest that Pcdhgc3 can localize to both presynaptic and postsynaptic compartments in the dorsal horn.

To directly test whether Pcdhgc3 has synaptogenic activity, we next used an *in vitro* synapse formation assay (Biederer and Scheiffele, 2007; Eldeiry et al., 2017; Südhof, 2018) to ask whether Pcdhgc3 is sufficient to promote clustering of postsynaptic proteins in neonatal spinal cord neurons. Since Pcdhgc3 is expressed broadly in spinal cord dorsal horn neurons, including ∼70% of dorsal horn excitatory neurons (Figures 1I and 1J), we speculated that Pcdhgc3 could promote clustering of postsynaptic machinery in the majority of cultured dorsal horn neurons. To test this possibility, we used an *in vitro* assay in which HEK 293 cells expressing Pcdhgc3 or other Pcdhgs were co-cultured with dorsal horn neurons and the clustering of the postsynaptic protein PSD-95, an excitatory postsynaptic scaffolding protein in contacting dendrites marked by MAP2, was visualized. As a positive control for the assay, HA-tagged neurexin-1β, a known synaptogenic protein, was expressed in HEK 293 cells and found to robustly induced clusters of the PSD-95 in the dendrites of spinal cord neurons that contacted the HEK 293 cells (Figure 6L) (Graf et al., 2004; Nam and Chen, 2005). Quantification of PSD-95 clusters on dendrites in contact with transfected HEK 293 cells expressing neurexin-1β revealed a significant increase in cluster number and intensity (Figures 6N and 6O) compared to GFP-expressing HEK 293 cells.

Interestingly, Pcdhgc3 expressed in HEK 293 cells also promoted a significant increase in the number and intensity of PSD-95 clusters in spinal cord neurons, when compared to cells expressing Pcdhgc3^ΔEcto^, which lacks all 6 extracellular cadherin repeats (Pcdhgc3^ΔEcto^), or GFP (Figures 6N and 6O). Not all Pcdhgs exhibited synaptogenic activity in this assay however, because Pcdhga7-GFP, which is expressed at a lower level in the dorsal horn, did not induce clustering of PSD-95 in contacting dendrites (Figures 6M-6O). These findings are consistent with the *in vivo* gain-of-function and loss-of-function experiments and suggest that Pcdhgc3 is sufficient to induce excitatory postsynaptic specialization in spinal cord neurons.

### Altered spinal cord dorsal horn response to touch in response to reduced mechanosensory inputs

The behavioral alterations and reductions in both LTMR central synapses and peripheral axonal branching in *Advillin^Cre^;Pcdhg^fcon3/fcon3^*mice raise the question about how changes in mechanosensory synaptic inputs to the dorsal horn affect dorsal horn somatosensory circuit function in the mutants. To address this question, *in vivo* multielectrode array recordings of the spinal cord dorsal horn were performed to assess dorsal horn neuron responses to a series of mechanical step indentations applied to hindpaw glabrous skin (Figure 7A). Interestingly, the baseline firing rates of dorsal horn units were significantly increased in *Advillin^Cre^;Pcdhg^fcon3/fcon3^* mice (Figure 7B). When step indentations were applied to the skin, dorsal horn neurons in control animals exhibited responses at both the onset and offset of the step indentations within the low force range, beginning around 5 mN (Figure 7C). Dorsal horn neurons also showed responses during the sustained phase of the step indentation, starting around 10 mN. In comparison, dorsal horn neurons in *Advillin^Cre^;Pcdhg^fcon3/fcon3^* mice exhibited reduced responses to both the onset and offset of step indentation, but not during the sustained phase of the step indentation (Figures 7D, 7E and S7A). Using a grid of indentation sites to map functional receptive fields of dorsal horn neurons (Figure 7F), dorsal horn units recorded in *Advillin^Cre^;Pcdhg^fcon3/fcon3^*mice showed significantly smaller receptive fields (Figure 7G). Taken together, the *Pcdhg* mutant animals have fewer synaptic inputs to the spinal cord dorsal horn, which results in an increase in tonic firing in dorsal horn neurons, reduced firing in response to mechanical stimuli acting on the skin, and smaller functional receptive fields.

**Figure 7.**
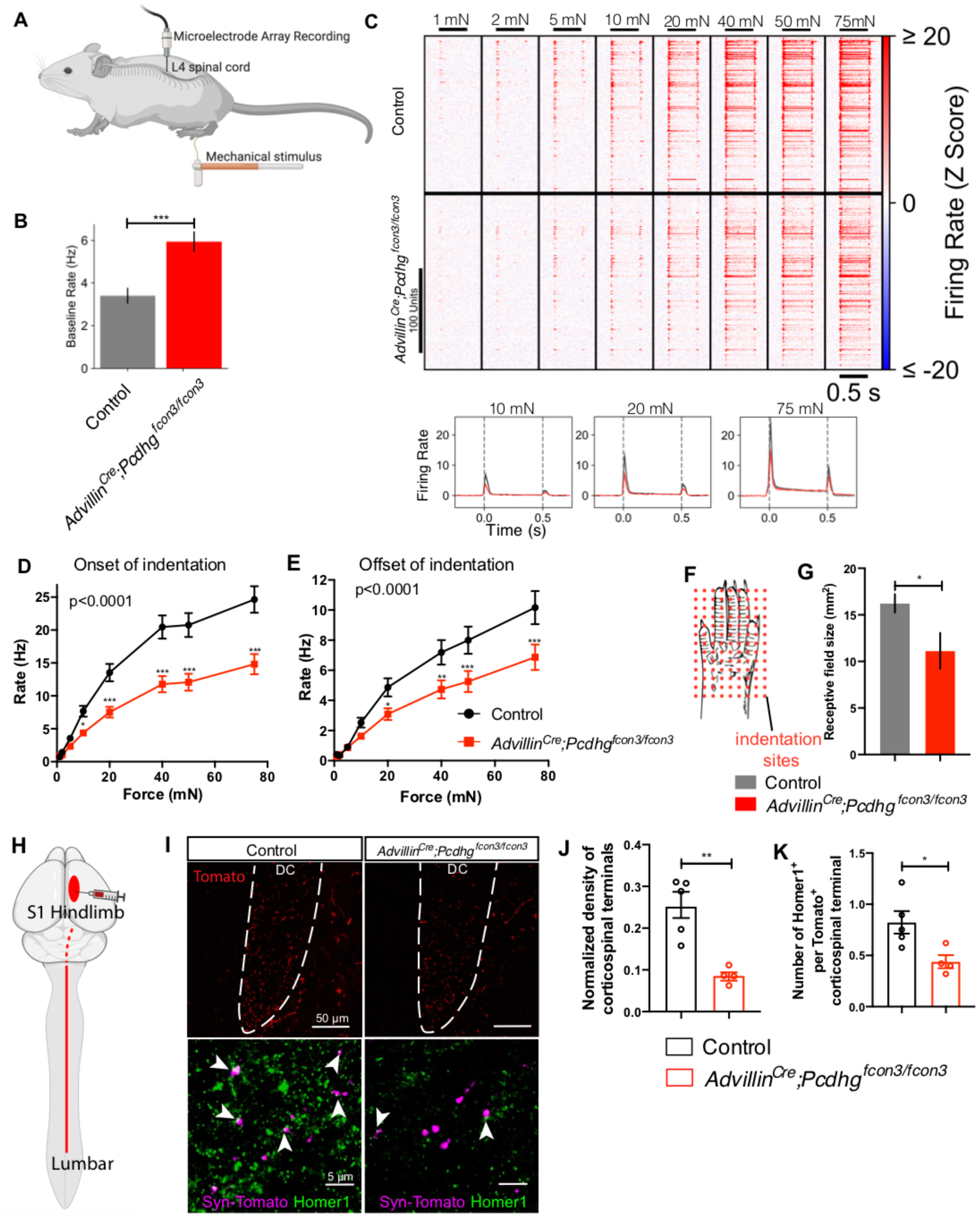
Aberrant physiological responses and corticospinal synaptic inputs in the dorsal horn of *Advillin^Cre^; Pcdhg^fcon3/fcon3^* mice due to reduced functional mechanosensory synapses. **(A)** MEA recordings from lumbar spinal cord dorsal horn in mice, while the glabrous skin from right hind paw is indented with a series of forces. **(B)** Quantifications of the baseline firing rates in littermate control and *Advillin^Cre^;Pcdhg^fcon3/fcon3^* mice. n = 3 animals per genotype. Student’s unpaired t test. **(C)** Summary of representative units responding to indentation of the skin with different forces. The amount of force is labeled on the top. The indentations applied are indicated by the black bars below the force labels. Average firing rates in response to the indentation are shown on the bottom row. n = 187 neurons for 5 controls and n = 207 neurons for 5 mutants. **(D and E)** Quantifications of the onset (D) and offset (E) responses in littermate control and *Advillin^Cre^;Pcdhg^fcon3/fcon3^* mice. Two-way ANOVA. **(F)** Diagram for mapping the functional receptive fields of the dorsal horn neurons. **(G)** Quantifications of the receptive field sizes in littermate control and *Advillin^Cre^;Pcdhg^fcon3/fcon3^* mice. **(H)** Diagram showing viral labeling of corticospinal projections in the lumbar spinal cord. **(I)** IHC images showing corticospinal terminals labeled with synaptophysin-tdTomato in the dorsal column, and IHC images showing corticospinal terminals colabeled with Homer1 in control and *Advillin^Cre^;Pcdhg^fcon3/fcon3^*mice. White dotted line outlines the shape of the dorsal column with labeling. Arrowheads point to the Tomato^+^ puncta with Homer1 labeling. **(J)** Quantification showing the density of Tomato^+^ corticospinal terminals in lamina III is reduced in the *Advillin^Cre^;Pcdhg^fcon3/fcon3^* mice (number of puncta per 10^4^ μm^2^). Each dot represents average number for an animal. Student’s unpaired t test. **(K)** Quantification showing the number of excitatory synapses (Homer1^+^ puncta) surrounding corticospinal terminals in lamina III is reduced in the *Advillin^Cre^;Pcdhg^fcon3/fcon3^* mice. Student’s unpaired t test. *p < 0.05; **p < 0.01; ***p < 0.001.

### Reduced corticospinal input to the dorsal horn is secondary to reduced mechanosensory inputs

The altered tonic and evoked firing patterns in the dorsal horn prompted us to ask whether loss of *Pcdhgs* in sensory neurons affects other aspects of dorsal horn mechanosensory circuit assembly. Indeed, our genetic analyses had suggested that the decrease of vGluT1^+^ terminals in *Advillin^Cre^;Pcdhg^fcon3/fcon3^*mice may reflect changes in the corticospinal inputs in lamina III, as the density of vGluT1^+^ somatosensory terminals remained unchanged in these mutants (Figures 3H). To directly test this possibility, AAV2-retro-hSyn-synaptophysin-tdTomato virus was injected into the hindlimb region of primary somatosensory cortex (S1) of control and *Advillin^Cre^;Pcdhg^fcon3/fcon3^*mice to visualize corticospinal synapses in the spinal cord dorsal horn (Figure 7H). Viral injections labeled comparable numbers of axons in the corticospinal tract in control (122.1 ± 11.3 axon per dorsal column section) and *Advillin^Cre^;Pcdhg^fcon3/fcon3^*mice (140.9 ± 5.9 axon per dorsal column section, p > 0.05, Figure 7I). However, the density of corticospinal terminals in medial lamina III, measured in the same region in which the vGluT1 and Homer1 synaptic analyses were done, was significantly reduced in *Advillin^Cre^;Pcdhg^fcon3/fcon3^* mice (Figure 7J). Consistent with the finding that Homer1^+^ puncta density is decreased throughout lamina III, the number of Homer1^+^ synapses associated with Tomato^+^ corticospinal terminals was reduced in the mutants (Figure 7J). Finally, we asked whether the absence of synaptic structures, per se, or the absence of synaptically-evoked activity in the dorsal horn accounts for the deficits in corticospinal inputs to lamina III observed in *Advillin^Cre^;Pcdhg^fcon3/fcon3^*mice. Thus, we tested whether silencing touch-evoked activity in somatosensory neurons leads to a deficit in corticospinal neuron innervation of lamina III, in a similar manner. For this we used *Cdx2^Cre^; Piezo2^f/f^*mice, in which Piezo2 the principal mechano-sensitive ion channel for LTMRs (Ranade et al., 2014) was deleted broadly in neurons below cervical level C2, including DRG neurons. *Cdx2^Cre^; Piezo2^f/f^* mice lack most if not all low-threshold, mechanically evoked responses in DRG neurons (Lehnert et al., 2021). Similar to the *Advillin^Cre^;Pcdhg^fcon3/fcon3^*mutant mice, *Cdx2^Cre^;Piezo2^flox/flox^* mutants exhibited a reduction in the density of corticospinal neuron terminals as well as their synapses in dorsal horn lamina III (Figures S7B-S7D). These findings suggest that the deficit in corticospinal synapses in the dorsal horn of *Advillin^Cre^;Pcdhg^fcon3/fcon3^* mutant mice is caused by a reduction in mechanically-evoked sensory neuron synaptic drive in the dorsal horn.

## Discussion

Here, we report unique functions of Pcdhg isoforms across different subcellular compartments of somatosensory neurons for the assembly of the somatosensory circuitry and behavioral reactivity to touch. Pcdhgc3 mediates homophilic interactions between sensory neuron presynaptic terminals and spinal cord neurons to promote synapse formation. In the periphery, one or more other *Pcdhg* family members promotes proper axonal branching patterns through mediating interactions between somatosensory neuron axons and Schwann cells. Functionally, Pcdhgs in somatosensory neurons are essential for both tonic and touch evoked physiological responses in dorsal horn neurons, behavioral responses to touch and sensorimotor control, and the development of corticospinal synapses in the spinal cord. Therefore, spatially coordinated patterns of expression of three or more clustered protocadherin gamma isoforms underlie the formation of synapses between LTMRs and dorsal horn neurons (Pcdhgc3), peripheral axonal branching (possibly Pcdhga7 and others), dorsal horn neuron survival (Pcdhgc4) (Wang et al., 2002b), corticospinal synapses with dorsal horn neurons (Pcdhgc3, non-cell autonomously), and thus step-wise assembly of the dorsal horn somatosensory circuitry.

Several lines of evidence have pointed to clustered protocadherins as potentially important molecular players in synapse development, function, or maintenance in the mammalian nervous system. Indeed, Pcdhgs are found at synaptic clefts and enriched in postsynaptic density (PSD) fractions of synaptosomes (Loh et al., 2016; Phillips et al., 2003; Wang et al., 2002b; Weiner et al., 2005) and their presence is associated with synapse maturation (LaMassa et al., 2021; Li et al., 2010; Phillips et al., 2003). Moreover, many synapse-associated molecules are found in Pcdhg protein complexes, including PSD-95, α-catenin, and neuroligin-1 (Han et al., 2010; Molumby et al., 2017), suggesting that Pcdhgs may directly control or modulate synapse assembly or composition. Despite this evidence, the roles of Pcdhgs in synapse formation and function have remained enigmatic (Missler et al., 2012; Südhof, 2017), in part due to their requirements in other developmental processes, including neuronal survival and dendrite elaboration (Flaherty and Maniatis, 2020; Garrett et al., 2012; 2019; Kostadinov and Sanes, 2015; Lefebvre et al., 2008; Molumby et al., 2016; Wang et al., 2002b; Weiner and Jontes, 2013; Weiner et al., 2005). Indeed, in the spinal cord, removing all *Pcdhgs* leads to loss of spinal cord neurons (Wang et al., 2002b). Although a reduction in synaptic density was observed when inhibiting apoptosis in the spinal cord of *Pcdhg* mutants (Weiner et al., 2005), it had remained unclear whether these synapse alterations were a reflection of Pcdhg functions in other developmental processes, such as astrocyte-neuron contacts or dendrite development (Garrett and Weiner, 2009; Garrett et al., 2012; Molumby et al., 2016; 2017). In the present study, we took advantage of the dispensability of Pcdhgs for DRG neuron survival (Prasad and Weiner, 2011) to determine their roles in somatosensory neuron synapse formation. Both *in vivo* and *in vitro* evidence supports a model in which Pcdhgc3 in sensory neurons and dorsal horn neurons acts homophilically to promote postsynaptic specialization.

We propose that Pcdhgc3 functions as a cell-adhesion molecule that tethers the pre- and postsynaptic membranes and instructs postsynaptic specialization through its unique intracellular domain, encoded by its variable exon, and association with postsynaptic proteins. Given the ability of Pcdhgs to form large two-dimensional parallel arrays between opposing membranes (Brasch et al., 2019), Pcdhgc3 may also coordinate the proper alignment or spacing of pre- and postsynaptic membranes. The unique variable juxtamembrane domain of the Pcdhgc3 intracellular region may induce specific downstream signaling, as illustrated by its binding to Axin1 and thus modulation of Wnt signaling (Mah et al., 2016). Future work will define the contributions of the unique Pcdhgc3 intracellular sequence in recruiting postsynaptic scaffolding proteins, such as PSD-95, and promoting postsynaptic specialization and synaptic maintenance. While the present work reveals a synaptogenic role for Pcdhgc3 in somatosensory neurons, other Pcdhg isoforms are widely expressed throughout the developing nervous system with extraordinarily complex patterns. Thus, it will be interesting to determine whether Pcdhgc3 and other isoforms promote synapse formation between neurons that express the same isoforms, thereby contributing to a synaptic specificity code for nervous system wiring (Canzio and Maniatis, 2019).

Our genetic analyses of Pcdhg isoforms show that different isoforms act in the same neuron to mediate synapse formation and peripheral axonal branching. These findings highlight a remarkable compartmentalization of Pcdhg function via homophilic interactions in the peripheral and central axonal branches of somatosensory neurons for distinct developmental processes. This compartmentalized signaling could be achieved by local differences in the proteins that interact with Pcdhgs to form unique membrane complexes. In the developing skin, nascent sensory axons are closely associated with peripheral Schwann cells (Jessen et al., 2015). Thus, neuron-glia interactions mediated by Pcdhgs may physically stabilize nascent sensory axon branches or activate downstream signals that directly promote axonal branching. It is possible that one particular isoform is required for peripheral axonal branching, such as the most highly expressed isoform Pcdhga7. Another possibility is that a critical total level of any Pcdhg isoform is sufficient, and the common Pcdhg intracellular region may signal through a pathway that promotes arborization, such as the FAK/PKC/MARCKS pathway which contributes to cortical dendrite arborization (Garrett et al., 2012). Future genetic and molecular dissection of Pcdhg isoforms and their intracellular signaling mechanisms as well as isoform expression patterns in developing glia and neurons will provide insights into Pcdhg function in peripheral axonal branching.

Our findings also show that the sensory neuron clustered protocadherins, presumably Pcdhgc3, are crucial for corticospinal synaptic inputs of the dorsal horn. We consider this to be a non-cell autonomous function of Pcdhgc3 because corticospinal synapses form onto spinal cord neurons and not onto the axon terminals of primary sensory neurons that lacked Pcdhgs in the conditional mutants (Hanaway and Smith, 1979; Valtschanoff et al., 1993). Moreover, deficits of corticospinal inputs to the dorsal horn in *Advillin^Cre^;Pcdhg^fcon3/fcon3^* mutant mice were similarly observed in *Piezo2* conditional knockout mice, which lack mechanically evoked synaptic drive of dorsal horn neurons. Together, these findings support a model in which functional synapses between sensory neurons and dorsal horn neurons and sensory-evoked synaptic drive in the dorsal horn are a prerequisite for the formation of corticospinal synapses onto dorsal horn neurons. Spontaneous and evoked sensory activity play essential roles in wiring the nervous system (Ebert and Greenberg, 2013; Kirkby et al., 2013; Leighton and Lohmann, 2016), although in the mechanosensory system less is known. Postnatal functional development of A-fiber sensory connectivity in the dorsal horn requires NMDA receptor activation, suggesting that the precise spinal cord wiring is activity-dependent (Beggs et al., 2002). In the rat, LTMRs exhibit spontaneous activity only during embryonic development, while evoked responses are detected beginning at late embryonic stages (Fitzgerald, 1987b; 1987a). Central deficits in somatosensory neuron *Pcdhg* conditional knockout animals appeared later than P0, suggesting that deficits in evoked synaptic activity, rather than spontaneous synaptic activity, is responsible for the alterations in corticospinal synaptic input observed in the mutants. Consistent with this, corticospinal innervation of the lumbar spinal cord (the area analyzed in the present study) begins ∼P4 (Hsu et al., 2006), suggesting that postnatal evoked synaptic activity regulates formation of synapses between nascent corticospinal axon terminals and spinal cord interneurons. The reduction in corticospinal input may decrease descending inhibition, therefore causing an increase in baseline firing rate observed in the mutant dorsal horns. Thus, the widespread deficits in corticospinal synapses in the dorsal horn suggest an obligatory role for postnatal Pcdhgc3-dependent touch evoked synaptic transmission in the assembly of dorsal horn mechanosensory circuitry. Future studies should aim to determine how mechanosensory synaptic transmission in the dorsal horn instructs this assembly.

## Acknowledgements

We thank members of the Ginty lab for discussions and feedback on the manuscript. We thank Josef Turecek for help with electrophysiology, the Harvard Medical School Neurobiology Imaging Facility for assistance with the small-RNA FISH study, and the Harvard Medical School Electron Microscopy Facility for help with the electron microscopy experiments. We thank Annie Chen for genotyping and mouse breeding. This work was supported by a Hanna Gray Fellowship (S.M.), NSF GRFP fellowship (E.H.), NIH 1R01 AT011447-01 (D.D.G.), NIH R35 5R35NS097344-05 (D.D.G.), the Edward R. and Anne G. Lefler Center for Neurodegenerative Disorders (D.D.G.), NIH R01 NS055272 (J.A.W.), NIH R01 EY031690 (A.M.G.), NIMH R01 5R01MH108579 (T.M.), and Canada Research Chair and CIHR PJT-148961 (J.L.L.). D.D.G. is an investigator of the Howard Hughes Medical Institute. This article is subject to HHMI’s Open Access to Publications policy. HHMI lab heads have previously granted a nonexclusive CC BY 4.0 license to the public and a sublicensable license to HHMI in their research articles. Pursuant to those licenses, the author-accepted manuscript of this article can be made freely available under a CC BY 4.0 license immediately upon publication.

## Author Contributions

S.M. and D.D.G. conceived the study. S.M. and B.K. performed RNA sequencing analysis. S.M., K.C., M.D., and A.T. performed behavioral experiments. S.M., K.C., E.O.A., and E.H. performed anatomy and artificial synapse experiments. A.C. performed *in vivo* spinal cord MEA recording experiments. E. H. performed stereotactic injections. Q.Z., K.C. and S.M. performed electron microscopy experiments. A.M.G., L.C.F., and J.A.W. generated and characterized the *Pcdhg* mutants using the CRISPR/Cas9 technique and provided mutant mouse tissues for analyses. E.K.F. and T.M. provided tissues from *Pcdhg^TAKO^* mutants. J.L.L. provided *Pcdhg^fcon3^* mice. S.M. and D.D.G. wrote the paper, with input from all authors.

## STAR METHODS

### EXPERIMENTAL MODEL AND SUBJECT DETAILS

All experiments were conducted in accordance with Harvard Medical School (Institutional Animal Care and Use Committee) IACUC protocols. Animals used in the study were group housed on a 12-hour light/dark cycle, with control and mutant animals in the same litters and cages. Littermates from the same genetic crosses were used as controls, to control for variability in mouse strains/backgrounds. No differences were observed between wildtype animals and floxed (*Pcdhg^fcon3/+^* and *Pcdhg^fcon3/fcon3^*) or Cre littermate control groups for the experiments performed. Wildtype C57BL/6J mice were used for the comparison among various *Pcdhg* deletion mutants.

#### Mouse Lines

*Pcdhg* lines used in this study include *Pcdhg^fcon3^* (Prasad et al., 2008), *Pcdhg^C3KO^*(Garrett, et al., 2019), *R26^LSL-Pcdhga1-mCherry^* (cA1-mCherry), and *R26^LSL-Pcdhgc3-mCherry^* (cC3-mCherry) (Lefebvre, et al., 2012), *Pcdhg^1R1^*, *Pcdhg^3R1^*, and *Pcdhg^3R2^* (Garrett, et al., 2019), and *Pcdhg^TAKO^*(Chen et al., 2012) were previously described. Pcdhg mutant mice were maintained on a C57/B6J background. Other mouse lines used include *R26^LSL-synaptophysin-tdTomato^*(Ai34) (Jax#012570); *R26^LSL-tdTomato^* (Ai14) (Jax# 007908); *Advillin^Cre^* (Hasegawa et al., 2007); *Advillin^CreERT2^*(Lau et al., 2011); *Lbx1^Cre^* (Sieber et al., 2007); *Dhh^Cre^*(Jaegle et al., 2003); *Cdx2^Cre^* (Coutaud and Pilon, 2013); *Piezo2^f/f^* (Woo et al., 2014); *Rpl22^HA^* (Sanz et al., 2009); *PV^CreER^* (Hippenmeyer et al., 2005); *TrkB^CreER^* (Rutlin et al., 2014) and *Ret^CreER^* (Luo et al., 2009). Male and female mice of C57BL/6J and CD1 backgrounds were used for these studies. Expression of GFP in the *Pcdhg^fcon3^*allele was assessed using histological verification.

### METHOD DETAILS

#### Tissue Fixation

Male and female mice of ages younger than 6 days were anesthetized in ice until unconscious, then sacrificed quickly by decapitation. Male and female mice of ages 6 days and older were anesthetized with isofluorane and a transcardial perfusion was performed with 10-15 mL modified Ames Media with heparin, followed by 15-25 mL 4% PFA. Back hairy skin was dissected following application of Nair commercial depilatory cream and consequent hair removal, and the underside of the skin was scrapped clean of fatty tissues and muscles. Back hairy skin and glabrous skin of the paws were dissected and post-fixed in Zamboni’s Fixative overnight at 4°C. Spinal cords and DRGs were post-fixed in 4% PFA overnight at 4°C. Following post-fixation, samples were washed in 1xPBS for 3-5 hours and stored in PBS + 0.01% sodium azide at 4°C for long-term storage.

#### Tamoxifen Treatment

Tamoxifen (Sigma, Cat# T5648-1g) was dissolved in 100% ethanol (20 mg/ml) on a rotator for 10 minutes, aliquoted into Eppendorf tubes and mixed with an equal volume of sunflower seed oil (Sigma, Cat# s5007). The mixture was vortexed for 15 minutes and centrifuged in a pressurized vacuum concentrator for 45 minutes to remove the ethanol. When removed from the vacuum concentrator, all tubes were checked carefully to ensure only sunflower seed oil remained, then stored at -20°C for long-term use. To calculate embryonic timepoint, females were checked every morning to monitor when mating occurred. The tamoxifen solution was delivered to pregnant females via oral gavage for embryonic tamoxifen treatment at the timepoints indicated. For all analyses, the morning after coitus was designated as E0.5 and the day of birth as P0. Tamoxifen was administered to control and littermate mice in order to excise the *Pcdhg^fcon3^* alleles using *Advillin^CreERT2^* in peripheral somatosensory neurons. Intraperitoneal injections of tamoxifen (1 mg per day) were administered to mice for 5 consecutive days, from either P15-P19. All mice in this *Advillin^CreERT2^*experiments received this tamoxifen regimen, and no changes in health or behaviors were observed in controls or mutant mice.

#### Immunohistochemistry

Spinal cord, DRG, and finger pad samples were cryoprotected in 30% sucrose in PBS overnight at 4°C. Samples were then rinsed in PBS, embedded in OCT in tissue molds over a 100% ethanol and dry ice bath, and stored at -80°C for at least an hour before cryosectioning. 20-25 μm sections were produced for spinal cord, DRG, and brainstem, and 30 μm transverse sections for fingertip sections. Mutant and littermate control sections were collected on the same slide. Slides were allowed to dry on the benchtop overnight and stored at - 20°C.

Slides were defrosted at room temperature for at least 30 minutes, then a hydrophobic barrier was applied with a PAP pen (H-4000, VWR/Vector Lab) around the samples. Slides were then rehydrated with filtered 1xPBS for 3x5 minutes, blocked for 1 hour at room temperature with 1xPBS with 0.1% Triton X-100 and 5% normal goat or donkey serum (Jackson Immuno, 005-000-121), and incubated with primary antibodies in blocking solution (5% serum without detergent) overnight at 4°C. Slides were washed 4x5 minutes with 1xPBS with 0.02% Tween-20, incubated with secondary antibodies in blocking solution (5% serum without detergent) at room temperature for 2 hours, washed 4x5 minutes with 1xPBS with 0.02% Tween-20 detergent, and mounted. Samples were mounted with fluoromount-G (Southern Biotech) or DAPI Fluoromount-G® (Southern Biotech) when visualization of nucleus is needed. All slides were stored at 4°C for up to 1 week until imaged.

The following primary antibodies were used: guinea pig anti-vGluT1 (Millipore Sigma, RRID: AB_2301751; 1:1000), rabbit anti-Homer1 (Synaptic Systems, Cat# 160003, 1:1000), Isolectin B4 647 (Invitrogen, RRID: SCR_014365, 1:500), chicken anti-NFH (Aves, Cat#NFH0211, 1:500); rabbit anti-DsRed (Clontech, RRID: AB_10013483, 1:500), goat anti-mCherry (Sicgen, RRID: AB_2333092, 1:500), goat anti-GFP (Abcam, RRID: AB_305635, 1:500), mouse anti-NeuN (Millipore, RRID: AB_2298772, 1:500), rabbit anti-S100 (Proteintech, Cat# 15146-1-AP, RRID:AB_2254244, 1:500) and rat anti-Troma1 (DSHB, RRID: AB_531826, 1:50). Secondary antibodies include Alexa 405, 488, 546 or 647 conjugated donkey or goat anti-mouse, rabbit, chicken, goat or guinea pig, and used at a 1:500 dilution (Life Technologies or Jackson ImmunoResearch).

#### Wholemount hairy skin immunostaining

Back hairy skin was dissected, and fat tissue under the skin was scraped off with a stainless-steel spatula. The skin was then fixed in Zamboni fixative (Fisher/VWR, NC9335034) at 4°C overnight. The tissue was rinsed in PBS three times and then washed with PBS containing 0.3% Triton X-100 (0.3% PBST) every 30 min for 5 to 8 hours. Then, the skin was incubated with primary antibodies in 0.3% PBST containing 5% goat or donkey serum and 20% DMSO at room temperature for 3 to 4 days. Skins were then washed with 0.3% PBST every 30-60 min for 5 to 8 hours. The tissues were transferred to secondary antibodies in 0.3% PBST containing 5% goat/donkey serum and 20% DMSO and incubated at room temperature for 2 to 3 days. The tissues were then washed with 0.3% PBST every 30-60 min for 5 to 8 hours. The skins were dehydrated in 50% methanol for 15 min, 80% methanol for 15 min, and 100% methanol for 15 min, and 100 % methanol overnight. For imaging, skins were cleared in BABB (Benzyl Alcohol, sigma 402834; Benzyl Benzoate, sigma B-6630; 1:2) at room temperature for 10 min.

#### RNA sequencing and data analysis

DRGs from P3 pups were dissected and RiboTag immunoprecipitation was performed as previously described (Sanz et al., 2009). DRGs from the thoracic levels of five pups were combined and subject to immunoprecipitation to obtain mRNA for each sequencing replicate, and three to five biological replicates were sequenced for each group of neurons. Briefly, pups were sacrificed and DRGs were dissected on ice in DMEM Complete Medium with 1% PenStrep (Cat# 15140122, Thermo Fisher Scientific). DRGs were snap-frozen and stored at -80°C. Samples were homogenized (homogenization buffer with 5mg/ml cycloheximide, cOmplete protease inhibitor cocktails, 100mg/ml heparin, RNAsin and 1M Dithiothreitol) and centrifuged at 10,000 rpm at 4 °C for 10min. Lysis buffer (RLT buffer from RNeasy kit, Cat# 74004, Qiangen) was added to each sample, and 800 μl were collected from each spun-down lysate. 5 μl of Mouse anti-HA antibody (Cat#H9658, Sigma-Aldrich), was added to each sample and all samples were incubated for 4 hours at 4 °C on a gentle spinner. 200μl of Dynabeads™ Protein G (Cat#10004D, Thermo Fisher) were equilibrated with 1000 μl of homogenization buffer (RNase-free water with 10% NP-40, 1M KCl, 1M MgCl_2_, 1.5M Tris, pH 7.4) for 10 min at 4 °C. Homogenization buffer was then suctioned off and antibody-tissue homogenate sample was added to the beads and incubated overnight at 4 °C on a gentle spinner/rotator. Next day, beads were washed with 800 μl of high salt buffer for 3 X 10 min at 4C in cold room on rotator. 350 μl of lysis buffer (RLT lysis buffer with βME) were added to each sample after removing final high salt wash buffer and standard RNA purification was performed using the manufacturer’s protocol (RNeasy kit, Cat# 74004, Qiangen).

Purified RNA samples with a concentration of 1-10 ng/μL for 8-10 μL and RNA integrity number (RIN) above 8 (except for one sample for Aβ SAI-LTMRs that had a RIN ∼7) were used for library preparation and RNA-sequencing at the Biopolymers Facility at Harvard Medical School. Libraries were prepared using Illumina NexteraXT using the manufacturer’s protocol. The libraries were pooled together and then sequenced using the Illumina NextSeq 500 sequencer with 150 bp paired-end reads. Reads were aligned and counted using STAR (Dobin et al., 2013). Both read count matrix and normalized data using transcript per million reads (TPM) were obtained. The matrix of read counts was transformed using a regularized logarithm (rlog) implemented in DESeq2 (Love et al., 2014). The heatmaps and dotplots were plotted using the ggplot2 package in R. Subtype uniquely enriched genes and differentially expressed genes analyses were performed as previously described (Zheng et al., 2019). Briefly, genes were then grouped based on the neuronal groups their expression is enriched in and ordered based on the enrichment. Genes with rlog values in the top 75% and displayed expression differences over 1 in all replicates were used for displaying the expression pattern. Differentially expressed genes (DEGs) were defined using the criteria of a 1.5-fold change with a false discovery rate (FDR) <0.01. Subtype uniquely enriched genes (SUEGs) were selected for genes that are enriched in each group based on pairwise comparison, thus the FDR for SUEGs after correction for multiple comparisons is ∼0.04. Rlog difference from the average rlog value was plotted in the main heatmaps using a scale of -2 to 2 and average rlog values were plotted in another heatmap to the right of the main heatmap.

#### Quantification of Meissner corpuscle density

Images of serial sections of glabrous pads from forepaw fingertips were sectioned at 30 μm. Images of sections were analyzed using ImageJ. In each section, the number of Meissner corpuscles, visualized by using S100 antibody (Meissner corpuscles form ovoid masses in the dermal papillae and Schwann cells surrounding axons are S100+) and NFH antibody. Then, the area of the epidermis was traced and measured in ImageJ. Density was calculated by dividing the number of corpuscles by the area of epidermis.

#### Dorsal column injections

Mice (8-12 weeks old) were anesthetized via continuous inhalation of isoflurane (2%) from a precision vaporizer for the 20-40 minutes duration of the surgery. Protective eye ointment (Puralube) was applied to the eyes. The animal’s breathing rate was monitored throughout the procedure and the isoflurane dose was adjusted as necessary. Hair from the back of the neck was treated by a commercial hair remover (Nair, Church and Dwight Co.; Princeton, NJ) for 1-3 min, and swabbed with water, Betadine, and 70% ethanol. A 7 mm incision was made in the midline of the skin at the cervical level. Muscles were separated from the midline until the spinal cord cervical vertebrae were exposed. A small incision was made on the dura and arachnoid membranes above the C1 cervical spinal vertebrae to expose the dorsal column. Next, 100 nl of AAV-Retro-Flex-PLAP (obtained from Dr. Sara Prescott from Harvard Medical School) was injected into the dorsal column above the C1 level using a glass pipette under visual guidance. Afterwards, muscles and skin were stitched together with sutures, and SR Buprenorphine (1.5 mg/kg) was applied for analgesia. Mice recovered from anesthesia on a warm pad for ∼1 hour and then returned to their home cage. The condition of the mice, including locomotor activity, the healing of wounds, body weight, and grooming, were monitored daily. 3-4 weeks after surgery, mice were sacrificed by CO2 asphyxiation followed by perfusion.

#### Alkaline phosphatase staining

AAV-Retro-Flex-PLAP virus was injected into the dorsal column on littermate control and mutant animals. 3-5 weeks following the injection, animal was perfused and back hairy skins were treated with commercial hair remover and rinsed with PBS. Skins were further dissected and carefully scrapped free of remaining fat tissue under the skin. Spinal cords and DRGs were dissected. Samples in PBS were placed in a 65-68°C incubator for 2 hours, washed 3x5 minutes in B3 buffer (0.1M Tris pH 9.5; 0.1M NaCl; 50mM MgCl_2_; 0.1% Tween-20), and incubated at room temperature for 24 hours in B3 buffer with 3.4 μL/mL each of NBT (Nitro Blue Tetrazolium, Roche 11-383-213-001) and BCIP (5-Bromo-4-chloro-3-indolyl-phosphate, 4-toluidine salt, Roche 11-383-221-001). Samples were then incubated with 4% PFA for 1 hour at room temperature, washed with 1xPBS, and dehydrated for ∼30 minutes with 50%, 75%, and 100% ethanol overnight. Samples were stored in 100% ethanol at -20°C until imaging. For imaging, benzyl alcohol and benzyl benzoate mixed 1:2 (BABB) was used to clear tissue for imaging. Samples were cleared with BABB on a rotator for 10 minutes before imaging.

#### Corticospinal neuron labeling

To label corticospinal neurons, mice underwent unilateral stereotactic injection of rAAV2-retro-CAG-tdt virus (Penn viral core) into the hindlimb region of primary somatosensory cortex (S1). Mice were injected with SR Buprenorphine (0.1 mg/kg), anesthetized with isoflurane and placed in a small animal stereotaxic frame (David Kopf Instruments). The skull was exposed under aseptic conditions and one craniotomy was made centered at -0.6 posterior and 1.75 lateral from bregma. A borosilicate glass pipette was lowered to a depth of 0.7 mm from the surface of the brain. 150 nL of virus was injected at a rate of 50 nL·min−1 using a Microinject system (World Precision Instruments). 4 weeks following the injection, animals were perfused and spinal cords were collected for immunohistochemistry.

#### RNAScope

P3 animals were anesthetized in ice for 2-4 minutes or until unconscious, then sacrificed quickly by decapitation. P21 mice were euthanized by CO_2_ followed by decapitation. DRGs and spinal cords from mice were rapidly dissected and the axial level was identified using the T13 DRG as a landmark. DRG were quickly frozen in dry-ice-cooled 2-metylbutane and stored at −80 °C. DRG were sectioned at a thickness of 15 μm and RNAs were detected by RNAscope (Advanced Cell Diagnostics) using the manufacturer’s protocol. The following probes were used: mm-Pcdhga2 (Cat# 2835811), mm-Pcdhga7-C2 (Cat# 835781-C2), Mm-Pcdhgb1-O1 (Cat# 837621), Mm-Pcdhgc3(Cat# 802841), Mm-Pcdhgc4-C3 (Cat# 835791-C3), Mm-Slc17a6-C3 (Cat# 319171-C3), Mm-Nefh (Cat# 443671), and Mm-Plp1 (Cat# 428181).

#### Electron microscopy and image analysis

Electron microscopy of the spinal cord was conducted as previously described (Zhang et al., 2019). Briefly, animals were transcardially perfused and fixed with 2% PFA (Electron Microscopy Sciences) and 2.5% glutaraldehyde (Electron Microscopy Sciences), and postfixed in the same fixative overnight. Spinal cords were dissected out and sectioned at 200 μm with a Leica VT1000 S vibratome. These sections were then osmicated with reduced osmium (1% osmium tetroxide (Electron Microscopy Sciences)/1.5% potassium ferrocyanide (MilliporeSigma)), stained with 1% uranyl acetate (Electron Microscopy Sciences), dehydrated with an ethanol series followed by propylene oxide, and infiltrated and embedded with an epoxy resin mix (LX-112, Ladd Research). Samples were then cured in the oven at 60 °C for 48-72 hours. Samples were sectioned using a Leica EM UC7 ultramicrotome with Diatome diamond knives, and ultrathin sections (40-60 nm) were picked up on glow discharged formvar/carbon films on slot grids (Ted Pella). Grids were post stained with 2% uranyl acetate in 50% acetone and 0.2% lead citrate (Electron Microscopy Sciences) before imaging. Ultrathin sections were imaged using a JEOL 1200EX transmission electron microscope at 80 kV accelerating voltage and 10,000× nominal magnification with an AMT XR- 611 CCD camera at a final pixel size of 1.84 nm.

Micrographs of the glomeruli were taken in lamina III of the spinal cord, based on morphological characteristics of glomeruli previously described (Ribeiro-da-Silva et al., 1985). Experimenter was blind to the genotypes. For quantification of the micrographs, images were adjusted with normalization using Fiji/ImageJ to enhance contrast. Morphometric classification of synapses and analysis of ultrastructural PSD parameters were performed as previously described (Robbins et al., 2010). Quantifications of the length and thickness of PSDs were performed with experimenter blind to the genotypes.

#### Synaptic puncta analysis

Z stack images for puncta analysis were obtained on the Zeiss LSM700 confocal microscope with the 63x objective. Images were taken in lamina III, within 150 μm of lamina IIiv marked by IB4. Imaging parameters were held constant for littermate control and mutants, which were immunostained on the same glass slide to reduce variation. At least 3 animals of each genotype or age were used, with at least ∼2000 vGluT1 and ∼8000 Homer1 puncta per animal. For each animal used in analysis, 4-6 sets of images, each image set comprising (4 to 5) 1 μm z stacks from a minimum of 3 separate spinal cord sections, was used for analysis. Homer and vGluT1 puncta analyses were performed using a custom script in ImageJ, creating a mask of vGluT1 terminals between 0.5 μm and 5 μm in diameter, thresholding Homer1, and counting puncta of 0.1 μm to 5 μm in diameter contained within that mask.

#### Whole-cell patch-clamp recordings using acute spinal cord slices

Mice (P13-P16) were anesthetized via continuous inhalation of isoflurane (2%) and then perfused with ice-cold choline solution. Lumbar enlargements (L1-L5) were dissected out from vertebral columns in ice-cold choline solution (in mM: 110 Choline Chloride, 2.5 KCl, 1.25 NaH_2_PO_4_, 25 NaHCO_3_, 25 Glucose, 5 Sodium Ascorbate, 7 MgCl_2_, 2 Ethylpyruvate, 0.5 CaCl_2_) and mounted in 4% LMP agarose. The lumbar spinal cords were sliced in a sagittal plane (200 μm, VT1200S, Leica), and the spinal cord slices were recovered at room temperature for 30 minutes in oxygenated (95% O_2_ and 5% CO_2_) ACSF solution (in mM: 2.5 CaCl_2_ 2H_2_O, 1.0 NaH_2_PO_4_ H2O, 119 NaCl, 2.5 KCl, 1.3 MgSO_4_ 7H_2_O, 26 NaHCO_3_, 25 Glucose, 1.3 Sodium L-ascorbate). Spinal cord slices were then transferred to a recording and cells were visualized by infrared differential interference contrast microscopy for patching (SliceScope Pro 6000, Scientifica). Whole-cell voltage-clamp recordings of random spinal cord neurons right below substantia gelatinosa were obtained at room temperature. The pipette resistance ranged from 3 to 6 MΩ, and the electrodes were filled with an intracellular solution (in mM: 130 Cs-gluconate, 5 TEA, 0.1 CaCl_2_, 1 EGTA, 10 HEPES, 1 QX-314, 0.2 D-600). Signals were acquired using a Multiclamp 700 B (Molecular Devices). Spontaneous miniature excitatory postsynaptic currents (mEPSCs) were monitored in the presence of tetrodotoxin (0.5 μM, Tocris Cat#1069) and 4-Aminopyridine (1 mM)., at -60 mV holding potential. Series resistance was left uncompensated. Synaptic currents were sampled at 10 kHz using a Digidata 1440A (Molecular Devices) and analyzed offline using Clampfit 9 (Molecular Devices) software. Events were analyzed using template matching search and each event was visually inspected for inclusion or rejection by an experimenter blind to genotype.

#### Behavioral Analysis

Both male and female mice of the C57BL6J background with a combination of *Advillin^Cre^, Pcdhg^fcon3/fcon3^,* and *Pcdhg^fcon3/+^* transgenes were used for the behavioral tests. Pregnant females were separated from breeding males before giving birth, and the progeny were allowed to reach weaning age when ear notching and genotyping was performed. Both male and female mutant animals were group-housed with their control littermates. For these assays, *Advillin^Cre^; Pcdhg^fcon3/+^*, *Advillin^Cre^, Pcdhg^fcon3/+^*and *Pcdhg^fcon3/fcon3^* were used as littermate control animals and were used to account for variabilities in the backgrounds of the animals tested. No differences were observed among these control genotypes. Testing was initiated at 6 to 8 weeks of age, and the animals completed Open Field, EPM, PPI, and Von Frey testing within 2 weeks of the start of testing and in that order. Balance beam and rough floor aversion were completed after this and held consistent within behavior cohorts. Prior to testing dates, animals were habituated to each new testing environment or room for approximately one hour to help eliminate anxiety during testing. All testing and analysis were performed by experimenters blind to genotypes.

#### Open Field Testing

Animals were habituated to the testing room in their cages with dim lights for 1 hour prior to testing. The lighting in the room was dimmed to limit anxiety as much as possible and to ensure the assays were as accurate to the natural behavior of the animals as possible. For this test, two 40 cm^3^ plexiglass arenas (40 cm x 40 cm x 40 cm) were used to test 2 animals concurrently and labeled for easy identification of the animals in each chamber during video analysis. The chambers utilized were opaque black and were cleaned thoroughly with 70% ethanol and water with paper towels before and after each trial. For the test, an animal was placed in the chamber and allowed to explore the chamber for 10 minutes. Following testing, videos of the animals’ position within the chamber were analyzed with custom MATLAB scripts. Total distance traveled and time spent away from walls (time in center) were analyzed for each video.

#### Elevated Plus Maze (EPM) Test

The EPM test was used to measure anxiety-like behavior. The custom-made maze utilized was constructed in the shape of a plus sign. The arms were each 30 cm long x 5 cm wide and two of the opposing arms have 15-cm high walls. The maze itself was raised on stilts to measure 40 cm above ground. Testing occurred in a dimly lit and quiet room, and the maze was positioned under a camera for video analysis following testing. After the animals were habituated to the testing room for at least 1 hour, they were placed on the center of the EPM construct and allowed to freely explore the open and closed arms of the maze for 10 minutes. MATLAB scripts were used to quantify the amount of the time each animal spent in the open and closed arms, as well as the center of the maze.

#### Prepulse Inhibition (PPI)

A San Diego Instruments startle reflex system (SR-LAB Startle Response System) was used to quantify the startle response of animals subjected to tactile and acoustic stimuli and measure the ability of a tactile or acoustic stimulus to inhibit startle response to a loud acoustic stimulus. Tactile PPI assay utilized a 50 ms 0.9 PSI air puff pre-stimulus delivered to the back of the animal, followed by a 20 ms 125 dB startle tone stimulus, to determine the responsiveness to tactile sensitivity in the back hairy skin.

For the tactile PPI test, ventilated cylindrical holders with removable and adjustable doors were used to contain the animals during testing. These holders also had a small opening on the top of the chamber to apply air puff to the back hairy skin. Both before and after testing, these cylindrical holders and the testing chambers were thoroughly cleaned with 70% ethanol and paper towels. The animals were placed in the holder, and the length of the holder were adjusted to match the length of the animal and eliminate extraneous movement during testing. The cylindrical holders were placed in the sanitized soundproof testing chambers. The testing consisted of an acclimation phase, and block I, block II, block III and block IV trials. The acclimation phase lasted 5 minutes, allowing the animal to become adjusted to the chambers and background noise of 75 dB. Block I of testing consisted of 5 startle stimuli alone and utilized 120 or 125 dB white noise to determine the initial startle reflex of the animal for measuring the initial startle reflex. Block II consisted of 5 prepulse stimuli alone (0.9 PSI air puff) to measure response to the prepulse stimulus. Block III utilized pseudorandomized prepulse/pulse, pulse alone and no stimulation trials. Finally, Block IV tested the animals’ habituation to the startle stimuli over the approximately total PPI session by delivering startle stimuli alone. Each trial lasted an average of 30 seconds. Postnatal day 5 mice were tested for reactivity to air puff stimuli applied to the back. This behavioral assay will be described in detail elsewhere.

For the acoustic PPI test, the acclimation tone was 65 dB during the acclimation phase. For Block I, the broadband white noise was 120 dB for determination of the initial startle reflex. For Block II, the prepulse stimuli consisted of 5 80 dB broadband white noise. Block III also utilized prepulse and pulse, pulse alone, and no stimulation trials which were used in a pseudo-randomized fashion. Block IV tested the animal’s habituation to the acoustic startle stimuli with the delivery of 5 startle stimuli.

#### Von Frey

Materials for this assay included transparent 4.5 in x 2 in x 2 in chambers with numerous 1cm wide ventilation holes, a wire grid stabilized between an acrylic frame and elevated to 14.5 in by metal columns, and an assortment of filaments with various forces. Prior to testing, animals were habituated to the testing setup by being placed on the wire grid inside the plastic chambers for 2 consecutive days for an hour. On the third day, the animals were habituated for 45 min before testing. Following habituation, the Von Frey filaments were applied to hairless plantar surface of the right hind paw until the filament buckled. Filaments were used beginning with the lightest force to the greatest, completing all trials with one filament before beginning the next. Each filament was used for 10 trials, with number of responses recorded. A withdrawing, sharp movement of the paw, or shaking of the paw behavior was defined as a response.

#### Balance beam assay

One hour prior to testing animals were habituated to testing room and investigator handling by undergoing tail inking. To ink the tail, each animal was gently lifted and placed on the wire food hopper facing away from investigator. Firmly holding the tail midway from the tail base, a blue soy ink marker was rolled across the tail forming parallel lines to indicate identifying ear notch number. Once inked animals were gently transferred back into the home cage.

The balance beam rod was constructed of 1 meter matte white acrylic with a flat surface measuring 12 mm or 6 mm in width. The beam rods were mounted on two poles 50 cm above the bench top. A black matte acrylic box was placed at each end of the beam to provide a start and end enclosure for the animal. To encourage the animal to traverse the beam home cage nesting material was placed in the end box to draw the animal to the end box. A hammock was stretched below the beam 10cm above the bench top to serve as a soft landing surface to cushion falls and contain the animal should a fall occur. Digital USB 2.0 CMOS video cameras were mounted above and to side of the balance beam to record crossings.

Balance Beam Training: Day 1 and Day 2. Animals are transported to the testing room at least 30 minutes prior to testing to acclimate to room lighting, sound, odor, and temperature. To train the animals to cross the beams unassisted, each animal was placed on the 12 mm rod and encouraged to cross at least 3 times before returning to the home cage. Once the cohort has rested for 30 minutes, they are then placed on the 6 mm rod and encouraged to complete 3 crossings. If animals refuse to cross, the investigator may gently touch the hindquarters to encourage forward movement. The beam and chambers were cleaned with dilute soapy water and 70% ethanol between animals.

Balance Beam Test: Day 3. First, the 12 mm beam was mounted onto the support poles with the start and end chambers connected to the beam ends. Each beam habituated animal was place into the start chamber with home cage bedding added to the end chamber. At least 2-4 successful crossings were video recorded for each animal. Each animal was allowed to rest for 10 seconds before each crossing. The animals were returned to their home cage once 2-4 successful trials were completed for each beam width. Time to cross, number of slips, number of falls and number of stalls were analyzed.

#### Texture aversion assay

One hour prior to testing animals were habituated to testing room and investigator handling by undergoing tail inking. The two test chambers constructed of black matte (inside) acrylic, 12 in x 11.8 in x 6 in, 0.25 in thick, centered side by side on a white matte acrylic floor under diffuse warm white light, 2700 K. A digital USB 2.0 CMOS video camera was mounted directly above the two test chambers. Testing for each texture was conducted over three days and consisted of two chamber habituation days and one test day. On day 1 and day 2 each animal was removed from the home cage placed in an empty test chamber resting on the white matte acrylic floor. Animals were allowed to freely explore and acclimate to the test environment for 10 minutes, then returned to their home cages. On day 3, the texture aversion assay consisted of two phases: a 10 minute explore phase and a 10 minute test phase. In the explore phase, the white matte acrylic floor was covered with a fresh sheet of construction paper to act as the control texture. Animals were placed in the test chamber and allowed to explore for 10 minutes, then returned to their home cage. The control paper was then discarded, and the test chambers were cleaned with 70% ethanol and unscented soapy water, then reset for the test phase. In the aversive texture test phase, two distinct sandpaper textures, extra fine grit SP400 (smooth) and medium coarse grit SP150 (rough) or SP400 (smooth) and averse coarse grit SP60 were taped together using 3M 1-inch scotch tape on the back of the paper with a ½ inch overlap. The test chambers were then placed onto the sandpaper sheets with the seams of the joined sandpaper centered in the middle of the test chamber, ensuring each texture covered half of the testing chamber. Animals were once again transferred from their home cage to the test chamber and allowed to explore the two sandpaper textures SP400 vs SP150 or SP400 vs SP60 for 10 minutes. At the completion of the 10 minute averse texture test animals were removed from the test chambers and returned to their home cage. Both exploration and test phases were video-recorded and custom Matlab tracking codes were used to track animal position in the chamber, time spent on each texture and preference for smooth vs rough textures were analyzed.

#### Artificial synapse formation assay

Primary spinal cord cultures were prepared from dorsal spinal cords of P0 mice and maintained in neurobasal medium with B27 supplement. P0 C57BL/6 mouse pups were anesthetized on ice and then decapitated using scissors. Spinal columns were dissected and washed with filter-sterilized HABG medium (Hibernate A with 2% B-27 and 0.5 mM GlutaMAX) to remove excess blood and tissue, as previously described (Eldeiry et al., 2017). 22 G needle and syringe filled with filter-sterilized HABG was inserted into the caudal end of the spinal column and flushed cranially, allowing the cord to exit into a Petri dish. Dorsal spinal cord is cut into small pieces about 0.5 mm in size. Spinal cord tissues were placed in HABG medium in a 30 °C incubator for 30 min and then digested with Papain (Sigma, Cat#10108014001) for 30 min at 30 °C (HA-Ca 10 mL, Papain, 25 mg, and 0.5 mM GlutaMAX). Papain solution was aspirated and spinal cord tissues were washed with 10 ml neurobasal media (Neurobasal A, 2% B-27, 0.5 mM GlutaMAX, and 1% Pen/Strep). Tissues were gently dissociated by pipetting in neurobasal media with a flame-polished Pasteur pipette. Cells were plated on laminin-covered cover glasses in tissue culture plates and placed in the 37 °C cell culture incubator.

Artificial synapse formation assays were performed as previously described (Biederer and Scheiffele, 2007). HEK 293 cells were maintained in DMEM (Thermo Fisher Scientific, Cat# 10564-011) with 10% HyClone™ Cosmic Calf™ Serum (VWR, Cat# SH3008703), 1% Sodium Pyruvate (Gibco, Cat# 11360-070), 1% MEM Non-Essential Amino Acids Solution (Gibco, Cat# 11140-050) and 1% Pennicillin-Streptomycin (Simga, Cat# P4333). HEK 293 cells were cultured in six-well plates and transfected at 50-70% confluency by using polyethylenimine according to standard protocol (4 ug of DNA mixed with 8 ug of polyethylenimine). One day after transfection, HEK 293 cells were resuspended and plated into dorsal spinal cord neurons at day 8 *in vitro*. Cells were fixed with 4% PFA for 10 min at room temperature and then were washed three times for 5 min with 1 ml of 0.3% PBST. The fixed cells were blocked with blocking buffer (5% normal goat serum in 0.3% PBST) for 2 h at RT. Then, primary antibody diluted in the blocking buffer was added to the wells at 4°C overnight. Cells were washed 3 × 10 min with 1 ml of 0.1% PBST, followed by secondary antibodies incubation for 1 hour at room temperature. The coverslips were washed for 3X 10 min before mounting on the Superfrost slides.

For fluorescent image analysis, cells were chosen randomly and imaged from four independent replicates. Fluorescent images were acquired with a Zeiss 700 confocal microscope using a Zeiss Achroplan 63/0.95 at 1,024 by 1,024-pixel resolutions. No detectable bleed-through occurred between different channels. The pixel intensities for the brightest samples were below saturation, with the exception that when contours of the HA or GFP signal from transfected HEK 293 cells need to be clearly defined. MAP2 channel was used to confirm dendrites contacting HEK 293 cells. vGlut2 channel was used to exclude extensive neuronal synapses formed near or within HEK293 cells. Images were using identical confocal settings. For analysis, images were thresholded by intensity, and PSD-95 puncta were quantified by counting the number of thresholded puncta. The puncta number and total integrated intensity of PSD-95 punta within the HEK 293 cells were calculated using ImageJ software. Both image acquisition and quantification were performed by experimenters who were blind to the experimental conditions.

#### *In vivo* spinal cord multielectrode array electrophysiology

Adult (>6 weeks) animals were administered dexamethasone (2 mg/kg) 1-2 hours prior to recording to prevent tissue swelling and were then anesthetized with urethane (1.5 g/kg, Sigma). Isoflurane (1%) was administered during surgery and removed prior to multielectrode array recordings. Surgical plane of anesthesia and the temperature of the animal were monitored and maintained throughout the recording session. Briefly, the hair surrounding the dorsal hump was shaved and the spinal column of T13 to L6 was exposed. The vertebrae above the recording site were stabilized using clamps. The vertebrae between L4 and L5 were retracted to expose the dorsal surface of the spinal cord and the dura was removed. The surface of the cord was bathed in mineral oil/submerged in saline. A 32-channel silicon probe (Neuronexus A1x32-Poly3-5mm-25s-177-

A32 or Cambridge Neurotech ASSY-37 H4 optrode) was inserted into the dorsal horn and advanced up to ∼700 μm below the dorsal surface. The probe was then kept in place for 20 minutes to ensure a stable recording. Signals were amplified, filtered (0.1 - 7.5 kHz bandpass), and digitized (20 kHz) using a headstage amplifier and recording controller (Intan Technologies RHD2132 and Recording Controller). Data acquisition was controlled with Intan Technologies Recording Controller (version 2.07). A 150-200 µm diameter Teflon-tipped indenter controlled by a dual-mode force controller (Aurora Scientific 300C-I) was used to stimulate the glabrous hindpaw. For mapping receptive field (RF) areas, the position of the indenter was controlled with two linear translation piezo stages and a stage controller (Physik Instrumente U-521.24 and C-867.2U2). The position, force, and displacement of the indenter were commanded with custom-written Matlab scripts.

The detailed data analyses will be described elsewhere. Briefly, JRCLUST (version 3.2.2) was used to automatically sort action potentials into clusters, manually refine clusters, and classify clusters as single or multi units. Action potentials were detected with a threshold of 4.5 times the standard deviation of the noise. Action potentials with similar times across sites were merged and then sorted into clusters. Outlier spikes were removed from each cluster. To isolate putative single units, manual cluster curation was performed with JRCLUST split and merge tools. Clusters were classified as single units if (1) the waveforms were large with respect to baseline; (2) there was a clear refractory period in the cross-correlogram (interspike intervals > 1 ms); (3) waveforms were clearly distinct from nearby clusters. Spike event times for clusters classified as single units were exported and processed in Python.

### QUANTIFICATION AND STATISTICAL ANALYSIS

All data are presented as the mean ± the standard error of the mean (SEM), unless otherwise noted in the figure legend. Student’s t test is used to compare the morphological/physiological comparisons between littermate control and mutants. One-way ANOVA tests are used to compare three or more groups in one condition and are expressed as an F-statistic and P value within brackets, and post hoc comparisons were performed using the post hoc test noted in the figure legend. Two-way ANOVA tests are used to compare at least two groups with multiple conditions. The number of cells or animals per group used in each experiment is described in the figure legends. Unless otherwise noted, *, p < 0.05, **, p < 0.01, and ***, p < 0.001. All statistics were performed using GraphPad Prism.

---

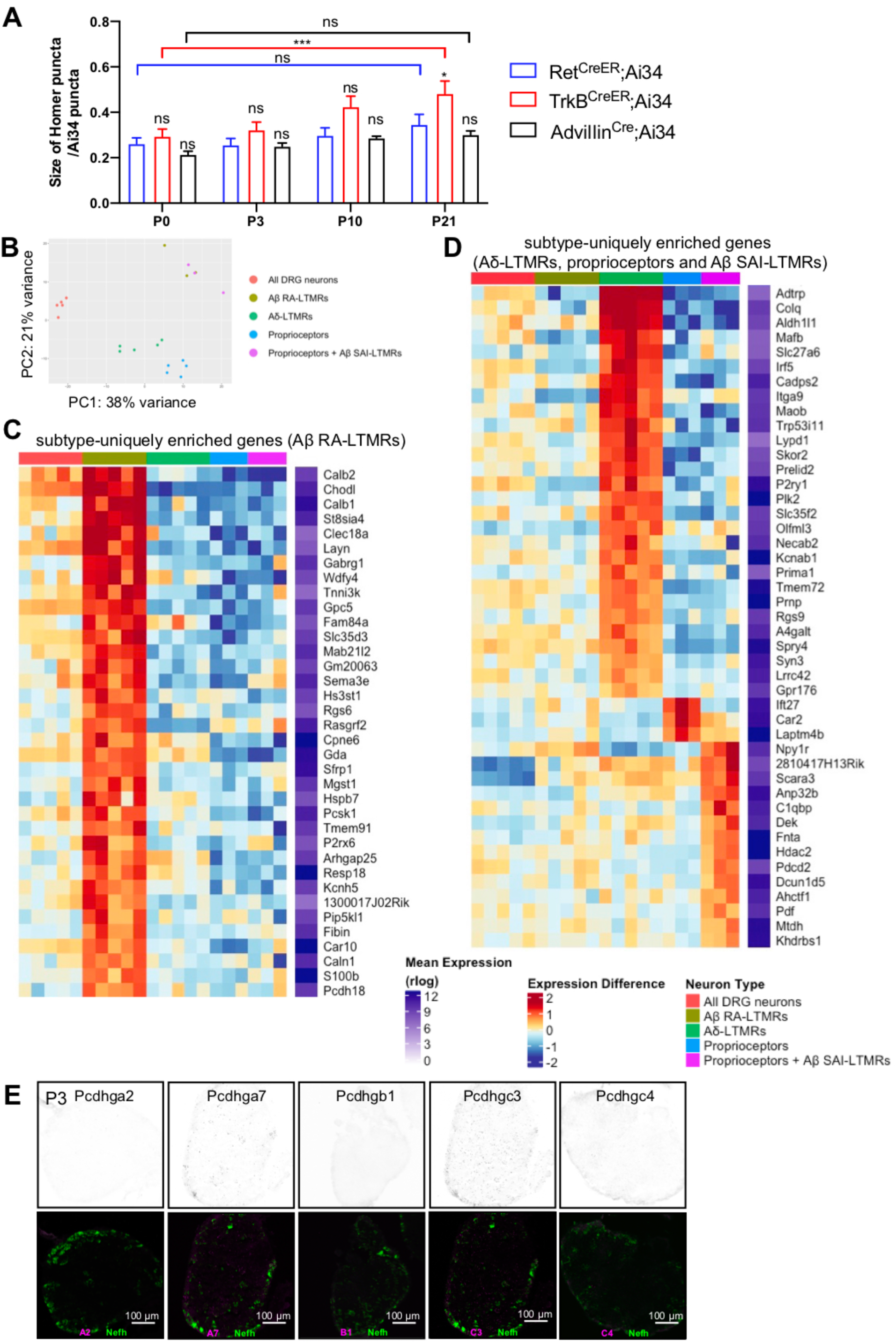

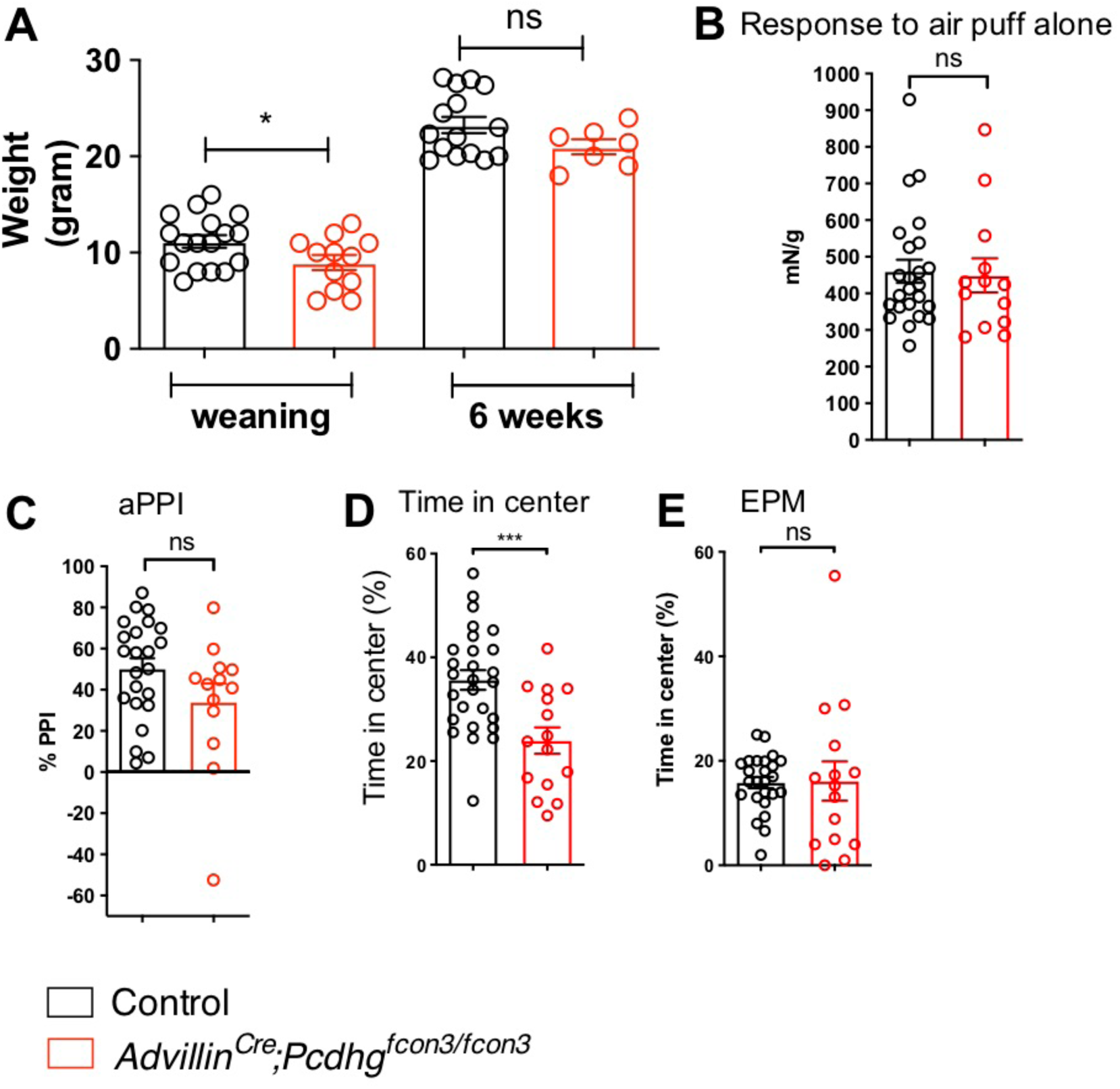

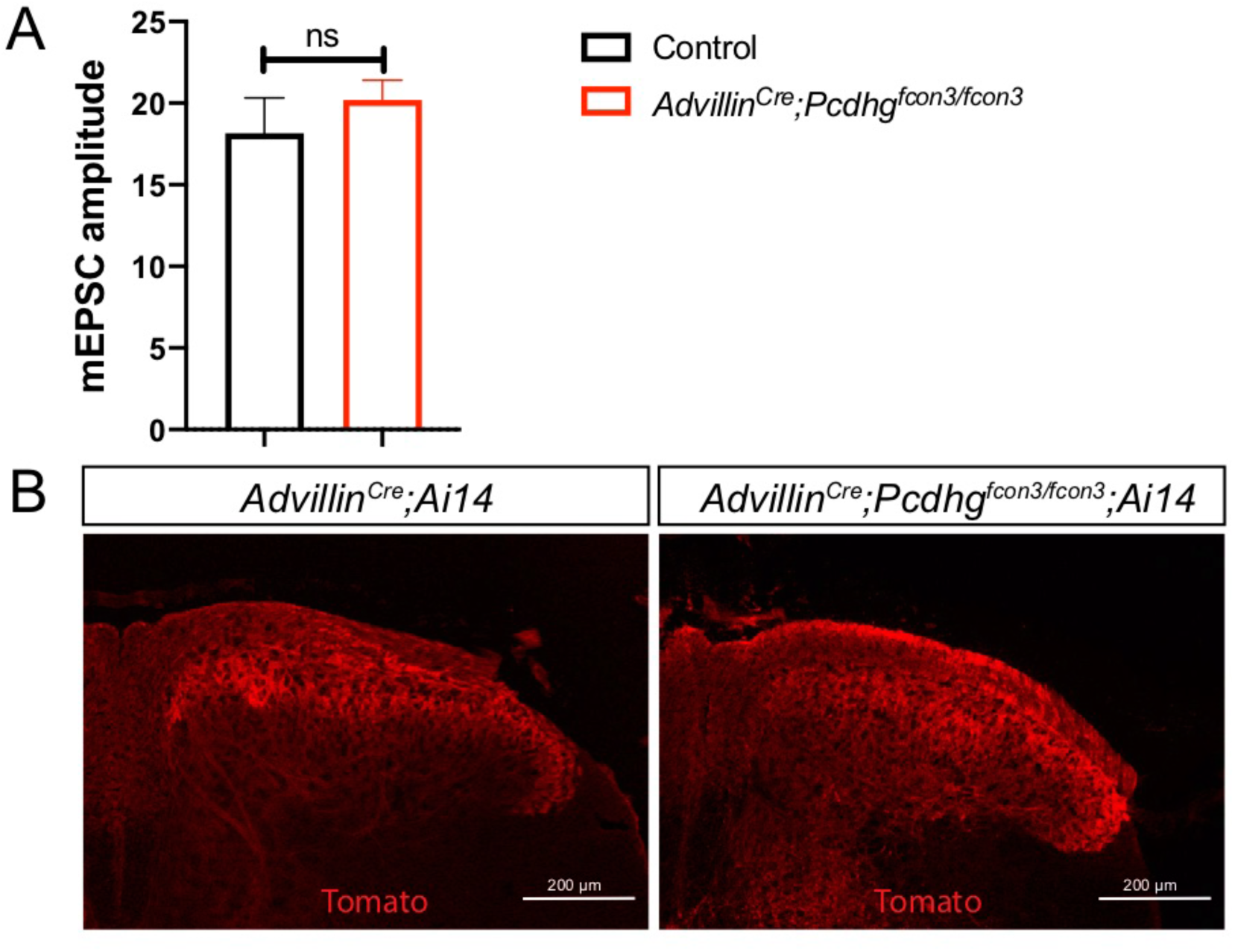

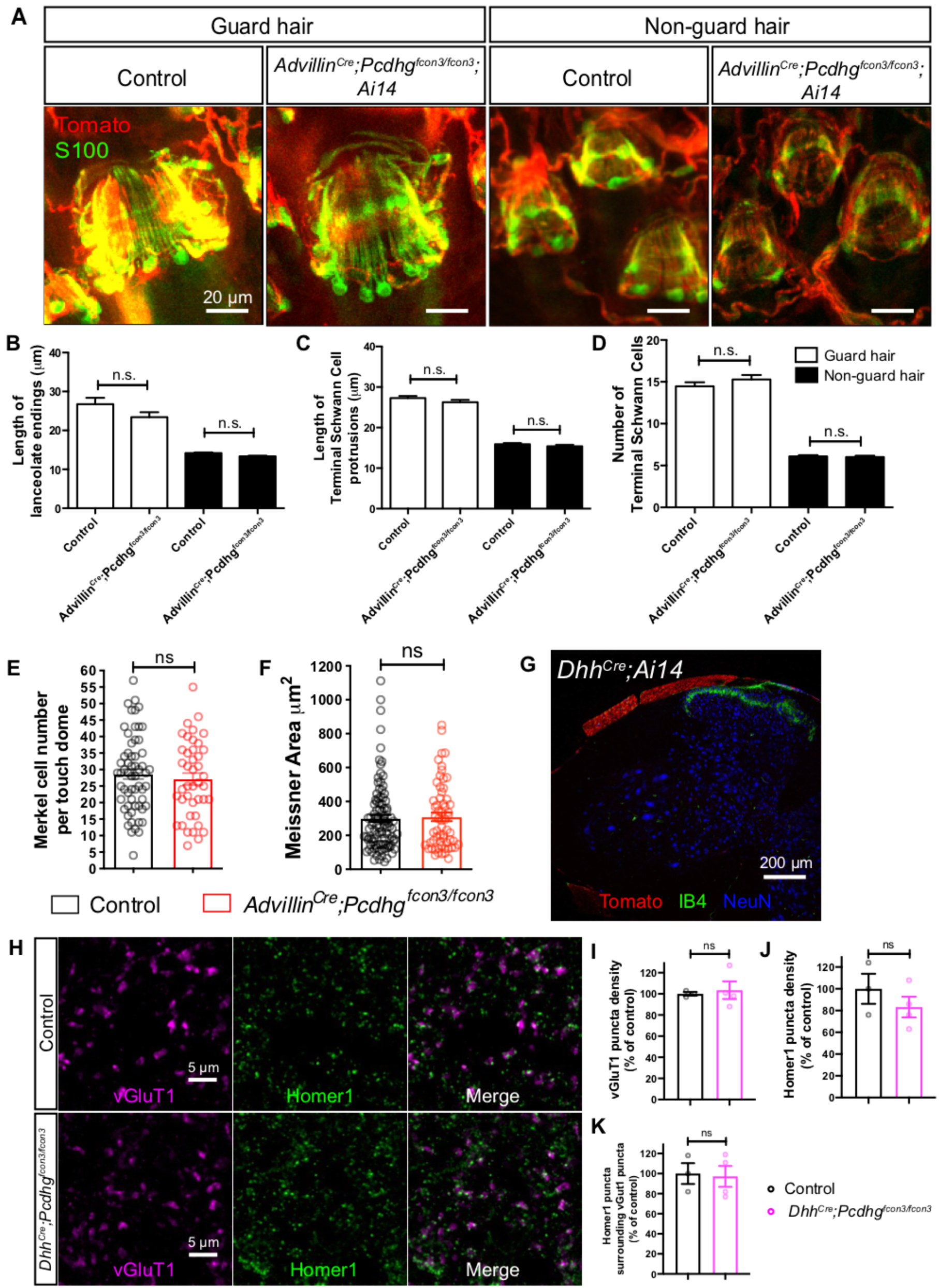

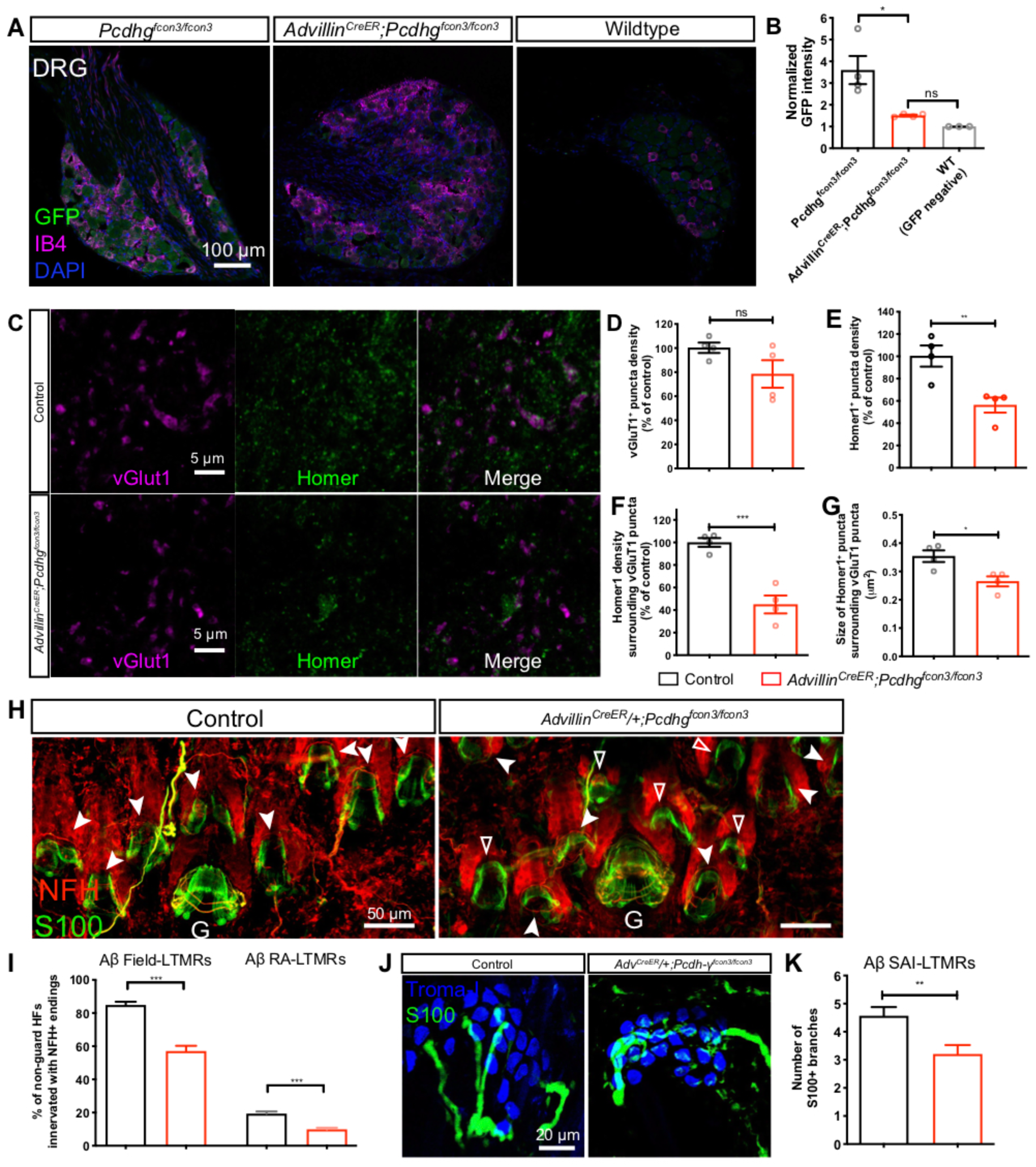

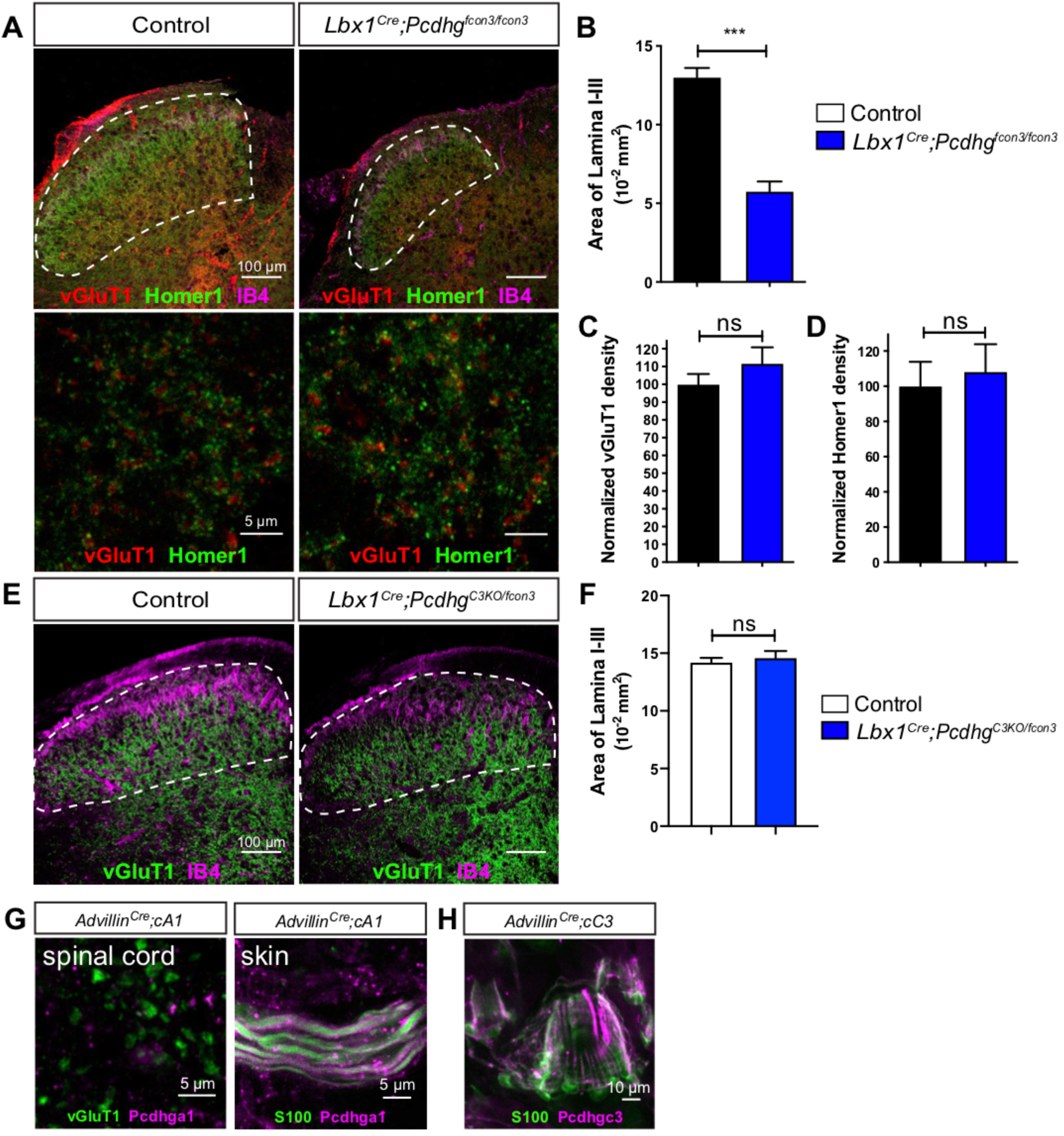

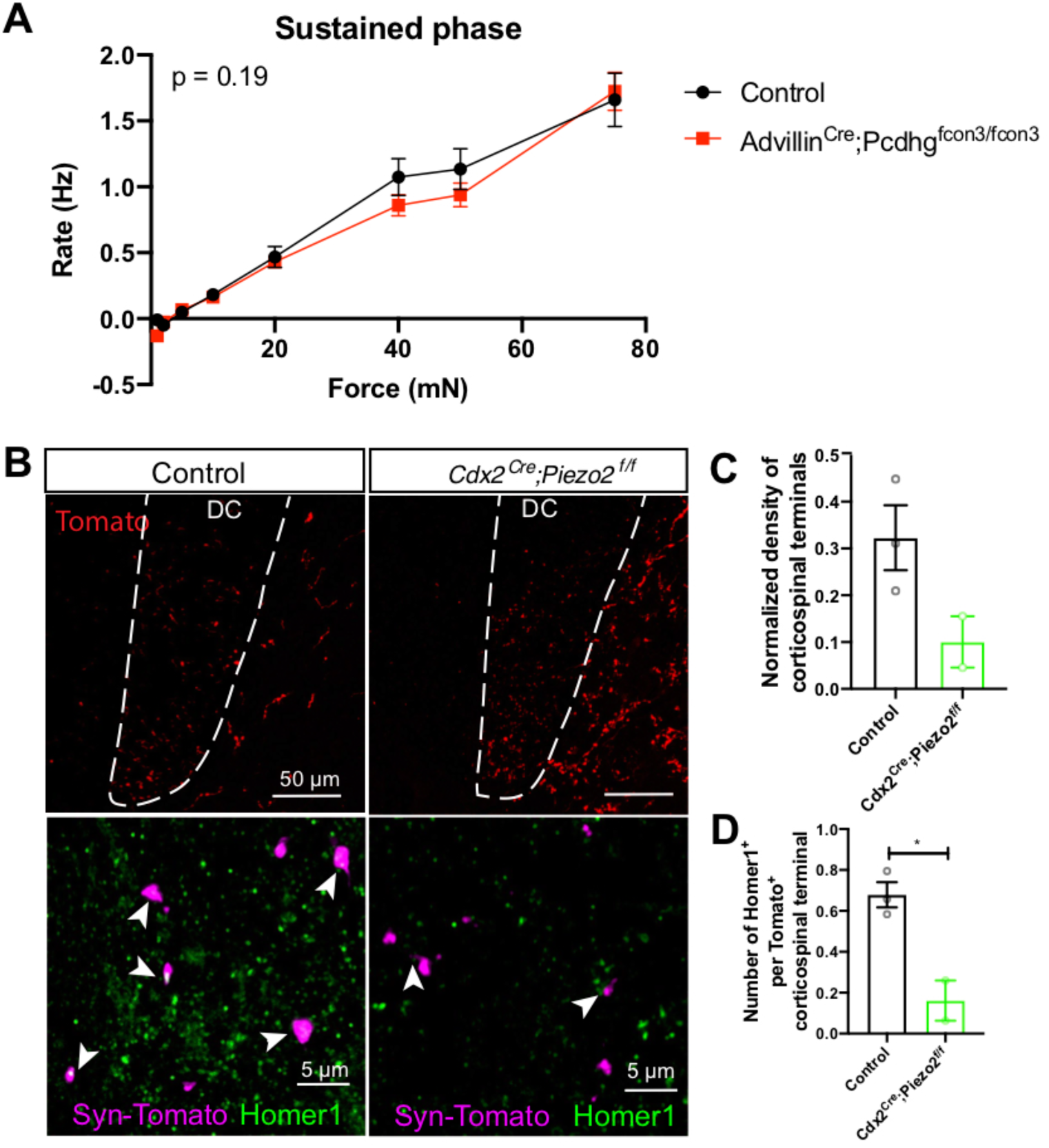

## References

Abraira, V.E., and Ginty, D.D. (2013). The Sensory Neurons of Touch. Neuron 79, 618–639.

Abraira, V.E., Kuehn, E.D., Chirila, A.M., Springel, M.W., Toliver, A.A., Zimmerman, A.L., Orefice, L.L., Boyle, K.A., Bai, L., Song, B.J., et al. (2017). The Cellular and Synaptic Architecture of the Mechanosensory Dorsal Horn. Cell 168, 295–310.e19.

Andrews, K., and Fitzgerald, M. (1994). The cutaneous withdrawal reflex in human neonates: sensitization, receptive fields, and the effects of contralateral stimulation. Pain 56, 95–101.

Bai, L., Lehnert, B.P., Liu, J., Neubarth, N.L., Dickendesher, T.L., Nwe, P.H., Cassidy, C., Woodbury, C.J., and Ginty, D.D. (2015). Genetic Identification of an Expansive Mechanoreceptor Sensitive to Skin Stroking. Cell 163, 1783–1795.

Beggs, S., Torsney, C., Drew, L.J., and Fitzgerald, M. (2002). The postnatal reorganization of primary afferent input and dorsal horn cell receptive fields in the rat spinal cord is an activity-dependent process. Eur J Neurosci 16, 1249–1258.

Biederer, T., and Scheiffele, P. (2007). Mixed-culture assays for analyzing neuronal synapse formation. Nature Protocols 2, 670–676.

Brasch, J., Goodman, K.M., Noble, A.J., Rapp, M., Mannepalli, S., Bahna, F., Dandey, V.P., Bepler, T., Berger, B., Maniatis, T., et al. (2019). Visualization of clustered protocadherin neuronal self-recognition complexes. Nature 569, 280–283.

Burgess, P.R., Petit, D., and Warren, R.M. (1968). Receptor types in cat hairy skin supplied by myelinated fibers. J Neurophysiol 31, 833–848.

Canzio, D., and Maniatis, T. (2019). The generation of a protocadherin cell-surface recognition code for neural circuit assembly. Current Opinion in Neurobiology 59, 213–220.

Chen, W.V., Alvarez, F.J., Lefebvre, J.L., Friedman, B., Nwakeze, C., Geiman, E., Smith, C., Thu, C.A., Tapia, J.C., Tasic, B., et al. (2012). Functional Significance of Isoform Diversification in the Protocadherin Gamma Gene Cluster. Neuron 75, 402–409.

Chen, W.V., Nwakeze, C.L., Denny, C.A., O’Keeffe, S., Rieger, M.A., Mountoufaris, G., Kirner, A., Dougherty, J.D., Hen, R., Wu, Q., et al. (2017). Pcdhαc2 is required for axonal tiling and assembly of serotonergic circuitries in mice. Science 356, 406–411.

Choi, S., Hachisuka, J., Brett, M.A., Magee, A.R., Omori, Y., Iqbal, N.-U.-A., Zhang, D., DeLisle, M.M., Wolfson, R.L., Bai, L., et al. (2020). Parallel ascending spinal pathways for affective touch and pain. Nature 587, 258–263.

Coutaud, B., and Pilon, N. (2013). Characterization of a novel transgenic mouse line expressing Cre recombinase under the control of the Cdx2 neural specific enhancer. Genesis 51, 777–784.

Dani, A., Huang, B., Bergan, J., Dulac, C., and Zhuang, X. (2010). Superresolution imaging of chemical synapses in the brain. Neuron 68, 843–856.

Dobin, A., Davis, C.A., Schlesinger, F., Drenkow, J., Zaleski, C., Jha, S., Batut, P., Chaisson, M., and Gingeras, T.R. (2013). STAR: ultrafast universal RNA-seq aligner. Bioinformatics 29, 15–21.

Ebert, D.H., and Greenberg, M.E. (2013). Activity-dependent neuronal signalling and autism spectrum disorder. Nature 493, 327–337.

Eldeiry, M., Yamanaka, K., Reece, T.B., and Aftab, M. (2017). Spinal Cord Neurons Isolation and Culture from Neonatal Mice. Journal of Visualized Experiments 125, 1–9.

Fernandez-Monreal, M., Kang, S., and Phillips, G.R. (2009). Gamma-protocadherin homophilic interaction and intracellular trafficking is controlled by the cytoplasmic domain in neurons. Mol Cell Neurosci 40, 344–353.

Fitzgerald, M. (1985). The post-natal development of cutaneous afferent fibre input and receptive field organization in the rat dorsal horn. J. Physiol. (Lond.) 364, 1–18.

Fitzgerald, M. (1987a). Spontaneous and evoked activity of fetal primary afferents in vivo. Nature 326, 603–605.

Fitzgerald, M. (1987b). Cutaneous primary afferent properties in the hind limb of the neonatal rat. J. Physiol. (Lond.) 383, 79–92.

Flaherty, E., and Maniatis, T. (2020). The role of clustered protocadherins in neurodevelopment and neuropsychiatric diseases. Current Opinion in Genetics & Development 65, 144–150.

Frank, M., Ebert, M., Shan, W., Phillips, G.R., Arndt, K., Colman, D.R., and Kemler, R. (2005). Differential expression of individual gamma-protocadherins during mouse brain development. Molecular and Cellular Neuroscience 29, 603–616.

Frey, M.V. (1896). Untersuchunger über die Sinnesfunctionen der menschlichen Haut. Bandes der Abhandlungen der mathematisch-physischen Classe der Königl. Sächsischen Gesellschaft Der Wissenschaften 23, 175–266.

Garrett, A.M., and Weiner, J.A. (2009). Control of CNS synapse development by γ-protocadherin-mediated astrocyte-neuron contact. J. Neurosci. 29, 11723–11731.

Garrett, A.M., Bosch, P.J., Steffen, D.M., Fuller, L.C., Marcucci, C.G., Koch, A.A., Bais, P., Weiner, J.A., and Burgess, R.W. (2019). CRISPR/Cas9 interrogation of the mouse Pcdhg gene cluster reveals a crucial isoform-specific role for Pcdhgc4. PLoS Genet 15, e1008554.

Garrett, A.M., Schreiner, D., Lobas, M.A., and Weiner, J.A. (2012). γ-protocadherins control cortical dendrite arborization by regulating the activity of a FAK/PKC/MARCKS signaling pathway. Neuron 74, 269–276.

Gibson, D.A., Tymanskyj, S., Yuan, R.C., Leung, H.C., Lefebvre, J.L., Sanes, J.R., Chédotal, A., and Ma, L. (2014). Dendrite self-avoidance requires cell-autonomous slit/robo signaling in cerebellar purkinje cells. Neuron 81, 1040–1056.

Goodman, K.M., Rubinstein, R., Dan, H., Bahna, F., Mannepalli, S., Ahlsen, G., Aye Thu, C., Sampogna, R.V., Maniatis, T., Honig, B., et al. (2017). Protocadherin cis-dimer architecture and recognition unit diversity. Proc. Natl. Acad. Sci. U.S.a. 114, E9829–E9837.

Goodman, K.M., Katsamba, P.S., Rubinstein, R., Ahlsen, G., Bahna, F., Mannepalli, S., Dan, H., Sampogna, R.V., Shapiro, L., and Honig, B. (2022). How clustered protocadherin binding specificity is tuned for neuronal self-/nonself-recognition. eLife 11.

Goodman, K.M., Rubinstein, R., Thu, C.A., Bahna, F., Mannepalli, S., Ahlsen, G., Rittenhouse, C., Maniatis, T., Honig, B., and Shapiro, L. (2016a). Structural Basis of Diverse Homophilic Recognition by Clustered α- and β-Protocadherins. Neuron 90, 709–723.

Goodman, K.M., Rubinstein, R., Thu, C.A., Mannepalli, S., Bahna, F., Ahlsen, G., Rittenhouse, C., Maniatis, T., Honig, B., and Shapiro, L. (2016b). γ-Protocadherin structural diversity and functional implications. eLife 5.

Graf, E.R., Zhang, X., Jin, S.-X., Linhoff, M.W., and Craig, A.M. (2004). Neurexins Induce Differentiation of GABA and Glutamate Postsynaptic Specializations via Neuroligins. Cell 119, 1013–1026.

Gutierrez-Mecinas, M., Kuehn, E.D., Abraira, V.E., Polgár, E., Watanabe, M., and Todd, A.J. (2016). Immunostaining for Homer reveals the majority of excitatory synapses in laminae I-III of the mouse spinal dorsal horn. Neuroscience 329, 171–181.

Han, M.-H., Lin, C., Meng, S., and Wang, X. (2010). Proteomics Analysis Reveals Overlapping Functions of Clustered Protocadherins. Mol Cell Proteomics 9, 71–83.

Hanaway, J., and Smith, J.M. (1979). Synaptic fine structure and the termination of corticospinal fibers in the lateral basal region of the cat spinal cord. J. Comp. Neurol. 183, 471–486.

Handler, A., and Ginty, D.D. (2021). The mechanosensory neurons of touch and their mechanisms of activation. Nat Rev Neurosci 22, 521–537.

Hasegawa, H., Abbott, S., Han, B.-X., Qi, Y., and Wang, F. (2007). Analyzing somatosensory axon projections with the sensory neuron-specific Advillin gene. J. Neurosci. 27, 14404–14414.

Hasegawa, S., Kumagai, M., Hagihara, M., Nishimaru, H., Hirano, K., Kaneko, R., Okayama, A., Hirayama, T., Sanbo, M., Hirabayashi, M., et al. (2016). Distinct and Cooperative Functions for the Protocadherin-α, -β and -γ Clusters in Neuronal Survival and Axon Targeting. Front. Mol. Neurosci. 9, 155.

Hertenstein, M.J., Verkamp, J.M., Kerestes, A.M., and Holmes, R.M. (2006). The communicative functions of touch in humans, nonhuman primates, and rats: a review and synthesis of the empirical research. Genet Soc Gen Psychol Monogr 132, 5–94.

Hippenmeyer, S., Vrieseling, E., Sigrist, M., Portmann, T., Laengle, C., Ladle, D.R., and Arber, S. (2005). A developmental switch in the response of DRG neurons to ETS transcription factor signaling. PLoS Biol 3, e159.

Horch, K.W., Tuckett, R.P., and Burgess, P.R. (1977). A key to the classification of cutaneous mechanoreceptors. J Invest Dermatol 69, 75–82.

Hsu, J.-Y.C., Stein, S.A., and Xu, X.-M. (2006). Development of the corticospinal tract in the mouse spinal cord: a quantitative ultrastructural analysis. Brain Res. 1084, 16–27.

Jaegle, M., Ghazvini, M., Mandemakers, W., Piirsoo, M., Driegen, S., Levavasseur, F., Raghoenath, S., Grosveld, F., and Meijer, D. (2003). The POU proteins Brn-2 and Oct-6 share important functions in Schwann cell development. Genes Dev. 17, 1380–1391.

Jenkins, B.A., and Lumpkin, E.A. (2017). Developing a sense of touch. Development 144, 4078– 4090.

Jessen, K.R., Mirsky, R., and Lloyd, A.C. (2015). Schwann Cells: Development and Role in Nerve Repair. Cold Spring Harbor Perspectives in Biology 7, a020487.

Johnson, K.O. (1983). Neural mechanisms of tactual form and texture discrimination. Fed Proc 42, 2542–2547.

Keeler, A.B., Molumby, M.J., and Weiner, J.A. (2015). Protocadherins branch out: Multiple roles in dendrite development. Cell Adh Migr 9, 214–226.

Kirkby, L.A., Sack, G.S., Firl, A., and Feller, M.B. (2013). A role for correlated spontaneous activity in the assembly of neural circuits. Neuron 80, 1129–1144.

Kostadinov, D., and Sanes, J.R. (2015). Protocadherin-dependent dendritic self-avoidance regulates neural connectivity and circuit function. eLife 4: e08964.

Kuehn, E.D., Meltzer, S., Abraira, V.E., Ho, C.-Y., and Ginty, D.D. (2019). Tiling and somatotopic alignment of mammalian low-threshold mechanoreceptors. Proc. Natl. Acad. Sci. U.S.a. 116, 9168–9177.

LaMassa, N., Sverdlov, H., Mambetalieva, A., Shapiro, S., Bucaro, M., Fernandez-Monreal, M., and Phillips, G.R. (2021). Gamma-protocadherin localization at the synapse is associated with parameters of synaptic maturation. J. Comp. Neurol. 529, 2407–2417.

Lau, J., Minett, M.S., Zhao, J., Dennehy, U., Wang, F., Wood, J.N., and Bogdanov, Y.D. (2011). Temporal control of gene deletion in sensory ganglia using a tamoxifen-inducible Advillin-Cre-ERT2 recombinase mouse. Mol Pain 7, 100.

Lefebvre, J.L., Kostadinov, D., Chen, W.V., Maniatis, T., and Sanes, J.R. (2012). Protocadherins mediate dendritic self-avoidance in the mammalian nervous system. Nature 488, 517–521.

Lefebvre, J.L., Sanes, J.R., and Kay, J.N. (2015). Development of Dendritic Form and Function. Annu. Rev. Cell Dev. Biol. 31, 741–777.

Lefebvre, J.L., Zhang, Y., Meister, M., Wang, X., and Sanes, J.R. (2008). gamma-Protocadherins regulate neuronal survival but are dispensable for circuit formation in retina. Development 135, 4141–4151.

Lehnert, B.P., Santiago, C., Huey, E.L., Emanuel, A.J., Renauld, S., Africawala, N., Alkislar, I., Zheng, Y., Bai, L., Koutsioumpa, C., et al. (2021). Mechanoreceptor synapses in the brainstem shape the central representation of touch. Cell 184, 5608–5621.e5618.

Leighton, A.H., and Lohmann, C. (2016). The Wiring of Developing Sensory Circuits-From Patterned Spontaneous Activity to Synaptic Plasticity Mechanisms. Front Neural Circuits 10, 71.

Li, L., Rutlin, M., Abraira, V.E., Cassidy, C., Kus, L., Gong, S., Jankowski, M.P., Luo, W., Heintz, N., Koerber, H.R., et al. (2011). The functional organization of cutaneous low-threshold mechanosensory neurons. Cell 147, 1615–1627.

Li, Y., Serwanski, D.R., Miralles, C.P., Fiondella, C.G., Loturco, J.J., Rubio, M.E., and De Blas, A.L. (2010). Synaptic and nonsynaptic localization of protocadherin-gammaC5 in the rat brain. J. Comp. Neurol. 518, 3439–3463.

Loh, K.H., Stawski, P.S., Draycott, A.S., Udeshi, N.D., Lehrman, E.K., Wilton, D.K., Svinkina, T., Deerinck, T.J., Ellisman, M.H., Stevens, B., et al. (2016). Proteomic Analysis of Unbounded Cellular Compartments: Synaptic Clefts. Cell 166, 1295–1307.e21.

Love, M.I., Huber, W., and Anders, S. (2014). Moderated estimation of fold change and dispersion for RNA-seq data with DESeq2. Genome Biol. 15, 550–21.

Luo, W., Enomoto, H., Rice, F.L., Milbrandt, J., and Ginty, D.D. (2009). Molecular identification of rapidly adapting mechanoreceptors and their developmental dependence on ret signaling. Neuron 64, 841–856.

Mah, K.M., Houston, D.W., and Weiner, J.A. (2016). The γ-Protocadherin-C3 isoform inhibits canonical Wnt signalling by binding to and stabilizing Axin1 at the membrane. Nature Publishing Group 6, 31665–17.

Main, M., and Stadtman, J. (1981). Infant response to rejection of physical contact by the mother: aggression, avoidance, and conflict. J Am Acad Child Psychiatry 20, 292–307.

McGill, B.E., Bundle, S.F., Yaylaoglu, M.B., Carson, J.P., Thaller, C., and Zoghbi, H.Y. (2006). Enhanced anxiety and stress-induced corticosterone release are associated with increased Crh expression in a mouse model of Rett syndrome. Proc Natl Acad Sci USA 103, 18267–18272.

Meltzer, S., Santiago, C., Sharma, N., and Ginty, D.D. (2021). The cellular and molecular basis of somatosensory neuron development. Neuron 1–22.

Mirnics, K., and Koerber, H.R. (1995). Prenatal development of rat primary afferent fibers: II. Central projections. J. Comp. Neurol. 355, 601–614.

Missler, M., Südhof, T.C., and Biederer, T. (2012). Synaptic cell adhesion. Cold Spring Harbor Perspectives in Biology 4, a005694–a005694.

Molumby, M.J., Anderson, R.M., Newbold, D.J., Koblesky, N.K., Garrett, A.M., Schreiner, D., Radley, J.J., and Weiner, J.A. (2017). gamma-Protocadherins Interact with Neuroligin-1 and Negatively Regulate Dendritic Spine Morphogenesis. CellReports 18, 2702–2714.

Molumby, M.J., Keeler, A.B., and Weiner, J.A. (2016). Homophilic Protocadherin Cell-Cell Interactions Promote Dendrite Complexity. CellReports 15, 1037–1050.

Mountoufaris, G., Canzio, D., Nwakeze, C.L., Chen, W.V., and Maniatis, T. (2018). Writing, Reading, and Translating the Clustered Protocadherin Cell Surface Recognition Code for Neural Circuit Assembly. Annu. Rev. Cell Dev. Biol. 34, 471–493.

Nam, C.I., and Chen, L. (2005). Postsynaptic assembly induced by neurexin–neuroligin interaction and neurotransmitter. Proc Natl Acad Sci USA 102, 6137–6142.

Neubarth, N.L., Emanuel, A.J., Liu, Y., Springel, M.W., Handler, A., Zhang, Q., Lehnert, B.P., Guo, C., Orefice, L.L., Abdelaziz, A., et al. (2020). Meissner corpuscles and their spatially intermingled afferents underlie gentle touch perception. Science 368, eabb2751.

Nicoludis, J.M., Vogt, B.E., Green, A.G., Schärfe, C.P., Marks, D.S., and Gaudet, R. (2016). Antiparallel protocadherin homodimers use distinct affinity- and specificity-mediating regions in cadherin repeats 1-4. eLife 5.

Obata, S., Sago, H., Mori, N., Davidson, M., St John, T., and Suzuki, S.T. (1998). A common protocadherin tail: multiple protocadherins share the same sequence in their cytoplasmic domains and are expressed in different regions of brain. Cell Adhes Commun 6, 323–333.

Obata, S., Sago, H., Mori, N., Rochelle, J.M., Seldin, M.F., Davidson, M., St John, T., Taketani, S., and Suzuki, S.T. (1995). Protocadherin Pcdh2 shows properties similar to, but distinct from, those of classical cadherins. Journal of Cell Science 108 ( *Pt 12*), 3765–3773.

Olausson, H., Lamarre, Y., Backlund, H., Morin, C., Wallin, B.G., Starck, G., Ekholm, S., Strigo, I., Worsley, K., Vallbo, A.B., et al. (2002). Unmyelinated tactile afferents signal touch and project to insular cortex. Nat Neurosci 5, 900–904.

Olson, W., Dong, P., Fleming, M., and Luo, W. (2016). The specification and wiring of mammalian cutaneous lowLJthreshold mechanoreceptors. WIREs Dev Biol 5, 389–404.

Orefice, L.L. (2020). Peripheral Somatosensory Neuron Dysfunction: Emerging Roles in Autism Spectrum Disorders. Neuroscience 445, 120–129.

Orefice, L.L., Mosko, J.R., Morency, D.T., Wells, M.F., Tasnim, A., Mozeika, S.M., Ye, M., Chirila, A.M., Emanuel, A.J., Rankin, G., et al. (2019). Targeting Peripheral Somatosensory Neurons to Improve Tactile-Related Phenotypes in ASD Models. Cell 178, 867–886.e24.

Orefice, L.L., Zimmerman, A.L., Chirila, A.M., Sleboda, S.J., Head, J.P., and Ginty, D.D. (2016). Peripheral Mechanosensory Neuron Dysfunction Underlies Tactile and Behavioral Deficits in Mouse Models of ASDs. Cell 166, 299–313.

Ozaki, S., and Snider, W.D. (1997). Initial trajectories of sensory axons toward laminar targets in the developing mouse spinal cord. J. Comp. Neurol. 380, 215–229.

Paixão, S., Loschek, L., Gaitanos, L., Alcalà Morales, P., Goulding, M., and Klein, R. (2019). Identification of Spinal Neurons Contributing to the Dorsal Column Projection Mediating Fine Touch and Corrective Motor Movements. Neuron 104, 749–764.e6.

Panek, I., Bui, T., Wright, A.T.B., and Brownstone, R.M. (2014). Cutaneous afferent regulation of motor function. Acta Neurobiol Exp (Wars) 74, 158–171.

Peek, S.L., Mah, K.M., and Weiner, J.A. (2017). Regulation of neural circuit formation by protocadherins. Cell Mol Life Sci 74, 4133–4157.

Phillips, G.R., Tanaka, H., Frank, M., Elste, A., Fidler, L., Benson, D.L., and Colman, D.R. (2003). Gamma-protocadherins are targeted to subsets of synapses and intracellular organelles in neurons. J. Neurosci. 23, 5096–5104.

Prasad, T., and Weiner, J.A. (2011). Direct and indirect regulation of spinal cord Ia afferent terminal formation by the γ-Protocadherins. Front. Mol. Neurosci. 1–12.

Ranade, S.S., Woo, S.-H., Dubin, A.E., Moshourab, R.A., Wetzel, C., Petrus, M., Mathur, J., Bégay, V., Coste, B., Mainquist, J., et al. (2014). Piezo2 is the major transducer of mechanical forces for touch sensation in mice. Nature 516, 121–125.

Reed, C.B., Feltri, M.L., and Wilson, E.R. (2021). Peripheral glia diversity. J Anat 00, 1–16.

Ribeiro-da-Silva, A., Pignatelli, D., and Coimbra, A. (1985). Synaptic architecture of glomeruli in superficial dorsal horn of rat spinal cord, as shown in serial reconstructions. J. Neurocytol. 14, 203–220.

Robbins, E.M., Krupp, A.J., Perez de Arce, K., Ghosh, A.K., Fogel, A.I., Boucard, A., Südhof, T.C., Stein, V., and Biederer, T. (2010). SynCAM 1 Adhesion Dynamically Regulates Synapse Number and Impacts Plasticity and Learning. Neuron 68, 894–906.

Rossignol, S., Dubuc, R., and Gossard, J.-P. (2006). Dynamic sensorimotor interactions in locomotion. Physiol Rev 86, 89–154.

Rubinstein, R., Thu, C.A., Goodman, K.M., Wolcott, H.N., Bahna, F., Mannepalli, S., Ahlsen, G., Chevee, M., Halim, A., Clausen, H., et al. (2015). Molecular logic of neuronal self-recognition through protocadherin domain interactions. Cell 163, 629–642.

Rutlin, M., Ho, C.-Y., Abraira, V.E., Cassidy, C., Bai, L., Woodbury, C.J., and Ginty, D.D. (2014). The Cellular and Molecular Basis of Direction Selectivity of Aδ-LTMRs. Cell 159, 1640–1651.

Sano, K., Tanihara, H., Heimark, R.L., Obata, S., Davidson, M., St John, T., Taketani, S., and Suzuki, S. (1993). Protocadherins: a large family of cadherin-related molecules in central nervous system. The EMBO Journal 12, 2249–2256.

Sanz, E., Yang, L., Su, T., Morris, D.R., McKnight, G.S., and Amieux, P.S. (2009). Cell-type-specific isolation of ribosome-associated mRNA from complex tissues. Proc. Natl. Acad. Sci. U.S.a. 106, 13939–13944.

Schreiner, D., and Weiner, J.A. (2010). Combinatorial homophilic interaction between gamma-protocadherin multimers greatly expands the molecular diversity of cell adhesion. Proc. Natl. Acad. Sci. U.S.a. 107, 14893–14898.

Sharma, N., Flaherty, K., Lezgiyeva, K., Wagner, D.E., Klein, A.M., and Ginty, D.D. (2020). The emergence of transcriptional identity in somatosensory neurons. Nature 1–25.

Sieber, M.A., Storm, R., Martinez-de-la-Torre, M., Müller, T., Wende, H., Reuter, K., Vasyutina, E., and Birchmeier, C. (2007). Lbx1 acts as a selector gene in the fate determination of somatosensory and viscerosensory relay neurons in the hindbrain. J. Neurosci. 27, 4902–4909.

Suazo, I., Vega, J.A., García-Mesa, Y., García-Piqueras, J., García-Suárez, O., and Cobo, T. (2022). The Lamellar Cells of Vertebrate Meissner and Pacinian Corpuscles: Development, Characterization, and Functions. Front Neurosci 16, 790130.

Sugino, H., Hamada, S., Yasuda, R., Tuji, A., Matsuda, Y., Fujita, M., and Yagi, T. (2000). Genomic organization of the family of CNR cadherin genes in mice and humans. Genomics 63, 75–87.

Südhof, T.C. (2012). The presynaptic active zone. Neuron 75, 11–25.

Südhof, T.C. (2017). Synaptic Neurexin Complexes: A Molecular Code for the Logic of Neural Circuits. Cell 171, 745–769.

Südhof, T.C. (2018). Towards an Understanding of Synapse Formation. Neuron 100, 276–293.

Tasic, B., Nabholz, C.E., Baldwin, K.K., Kim, Y., Rueckert, E.H., Ribich, S.A., Cramer, P., Wu, Q., Axel, R., and Maniatis, T. (2002). Promoter choice determines splice site selection in protocadherin alpha and gamma pre-mRNA splicing. Molecular Cell 10, 21–33.

Thu, C.A., Chen, W.V., Rubinstein, R., Chevee, M., Wolcott, H.N., Felsovalyi, K.O., Tapia, J.C., Shapiro, L., Honig, B., and Maniatis, T. (2014). Single-cell identity generated by combinatorial homophilic interactions between α, β, and γ protocadherins. Cell 158, 1045–1059.

Togashi, H., Sakisaka, T., and Takai, Y. (2009). Cell adhesion molecules in the central nervous system. Cell Adh Migr 3, 29–35.

Usoskin, D., Furlan, A., Islam, S., Abdo, H., Lönnerberg, P., Lou, D., Hjerling-Leffler, J., Haeggström, J., Kharchenko, O., Kharchenko, P.V., et al. (2015). Unbiased classification of sensory neuron types by large-scale single-cell RNA sequencing. Nat Neurosci 18, 145–153.

Valtschanoff, J.G., Weinberg, R.J., and Rustioni, A. (1993). Amino acid immunoreactivity in corticospinal terminals. Exp Brain Res 93, 95–103.

Wang, X., Su, H., and Bradley, A. (2002a). Molecular mechanisms governing Pcdh-gamma gene expression: evidence for a multiple promoter and cis-alternative splicing model. Genes Dev. 16, 1890–1905.

Wang, X., Weiner, J.A., Levi, S., Craig, A.M., Bradley, A., and Sanes, J.R. (2002b). Gamma Protocadherins Are Required for Survival of Spinal Interneurons. Neuron 36, 843–854.

Weiner, J.A., and Jontes, J.D. (2013). Protocadherins, not prototypical: a complex tale of their interactions, expression, and functions. Front. Mol. Neurosci. 6, 4.

Weiner, J.A., Wang, X., Tapia, J.C., and Sanes, J.R. (2005). Gamma protocadherins are required for synaptic development in the spinal cord. Proc Natl Acad Sci USA 102, 8–14.

Woo, S.-H., Ranade, S., Weyer, A.D., Dubin, A.E., Baba, Y., Qiu, Z., Petrus, M., Miyamoto, T., Reddy, K., Lumpkin, E.A., et al. (2014). Piezo2 is required for Merkel-cell mechanotransduction. Nature 509, 622–626.

Wu, Q., and Maniatis, T. (1999). A striking organization of a large family of human neural cadherin-like cell adhesion genes. Cell 97, 779–790.

Zhang, Q., Lee, W.-C.A., Paul, D.L., and Ginty, D.D. (2019). Multiplexed peroxidase-based electron microscopy labeling enables simultaneous visualization of multiple cell types. Nat Neurosci 22, 828–839.

Zheng, Y., Liu, P., Bai, L., Trimmer, J.S., Bean, B.P., and Ginty, D.D. (2019). Deep Sequencing of Somatosensory Neurons Reveals Molecular Determinants of Intrinsic Physiological Properties. Neuron 103, 598–616.e7.

Zimmerman, A.L., Bai, L., and Ginty, D.D. (2014). The gentle touch receptors of mammalian skin. Science 346, 950–954.

Zimmerman, A.L., Kovatsis, E.M., Pozsgai, R.Y., Tasnim, A., Zhang, Q., and Ginty, D.D. (2019). Distinct Modes of Presynaptic Inhibition of Cutaneous Afferents and Their Functions in Behavior. Neuron 102, 420–434.e428.

