## Supplemental File for "γ-Protocadherins control synapse formation and peripheral branching of touch sensory neurons"

**KEY RESOURCES TABLE**

| REAGENT or RESOURCE | SOURCE | IDENTIFIER |
| --- | --- | --- |
| Antibodies | | |
| Guinea pig polyclonal anti-VGlut1 | Millipore Sigma | Cat# AB5905;  RRID: AB_2301751 |
| Rabbit polyclonal anti-Homer1 | Synaptic Systems | Cat# 160003 |
| Monoclonal Anti-HA antibody produced in mouse | Sigma-Aldrich | Cat# H3663; RRID:AB_262051 |
| Isolectin B4 647 | Invitrogen | Cat# I32450;  RRID: SCR_014365 |
| Chicken polyclonal anti Neurofilament (NFH) | Aves | Cat# NFH0211 |
| Rabbit polyclonal anti DsRed | Clontech | Cat# 632496;  RRID: AB_10013483 |
| Goat polyclonal anti mCherry | Sicgen | Cat# Ab0040-200; RRID: AB_2333092 |
| Goat polyclonal anti GFP-FITC | Abcam | Cat# ab6662;  RRID: AB_305635 |
| Mouse monoclonal anti-Neuronal Nuclei (NeuN) | Millipore | Cat# MAB377; RRID: AB_2298772 |
| Rat polyclonal anti-Troma1 (cytokeratin endo A) supernatant | U of Iowa (DSHB) | Cat# TROMA-I; RRID: AB_531826 |
| Rabbit anti GFP | Thermo Fisher Scientific | Cat# A-11122; RRID:AB_221569 |
| Rabbit anti S100 | Proteintech | Cat# 15146-1-AP; RRID:AB_2254244 |
| Mouse anti PSD-95 | Thermo Fisher Scientific | Cat# MA1-045; RRID:AB_325399 |
| Rabbit anti Human HA | Sigma-Aldrich | Cat# H6908; RRID:AB_260070 |
| Chicken anti MAP2 | Millipore | Cat# AB5543; RRID:AB_571049 |
| Guinea pig anti VGLUT2 | Millipore | Cat# AB2251; RRID:AB_1587626 |
| Goat anti-Rabbit IgG (H+L) Cross-Adsorbed Secondary Antibody, Alexa Fluor 488 | Thermo Fisher Scientific | Cat# A-11008; RRID:AB_143165 |
| Goat anti-Guinea Pig IgG (H+L) Highly Cross-Adsorbed Secondary Antibody, Alexa Fluor 488 | Thermo Fisher Scientific | Cat# A-11073; RRID:AB_2534117 |
| Goat anti-Guinea Pig IgG (H+L) Highly Cross-Adsorbed Secondary Antibody, Alexa Fluor 555 | Thermo Fisher Scientific | Cat# A-21435; RRID:AB_2535856 |
| Alexa Fluor 647-AffiniPure Donkey Anti-Chicken | Jackson ImmunoResearch Labs | Cat# 703-605-155; RRID:AB_2340379 |
| Donkey anti-Goat IgG (H+L) Cross-Adsorbed Secondary Antibody, Alexa Fluor 546 | Thermo Fisher Scientific | Cat# A-11056; RRID:AB_2534103 |
| Donkey anti-Rabbit IgG (H+L) Highly Cross-Adsorbed Secondary Antibody, Alexa Fluor 647 | Thermo Fisher Scientific | Cat# A-31573; RRID:AB_2536183 |
| Donkey anti-Rabbit IgG (H+L) Highly Cross-Adsorbed Secondary Antibody | Thermo Fisher Scientific | Cat# A-21206; RRID:AB_2535792 |
| Alexa Fluor 647-AffiniPure Donkey Anti-Guinea Pig IgG (H+L) | Jackson ImmunoResearch Labs | Cat# 706-605-148; RRID:AB_2340476 |
| Bacterial and Virus Strains | | |
| AAV2-retro-hSyn-synaptophysin-tdTomato | Penn Viral Core | Lehnert et al., 2021 |
| AAV-Retro-Flex-PLAP | Sara L. Prescott | Prescott et al., 2020 |
| Chemicals, Peptides, and Recombinant Proteins |  |  |
| Zamboni’s Fixative | Fisher Scientific | Cat# NC9335034 |
| Paraformaldehyde, reagent grade, crystalline | Millipore Sigma | Cat# P6148-500G |
| 4-nitro blue tetrazolium chloride | Sigma | Cat# 11383213001 |
| 5-bromo-4-chloro-3-indolyl-phosphate, 4-toluidine salt | Sigma | Cat# 11383221001 |
| Benzyl Benzoate | Sigma-Aldrich | Cat# B-6630 |
| Benzyl Alcohol | Sigma-Aldrich | Cat# 402834 |
| Tamoxifen | Sigma | Cat# T5648-1g |
| NAIR, commercial hair remover | Church and Dwight Co., Princeton NJ | ASIN# B001G7PTWU |
| Isofluorane | Henry Schein, supplied by Covetrus | Cat# 029405; |
| Sunflower seed oil | Sigma | Cat# S5007 |
| Tetrodotoxin citrate | Tocris | Cat# 1069 |
| Critical Commercial Assays |  |  |
| RNAscope Fluorescent Multiplex Assay | ACD Bio | Cat# 320850 |
| RNeasy Mini Kit | Qiagen | Cat# 74104 |
| Nextera XT DNA Library Prep Kit | illumina | Cat# FC-131 |
| SsoAdvanced Universal SYBR Green Supermix | Bio-Rad | Cat# 1725271 |
| Deposited Data | | |
| RNA-sequencing Data | This manuscript | GEO (https://www.ncbi.nlm.nih.gov/geo/) |
| Experimental Models: Cell Lines |  |  |
| HEK 293T cells | DSMZ | Cat# CRL-3216, RRID: CVCL_0063 |
| Experimental Models: Organisms/Strains | | |
| Mouse: *Advillin^Cre^* | Hasegawa et al., 2007 | RRID: IMSR_JAX:032536; MGI:1333798 |
| Mouse: *Advillin^CreER^* | Lau et al., 2011 | Strain #:026516  RRID:IMSR_JAX:026516 |
| Mouse: *Lbx1^Cre^* | Sieber et al., 2007 | N/A |
| Mouse: *Dhh^Cre^* | Jaegle et al., 2003 | Strain #:012929 RRID:IMSR_JAX:012929 |
| Mouse: *Cdx2^Cre^* | Coutaud and Pilon, 2013 | N/A |
| Mouse: *Piezo2^fl^* | JAX | 027720 |
| Mouse: *Pcdhg^fcon3^* | Prasad et al., 2008; Lefebvre et al., 2008 | Strain #:012644; RRID:IMSR_JAX:012644 |
| Mouse: *Rosa26^LSL-synaptophysin-tdTomato^* (Ai34) | Jackson Laboratory stock | Jax:012570 |
| Mouse: B6;129S6-Gt(ROSA)26Sor^tm14(CAG-tdTomato)Hze^/J (Ai14) | Jackson Laboratory stock; Zeng H. 2011 | Jax:007908 |
| Mouse: *Pcdhg^3R1^* | Garrett et al., 2019 | N/A |
| Mouse: *Pcdhg^3R2^* | Garrett et al., 2019 | N/A |
| Mouse: *Pcdhg^1R1^* | Garrett et al., 2019 | N/A |
| Mouse: *Pcdhgc3^KO^* | Garrett et al., 2019 | N/A |
| Mouse: *Pcdhg^TAKO^* | Chen et al., 2012 | N/A |
| Mouse: *TrkB^CreER^* | Rutlin et al., 2014 | MGI:5616440 |
| Mouse: *Ret^CreER^* | Luo et al., 2009 | N/A |
| Mouse: *PV^ires-Cre^* | Hippenmeyer et al., 2005 | MGI:3590684 |
| Mouse: *Rpl22^HA^* | Sanz et al., 2009 | Strain #:011029 RRID:IMSR_JAX:011029 |
| Oligonucleotides |  |  |
| Primers for mouse genotyping (see Table S1) | This paper | N/A |
| Mm-Pcdhga2 | ACD Bio | Cat# 2835811 |
| Mm-Pcdhga7-C2 | ACD Bio | Cat# 835781-C2 |
| Mm-Pcdhgb1-O1 | ACD Bio | Cat# 837621 |
| Mm-Pcdhgc3 | ACD Bio | Cat# 802841 |
| Mm-Pcdhgc4-C3 | ACD Bio | Cat# 835791 |
| Mm-Slc17a6-C3 | ACD Bio | Cat# 319171 |
| Mm-Nefh | ACD Bio | Cat# 443671 |
| Mm-Plp1 | ACD Bio | Cat# 428181 |
| Software and Algorithms |  |  |
| ImageJ | Schneider et al., 2012 | imagej.nih.gov/ij |
| Clampex 10 | Molecular Devices | https://www.moleculardevices.com; RRID: SCR_011323 |
| ZEN | Zeiss | https://www.zeiss.com/microscopy/us/downloads.html; RRID: SCR_013672 |
| R and RStudio | R and RStudio | https://www.r-project.org/ https://www.rstudio.com/ ; RRID: SCR_000432 |
| DESeq2 | Love et al., 2014 | https://bioconductor.org/packages/release/bioc/html/DESeq2.html ; RRID: SCR_015687 |
| ComplexHeatmap | Gu et al., 2016 | https://bioconductor.org/packages/release/bioc/html/ComplexHeatmap.html |
| MATLAB | Mathworks | https://www.mathworks.com/products/matlab.html; RRID: SCR_001622 |
| JRCLUST | Jun et al., 2017 | https://www.biorxiv.org/content/10.1101/101030v1 |

**Supplemental Figure Legends**

**Figure S1. Deep RNA sequencing of postnatal DRG neuron groups**

**(A)** Quantifications of the sizes of Homer1^+^ synapses surrounding sensory terminals labeled by synaptophysin-tdTomato at P0, P3, P10 and P21. Two-way ANOVA. ns, not significant; *p < 0.05; ***p < 0.001.

**(B)** Principal component analysis (PCA) of samples showing biological replicates for the same neuronal group largely cluster with each other. “Proproceptors” (labeled with *PV^Cre^*) and “Proprioceptors + Aβ SAI-LTMRs” (labeled with *TrkC^CreER^*) partially overlap, because proprioceptors are labeled in both groups. Expression levels are plotted using rlog count values.

**(C and D)** Heatmap depicting expression patterns of SUEGs for developing Aβ RA-LTMRs (C), Aδ-LTMRs, Aβ SAI-LTMRs and proprioceptors (D). Expression differences for each gene compared to the average expression levels for the same gene are plotted in the left heatmap. Average expression level of a gene is plotted in a second heatmap on the right of the main heatmap.

**(E)** *in situ* hybridization of 5 *Pcdhg* isoforms using P3 DRG sections (n = 3 animals), showing that *Pcdhgc3* is expressed throughout the DRG and *Pcdhga7* expression is higher in a subset of DRGs. The expression levels of *Pcdhga2*, *Pcdhgb1*, and *Pcdhgc4* are below detection, similar to the expression levels revealed by RNA sequencing.

**Figure S2. Loss of *Pcdhgs* in the sensory neurons do not affect adult body weight, startle response to air puff alone, acoustic PPI or EPM performance**

**(A)** Body weight of *Advillin^Cre^;Pcdhg^fcon3/fcon3^* mice is decreased at weaning age (P20-P22), but not significantly different from control mice at 6 weeks. Student’s unpaired t test.

**(B)** Response to a light air puff stimulus alone directed to the back hairy skin, showing that *Advillin^Cre^;Pcdhg^fcon3/fcon3^* mice exhibited normal level of movements in response to a light air puff stimulus. Student’s unpaired t test.

**(C)** aPPI with 250 ms ISI for littermate controls and *Advillin^Cre^;Pcdhg^fcon3/fcon3^* mice, showing no significant difference between genotypes. Student’s unpaired t test.

**(D)** Time in center of open field. Student’s unpaired t test.

**(E)** No different in the percent of time spent in the open arms of the EPM test between littermate controls and *Advillin^Cre^;Pcdhg^fcon3/fcon3^* mice. Student’s unpaired t test.

ns, not significant; *p < 0.05; ***p < 0.001.

**Figure S3. Normal mEPSC amplitude and overall sensory axonal projections in the *Advillin^Cre^;Pcdhg^fcon3/fcon3^* mice**

**(A)** Average mEPSC amplitudes in the control and *Advillin^Cre^;Pcdhg^fcon3/fcon3^;Ai14* mice. ns, not significant. Student’s unpaired t test.

**(B)** Representative images from *Advillin^Cre^;Ai14* and *Advillin^Cre^;Pcdhg^fcon3/fcon3^;Ai14* animals (n = 3 per genotype), showing the overall sensory axonal projection pattern in the dorsal horn in the mutants appear indistinguishable from controls.

**Figure S4. Pcdhgs are not required for sensory axon patterning around hair follicles**

**(A)** Representative images of back hairy skin from *Advillin^Cre^;Ai14* and *Advillin^Cre^;Pcdhg^fcon3/fcon3^;Ai14* mice. Lanceolate and circumferential sensory endings around hair follicles are visualized using tdTomato fluorescence (red), while terminal Schwann Cells are labeled using S100 immunostaining (green). Scale bar represents 20 μm.

**(B)** *Advillin^Cre^;Pcdhg^fcon3/fcon3^;Ai14* mice have normal length of lanceolate endings around guard hairs (n = 7 hair follicles for control and n= 13 for *Advillin^Cre^;Pcdhg^fcon3/fcon3^;Ai14*) and non-guard hairs (n = 45 hair follicles for control and n = 54 hair follicles for *Advillin^Cre^;Pcdhg^fcon3/fcon3^;Ai14*). Student’s unpaired t test.

**(C and D)** *Advillin^Cre^;Pcdhg^fcon3/fcon3^;Ai14* mice have normal length of terminal Schwann cell protrusions (C) and normal number of terminal Schwann cells (D) around guard hairs (n = 15 hair follicles for control and n= 18 for *Advillin^Cre^;Pcdhg^fcon3/fcon3^;Ai14*) and non-guard hairs (n = 45 hair follicles for control and n = 54 hair follicles for *Advillin^Cre^;Pcdhg^fcon3/fcon3^;Ai14*). Student’s unpaired t test.

**(E)** Number of Merkel cell per touch dome in the control and *Advillin^Cre^;Pcdhg^fcon3/fcon3^* mice. Student’s unpaired t test.

**(F)** Area of Meissner corpuscle in the control and *Advillin^Cre^;Pcdhg^fcon3/fcon3^* mice. Student’s unpaired t test.

**(G)** Representative spinal cord IHC image of a *Dhh^Cre^;Ai14* animal, showing that Tomato signal is only found in dorsal root, but not found in any cells in the spinal cord.

**(H-K)** Representative IHC images of lamina III from control and *Dhh^Cre^;Pcdhg^fcon3/fcon3^* mice (H), showing no difference observed in the vGluT1^+^ puncta density (I), Homer1^+^ puncta density (J), and the density of Homer1^+^ puncta surrounding vGluT1^+^ puncta (K).

ns, not significant.

**Figure S5. *Pcdhgs* function in the sensory neurons to maintain synapse and peripheral axonal branching**

**(A)** IHC images of dorsal root ganglion (DRG) showing the GFP fused-Pcdhg in, *Pcdhg^fcon3/fcon3^* (positive control), *Advillin^CreER^;Pcdhg^fcon3/fcon3^*, and wildtype (negative control) mice. IB4 labels small diameter non-peptidergic nociceptors. Scale bars represent 50 μm.

**(B)** Quantification showing the normalized level of GFP intensity in the *Advillin^CreER^;Pcdhg^fcon3/fcon3^* is reduced to wildtype level. one-way ANOVA test.

**(C)** IHC images of spinal cord lamina III from P21 littermate control (n = 4 animals) and *Advillin^CreER^;Pcdhg^fcon3/fcon3^* mice (n = 4 animals). All animals received the same tamoxifen treatment to control for potential neuronal toxicity from tamoxifen.

**(D-G)** Normalized densities of vGluT1^+^ puncta (D) and Homer1^+^ puncta (E). The normalized density and size of Homer1^+^ puncta around vGluT1^+^ puncta are quantified in (F) and (G), respectively. Student’s unpaired t test.

**(H)** Whole-mount immunostaining images of adult back hairy skin sections from control and *Advillin^Cre^;Pcdhg^fcon3/fcon3^* mice. Aβ field-LTMRs form circumferential endings around hair follicles and are NFH^+^ (white arrowheads). Aβ RA-LTMRs form lanceolate endings around guard hair follicles (noted as “G”) and awl/auchene hairs. TSCs are labeled using S100 immunostaining (green).

**(I)** Quantification of the percentage of non-guard hair follicles innervated by NFH^+^ Aβ field-LTMRs circumferential endings and Aβ RA-LTMRs lanceolate endings (n = 552 hair follicles from 4 control animals and n = 674 hair follicles from 4 *Advillin^CreER^;Pcdhg^fcon3/fcon3^* mice). Student’s unpaired t test.

**(J and K)** Example whole-mount immunostaining images of the Merkel cell complex (J), showing that the number major S100^+^ branches in *Advillin^Cre^;Pcdhg^fcon3/fcon3^* mice (n = 17 touch domes from 4 animals) is reduced compared to control (n = 37 touch domes from 4 animals) (K). Student’s unpaired t test.

ns, not significant; *p < 0.05; **p < 0.01; ***p < 0.001.

**Figure S6. Increased spinal cord neuron death in the *Lbx1^Cre^;Pcdhg^fcon3/fcon3^* mice and the localization of Pcdhgs in axons**

**(A-D)** IHC images of spinal cord from littermate control (n = 3 animals) and *Lbx1^Cre^;Pcdhg^fcon3/fcon3^* mice (n = 2 animals) (A), showing that the size of the dorsal horn (outlined with white dotted lines) is largely reduced in the *Lbx1^Cre^;Pcdhg^fcon3/fcon3^* mice (B). Normalized densities of vGluT1^+^ puncta (C) and Homer1^+^ puncta (D) in medial lamina III remain unchanged in the *Lbx1^Cre^;Pcdhg^fcon3/fcon3^* mice. Student’s unpaired t test.

**(E and F)** IHC images of spinal cord from littermate control (n = 5 animals) and *Lbx1^Cre^;Pcdhg^C3KO/fcon3^* mice (n = 3 animals) (E), showing that the size of the dorsal horn remains the same in the *Lbx1^Cre^;Pcdhg^C3KO/fcon3^* mice (F). Student’s unpaired t test.

**(G)** Representative spinal cord and skin images from P21 *Advillin^Cre^;cA1* animals. Pcdhga1-mCherry is found as puncta in the spinal cord, while Pcdhga1-mCherry is found in the sensory nerves in the hairy skin. Note that no Pcdhga1-mCherry signal is found in the sensory endings around hair follicles (not shown).

**(H)** Representative skin images from P21 *Advillin^Cre^;cC3* animals, showing that Pcdhgc3-mCherry is throughout the sensory axons, including the lanceolate and circumferential endings around hair follicles.

ns, not significant; ***p < 0.001.

**Figure S7. Decreased corticospinal innervation in lamina III of the *Cdx2^Cre^;Piezo2^f/f^* mutants.**

(**A**) Quantifications of the sustained phase neuronal responses in littermate control and *Advillin^Cre^;Pcdhg^fcon3/fcon3^* mice at various indentation forces. Two-way ANOVA.

(**B**) IHC images showing corticospinal terminals labeled with synaptophysin-tdTomato in the dorsal column, and IHC images showing corticospinal terminals colabeled with Homer1 in control and *Cdx2^Cre^;Piezo2^f/f^* mice. White dotted lines outline the shape of the dorsal column with labeling. Arrowheads point to the Tomato^+^ puncta with Homer1 labeling.

(**C and D**) Quantification showing the density of Tomato^+^ corticospinal terminals (C) and the number of Homer1^+^ excitatory synapses surrounding corticospinal terminals (D) in control and *Cdx2^Cre^;Piezo2^f/f^* mice. Each dot represents average number for an animal. Student’s unpaired t test.

**Table S1. Oligonucleotide List. Related to Key Resources Table.**

| **Mouse line** | **Oligo** | **Sequence (5’ to 3’)** |
| --- | --- | --- |
| *AdvillinCre* | transgene forward | CCC TGT TCA CTG TGA GTA GG |
|  | transgene reverse | AGT ATC TGG TAG GTG CTT CCA G |
|  | internal control | GCG ATC CCT GAA CAT GTC CAT C |
| *Advillin^CreER^* | AdvilCreERT2_1 | CCC TGT TCA CTG TGA GTA GG |
|  | AdvilCreERT2_2 | AGT ATC TGG TAG GTG CTT CCA G |
|  | AdvilCreERT2_3 | GCG ATC CCT GAA CAT GTC CAT C |
| *Ret^CreER^* | Ret forward | AGCGCAGGTCTCTCATCAGT |
|  | Ret WT reverse | GCAGGAGCAAAATCAGCTTC |
|  | RetCreERT2 reverse | GCGCGCCTGAAGATATAGAA |
| *TrkB^CreER^* | TrkB CreER-1 | GACACGCACTCCGACTGA |
|  | TrkB CreER-2 | ACACCTGCCTGATTCCTGAG |
|  | TrkB CreER-3 | TCCTCATCCTCTCCCACATC |
| *Ai34* | Rosa WT forward | AAGGGAGCTGCAGTGGAGTA |
|  | Rosa WT reverse | CCGAAAATCTGTGGGAAGTC |
|  | Ai34 forward | GGA GTG TGC CAA CAA GAC GGA GA |
|  | Ai34 reverse | CCA GCC TGT CTC CTT GAA CAC GA |
| *Pcdhg^fcon3^* | 5091-reverse | GACTAGTTGTCCCCAGGCATTGAAAAGAGG |
|  | 5090-forward | AGCAAGGTAGCTGGGCTGTTGGGGTGACCG |
|  | C3-GFP-reverse | CGGTGAACAGCTCCTCGCCCTTGCTCACCA |
|  | C3-GFP-forward | CAAGGTTGTCACTGACCCCATCTGACCATC |
| *ROSA26-CAG::lox-Stop-lox-Pcdhga1-mCherry* | Rosa-A1 Forward | AAA TGG ACT GAC TGG CCT GC |
|  | mCherry Reverse | CTC CCA CTT GAA GCC CTC GG |
| *ROSA26-CAG::lox-Stop-lox-Pcdhgc3-mCherry* | Rosa-C3 Forward | GTG AGC TCC CTG TAC CGA AC |
|  | mCherry Reverse (Common reverse for A1 and C3 forwards) | CTC CCA CTT GAA GCC CTC GG |
| *Pcdhg^1R1^* | PCDHGA1KF | TCTCTGGAGCTACTGCTGGA |
|  | PCDHGA1KR | AGCTCTTCCCGGTCTATCCT |
|  | PCDHGC3KR | TTACAGTGCAGGAGGGCAGCGT |
| *Pcdhg^3R1^* | PCDHGB2KF | AGACGCGTTAGGAAACTGGG |
|  | PCDHGB2KR | AATCCTTCTCTGCGCTGACC |
| *Pcdhg^3R2^* | PCDHGC4KF | AGAATTAGCGGATGGCAGCA |
|  | PCDHGC4KR | TTTGGTTCACCTCTCCAGCG |
| *Pcdhgc3^KO^* | KOCET forward | CCG GGA TGA GGC AGA GAC TGA A |
|  | KOCET reverse | ACT CCC ACC GTT CTC CAG G |
| *Ai14* | Rosa WT forward | AAGGGAGCTGCAGTGGAGTA |
|  | Rosa WT reverse | CCGAAAATCTGTGGGAAGTC |
|  | R26TOM-Mut_reverse | GGCATTAAAGCAGCGTATCC |
|  | R26TOM-Mut_forward | CTGTTCCTGTACGGCATGG |
| *Lbx1^Cre^* | LbxCre_Type_5'upper | CGCCTTCCTCTCGCACCGTC |
|  | LbxCre_Type_5'lower | GGCAGCCCGGACCGAC |
|  | LbxCre_Type_3'upper | GATGCGGTGGGCTCTATGGC |
|  | LbxCre_Type_3'lower | ACACTGCGTGTGGGCGACT |
| *Rpl22* | 9508 | GGG AGG CTT GCT GGA TAT G |
|  | 9509 | TTT CCA GAC ACA GGC TAA GTA CAC |
| Cre | JNTB84 | TGCCACGACCAAGTGACAGCAATG |
|  | JNTB85 | ACCAGAGACGGAAATCCATCGCTC |

**References**

Chen, W.V., Alvarez, F.J., Lefebvre, J.L., Friedman, B., Nwakeze, C., Geiman, E., Smith, C., Thu, C.A., Tapia, J.C., Tasic, B., et al. (2012). Functional Significance of Isoform Diversification in the Protocadherin Gamma Gene Cluster. Neuron *75*, 402–409.

Coutaud, B., and Pilon, N. (2013). Characterization of a novel transgenic mouse line expressing Cre recombinase under the control of the Cdx2 neural specific enhancer. Genesis *51*, 777–784.

Garrett, A.M., Bosch, P.J., Steffen, D.M., Fuller, L.C., Marcucci, C.G., Koch, A.A., Bais, P., Weiner, J.A., and Burgess, R.W. (2019). CRISPR/Cas9 interrogation of the mouse Pcdhg gene cluster reveals a crucial isoform-specific role for Pcdhgc4. PLoS Genet *15*, e1008554.

Gu, Z., Eils, R., and Schlesner, M. (2016). Complex heatmaps reveal patterns and correlations in multidimensional genomic data. Bioinformatics *32*, 2847–2849.

Hasegawa, H., Abbott, S., Han, B.-X., Qi, Y., and Wang, F. (2007). Analyzing somatosensory axon projections with the sensory neuron-specific Advillin gene. J. Neurosci. *27*, 14404–14414.

Hippenmeyer, S., Vrieseling, E., Sigrist, M., Portmann, T., Laengle, C., Ladle, D.R., and Arber, S. (2005). A developmental switch in the response of DRG neurons to ETS transcription factor signaling. PLoS Biol *3*, e159.

Jaegle, M., Ghazvini, M., Mandemakers, W., Piirsoo, M., Driegen, S., Levavasseur, F., Raghoenath, S., Grosveld, F., and Meijer, D. (2003). The POU proteins Brn-2 and Oct-6 share important functions in Schwann cell development. Genes Dev. *17*, 1380–1391.

Jun, J.J., Mitelut, C., Lai, C., Gratiy, S.L., Anastassiou, C.A., and Harris, T.D. (2017). Real-time spike sorting platform for high-density extracellular probes with ground-truth validation and drift correction. bioRxiv 101030.

Lau, J., Minett, M.S., Zhao, J., Dennehy, U., Wang, F., Wood, J.N., and Bogdanov, Y.D. (2011). Temporal control of gene deletion in sensory ganglia using a tamoxifen-inducible Advillin-Cre-ERT2 recombinase mouse. Mol Pain *7*, 100.

Lefebvre, J.L., Zhang, Y., Meister, M., Wang, X., and Sanes, J.R. (2008). gamma-Protocadherins regulate neuronal survival but are dispensable for circuit formation in retina. Development *135*, 4141–4151.

Lehnert, B.P., Santiago, C., Huey, E.L., Emanuel, A.J., Renauld, S., Africawala, N., Alkislar, I., Zheng, Y., Bai, L., Koutsioumpa, C., et al. (2021). Mechanoreceptor synapses in the brainstem shape the central representation of touch. Cell *184*, 5608–5621.e5618.

Love, M.I., Huber, W., and Anders, S. (2014). Moderated estimation of fold change and dispersion for RNA-seq data with DESeq2. Genome Biol. *15*, 550–21.

Luo, W., Enomoto, H., Rice, F.L., Milbrandt, J., and Ginty, D.D. (2009). Molecular identification of rapidly adapting mechanoreceptors and their developmental dependence on ret signaling. Neuron *64*, 841–856.

Prasad, T., Wang, X., Gray, P.A., and Weiner, J.A. (2008). A differential developmental pattern of spinal interneuron apoptosis during synaptogenesis: insights from genetic analyses of the protocadherin-gamma gene cluster. Development *135*, 4153–4164.

Prescott, S.L., Umans, B.D., Williams, E.K., Brust, R.D., and Liberles, S.D. (2020). An Airway Protection Program Revealed by Sweeping Genetic Control of Vagal Afferents. Cell *181*, 574–589.e14.

Rutlin, M., Ho, C.-Y., Abraira, V.E., Cassidy, C., Bai, L., Woodbury, C.J., and Ginty, D.D. (2014). The Cellular and Molecular Basis of Direction Selectivity of Aδ-LTMRs. Cell *159*, 1640–1651.

Sanz, E., Yang, L., Su, T., Morris, D.R., McKnight, G.S., and Amieux, P.S. (2009). Cell-type-specific isolation of ribosome-associated mRNA from complex tissues. Proc. Natl. Acad. Sci. U.S.a. *106*, 13939–13944.

Schneider, C.A., Rasband, W.S., and Eliceiri, K.W. (2012). NIH Image to ImageJ: 25 years of image analysis. Nat Methods *9*, 671–675.

Sieber, M.A., Storm, R., Martinez-de-la-Torre, M., Müller, T., Wende, H., Reuter, K., Vasyutina, E., and Birchmeier, C. (2007). Lbx1 acts as a selector gene in the fate determination of somatosensory and viscerosensory relay neurons in the hindbrain. J. Neurosci. *27*, 4902–4909.
